# REBEL, Reproducible Environment Builder for Explicit Library resolution

**DOI:** 10.64898/2026.04.04.716498

**Authors:** Eliseo Martelli, Maria Luisa Ratto, Beatrice Nuvolari, Maddalena Arigoni, Jianli Tao, Francesco Maria Antonio Micocci, Luca Alessandri

**Affiliations:** University of Turin; university of Turin; Boston Children's Hospital, Harvard Medical School

**Keywords:** reproducibility, bioinformatics, dependency resolution, Docker, FAIR, software environments, package management

## Abstract

**Background:** Achieving FAIR-compliant computational research in bioinformatics is systematically undermined by two compounding challenges that existing tools leave unresolved: long-term reproducibility and accessibility. Standard package managers re-download dependencies from live repositories at every build, making environments vulnerable to library disappearance and version drift, and pinning a package version does not pin the versions of its transitive dependencies, causing divergences between builds performed at different points in time. Compounding this, packages from repositories such as CRAN, Bioconductor, and PyPI frequently omit critical system-level dependencies from their installation metadata, leaving users to manually discover which underlying library is missing or which version is required. Beyond these technical failures, constructing a truly reproducible environment demands expertise in containerization making reproducibility in practice a privilege and not a standard.

**Findings:** We present REBEL (Reproducible Environment Builder for Explicit Library Resolution), a framework that addresses both challenges through three dependency inference heuristics: (i) Deep Inspection of source code, (ii) Fuzzy Matching against a manually curated knowledge base, and (iii) Conservative Dependency Locking. The resolved dependency stack is then archived into a self-contained local store, enabling offline and deterministic rebuilds at any future time. We compared the installation of 1,000 randomly sampled CRAN packages in isolated Docker containers versus the standard package manager and REBEL resolved 149 of 328 standard installation failures (45.4%). Moreover through its DockerBuilder component, REBEL further generates fully reproducible Docker images from a plain text requirements file, making deterministic environment construction accessible without expertise in containerization.

**Conclusions:** REBEL provides a practical foundation for FAIR-compliant, long-term reproducible bioinformatics analyses, making deterministic environment construction accessible to researchers regardless of their technical background.

REBEL is freely available at https://github.com/Rebel-Project-Core

## Findings

### Background and Motivation

Reproducibility is a cornerstone of scientific progress[1,2], yet a large proportion of published bioinformatics analyses cannot be independently verified or reused. A 2016 survey of 1,500 scientists found that more than 70% had failed to reproduce another researcher’s results [3], and computational biology is no exception. The root cause is rarely fraud or error in the analysis itself, but rather the inability to reconstruct the software environment in which the analysis was originally executed. Indeed, modern bioinformatics workflows depend on complex software stacks spanning multiple programming languages, package repositories, and system-level libraries [4], though, long-term reproducibility in this setting is critically undermined by a structural flaw in how package managers handle versioning.

Pinning the version of a package does not pin the versions of its dependencies: standard package managers install the newest available versions of all transitive dependencies rather than those valid at the time of the original release. In practice, widely used packages can carry dozens or even hundreds of transitive dependencies, each of which is free to change independently. A package such as Seurat[4–6], for instance, depends on a large ecosystem of R and system libraries whose versions evolve continuously: Seurat alone requires 433 (Supplementary Table S1,S2) transitive dependencies across CRAN and system package managers, meaning that two installations of the same version performed months apart may produce different environments, different numerical results, or outright build failures.

Existing tools only partially address this problem: Conda [7]and virtual environments provide isolation but re-download dependencies from live repositories at every build, making them vulnerable to library disappearance, package renaming, and API changes over time, while container systems such as Docker [7,8] and Singularity [9] can preserve a functioning environment as a static image but cannot reconstruct it deterministically from scratch, as they capture a snapshot rather than a reproducible recipe.

Accessibility represents an equally important barrier, as many packages from repositories such as CRAN, Bioconductor, or PyPI omit critical system-level dependencies from their installation metadata, so when an installation fails, users are left digging into a rabbit hole of error logs with no systematic guidance on which underlying library is missing or which version is required. Building a reproducible Docker environment under these conditions requires expertise in Dockerfiles, system package managers, and the often non-obvious relationship between high-level package names and their system-level requirements, a combination of skills that effectively restricts reproducibility tools to computationally sophisticated labs and excludes the majority of the bioinformatics community.

Our previous framework, CREDO [9,10], introduced the concept of capturing installation steps to enable repeatable reconstructions, but it assumed that installations would succeed and did not address the underlying unreliability of dependency resolution. Earlier efforts such as rCASC [11] and the Reproducible Bioinformatics Project [11,12] further highlighted the practical impact of incomplete environment specification on real-world pipelines, yet the persistent failure of dependency-unsafe packages and the inability to reconstruct historical dependency sets remained unresolved.

To address both problems simultaneously, we introduce REBEL (Reproducible Environment Builder for Explicit Library Resolution), a framework that operates at the dependency resolution layer itself rather than assuming successful installation, providing the stable execution substrate on which reproducibility tools rely and making deterministic environment construction both technically achievable and genuinely accessible to researchers regardless of their computational background.

### Comparison with Existing Tools

A number of tools have been developed to address reproducibility in computational research, yet each leaves at least one of the two structural problems identified above, namely unreliable dependency resolution and the inability to reconstruct environments deterministically without network access, substantially unresolved. Table 1 summarizes the key functional differences between the most widely adopted solutions and REBEL across the dimensions most relevant to long-term reproducibility and accessibility.

**Table 1.**
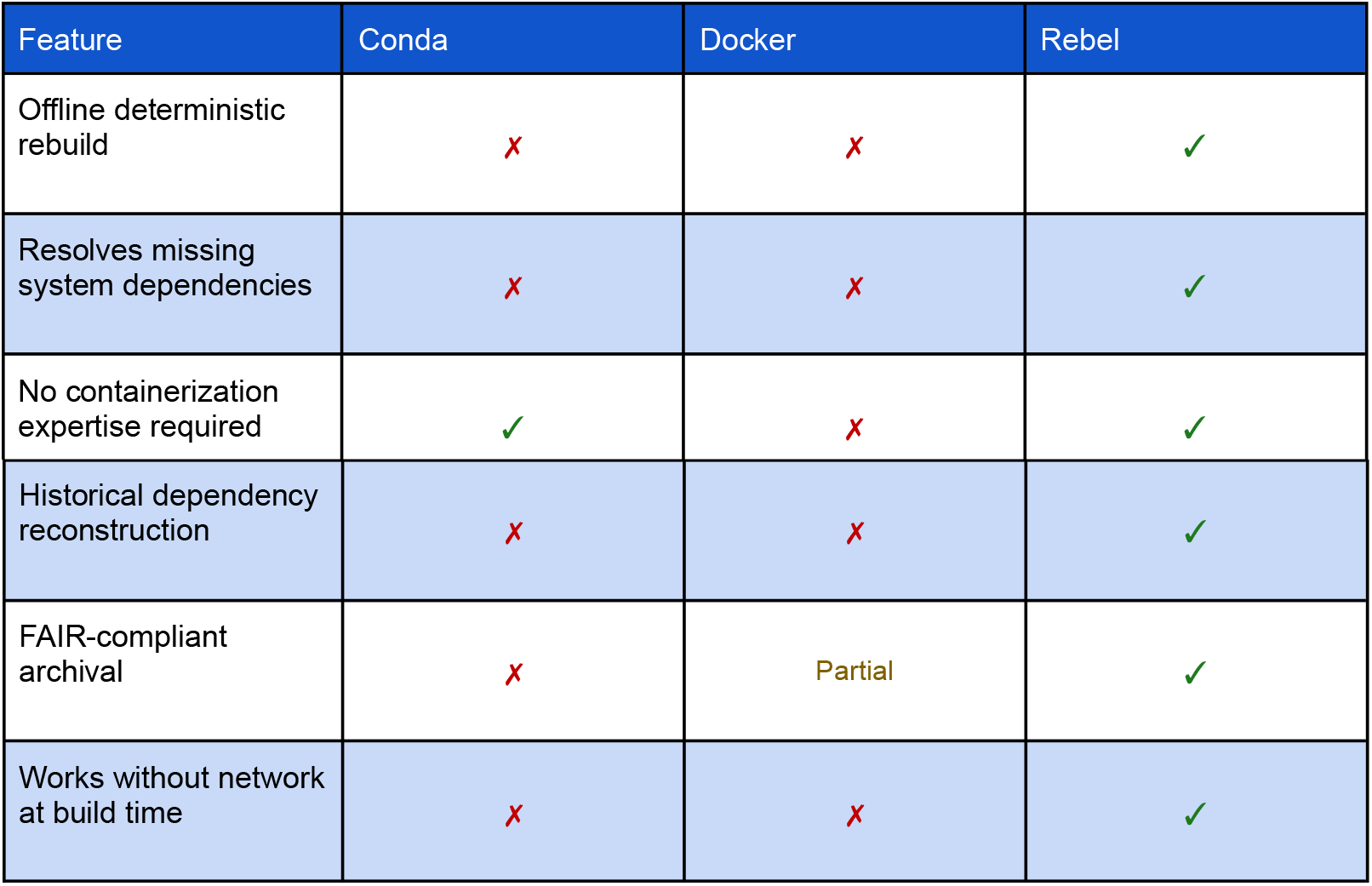
Comparison of REBEL with existing environment management tools.

Conda provides environment isolation and does not require expertise in containerization, making it accessible to a broad user base, but in its standard configuration it re-downloads all dependencies from live repositories at every build, meaning that the resulting environment is only reproducible as long as the upstream packages remain available at the same versions and the repository indices remain unchanged, a condition that is routinely violated over timescales relevant to scientific publication. Docker can preserve a functioning environment either as a static image or as a Dockerfile that specifies the build recipe, but neither approach solves the underlying problem: a static image is immediately executable but cannot be reconstructed from scratch, while a Dockerfile that pulls packages from live repositories at build time is subject to exactly the same version drift and dependency disappearance that affects Conda, since every rebuild fetches whatever versions happen to be currently available in the upstream repositories rather than those that were valid at the time of the original build. None of these tools perform any form of historical dependency reconstruction, meaning that installing a package at a specific version will silently pull in the newest available versions of all transitive dependencies rather than those that were valid at the time of the original release, exposing users to the version drift and silent numerical inconsistencies described in the previous section.

### Implementation

REBEL organizes the environment reconstruction process into three distinct phases: Discovery, Save, and Apply (Fig. 1A). When a user requests a package via rebel install, the framework does not blindly trust the declared installation metadata but instead applies three core heuristics to actively resolve the full dependency graph before attempting any installation.

**Figure 1.**
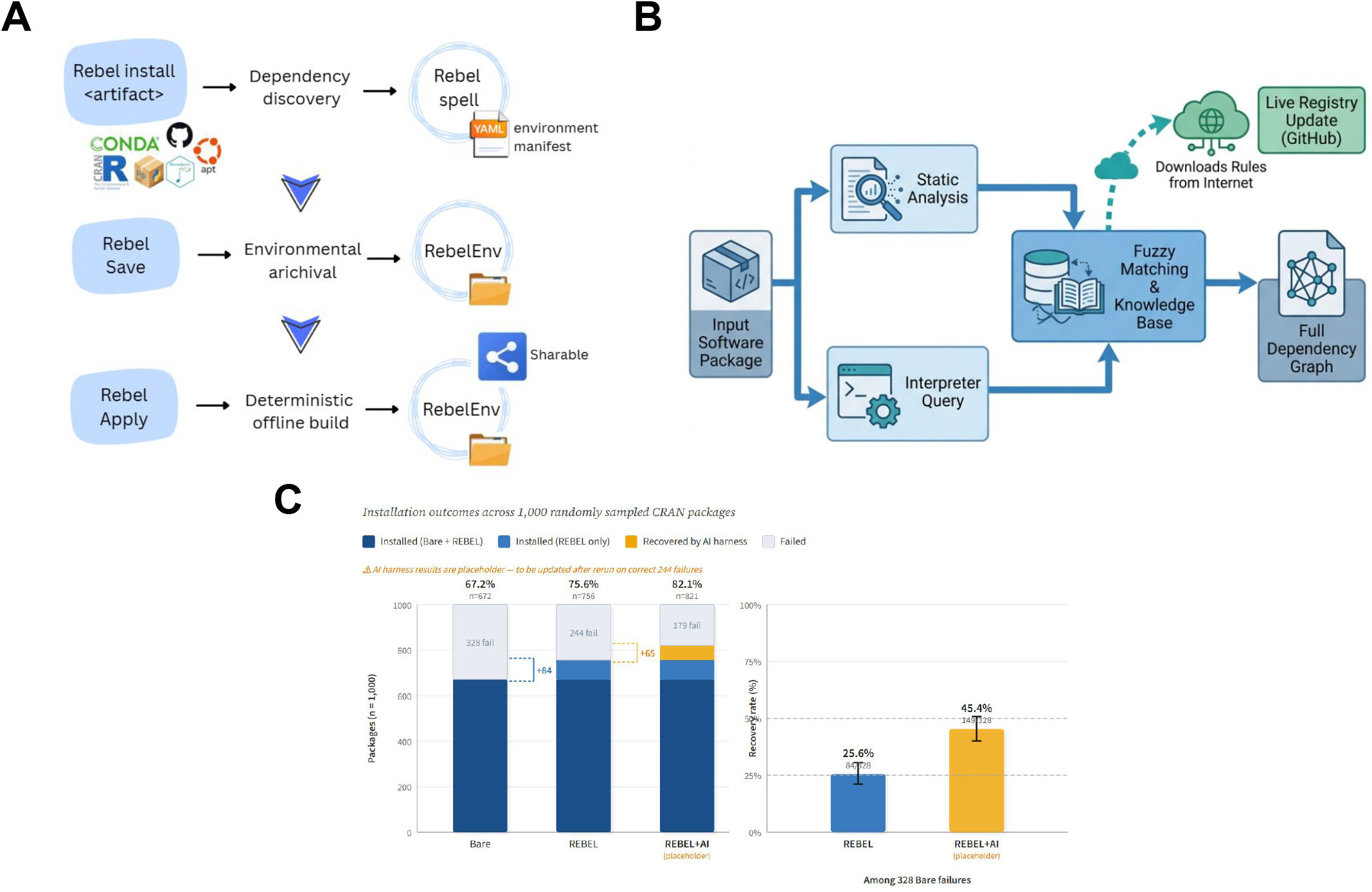
Architecture, Heuristics and Performance of the REBEL Framework. (A) Workflow Overview. REBEL organizes reproducibility into three distinct phases: Discovery, where dependencies are modeled into a persistent manifest (rebelspell.yaml); Save, where all artifacts including interpreters and system libraries are downloaded into a local archive (rebelenv/); and Apply, which performs deterministic, offline-capable environment reconstruction. (B) Dependency Resolution Heuristics. To resolve incomplete or implicit metadata, REBEL employs a multi-strategy discovery pipeline. Static Analysis scans source code for undeclared system requirements, while Interpreter Version Retracing retraces runtime-compatible dependency versions. These streams converge on a Knowledge Base module that employs fuzzy matching to resolve ambiguity. This module features a Live Registry Update mechanism (green), allowing the system to fetch new resolution rules from GitHub at runtime without requiring software updates. C — Empirical Validation. Installation outcomes for 1,000 randomly sampled CRAN packages. Left panel: stacked outcome breakdown for standard BiocManager installation (Bare) and REBEL. Of the 328 packages that failed under Bare installation, REBEL successfully resolved 84 (25.6%), recovering one in four failures through its dependency inference heuristics. The remaining 244 packages failed under both conditions. Right panel: zoom on the 328 Bare failures, showing REBEL’s recovery rate. All REBEL-installed packages were verified to be deterministically rebuildable across machines. Error bars represent 95% Wilson confidence intervals on the recovery rate.

The first heuristic, Deep Inspection, performs a static analysis of the package source code to uncover hidden system dependencies that authors have omitted from the official metadata identifying them proactively by scanning the source directly, allowing the full set of requirements to be known before installation begins.

The second heuristic, Fuzzy Matching, addresses the translation problem between high-level package names and their underlying system-level equivalents. Since the name used to import a package often differs from the name of the corresponding system package, REBEL employs a fuzzy-matching engine backed by a dynamic Knowledge Base (Fig. 1B). This knowledge base handles common naming patterns via regular expressions and resolves complex edge cases through a manually curated database of verified resolution rules that can be updated from GitHub at runtime, without requiring any software update to REBEL itself.

The third heuristic, Conservative Dependency Locking, addresses version instability in the dependency graph. A critical flaw in standard package managers is that when a dependency lacks an explicit version constraint, the manager defaults to installing the newest available release, which may be incompatible with the user’s interpreter or existing environment. REBEL resolves this by enforcing a backward-iterative version resolution strategy: it iteratively attempts to install candidate versions of each dependency, from the most recent backward, until one compiles and loads successfully in the target environment. In practice, this mechanism prevents the kind of breakage that commonly affects packages such as Seurat, where the dependency Matrix is frequently updated to a version incompatible with the system R interpreter, causing installation failures that users would otherwise have to diagnose manually from verbose build logs.

For packages that remain unresolved after all three heuristics have been applied, REBEL provides an AI-driven harness that extends coverage by continuously expanding the Knowledge Base underpinning the Fuzzy Matching engine. The harness extracts the most informative fragments from installation logs via TF-IDF ranking, which filters out low-signal noise and retains only the error messages most likely to indicate a missing or misidentified dependency, and submits them to a large language model that generates candidate resolution rules. These candidate rules are reviewed and, when validated, integrated into the manually curated knowledge base, expanding its coverage without requiring manual log inspection by the development team. The harness is designed to run cyclically on the full set of failed packages across CRAN, Bioconductor, and PyPI, progressively reducing the pool of unresolvable cases over time as new rules accumulate and the knowledge base matures.

Once the full dependency graph has been resolved, the final environment state is recorded in a rebelspell.yaml manifest that explicitly specifies every required component and its validated version. rebel save then downloads all of these components, including packages, system libraries, and binary assets, into a local RebelEnv archive. Finally, rebel apply reconstructs the environment using exclusively these archived local assets, with no network access required, ensuring that the workflow can be rebuilt deterministically at any future time and on any machine with the same architecture.

For maximum reproducibility, particularly in the context of scientific publications, we recommend deploying REBEL within Docker containers using a fixed base image (ubuntu:24.04), which eliminates residual variability from OS-level differences in kernel versions, system library implementations, or compiler toolchains that may persist even with fully locked dependency graphs. To make this recommended workflow accessible without requiring Docker expertise, REBEL includes DockerBuilder, an automated tool that transforms a simple requirements file into a fully reproducible Docker image without any knowledge of Dockerfile syntax or container internals. Users populate a plain text file with the packages they need, specifying the source repository for each entry (e.g., cran Seurat, pip pandas, github owner/repo), and execute a single script available for both Linux and macOS. DockerBuilder then handles the entire process in two automated stages. In the first stage, a temporary Docker container resolves all dependencies via rebel install and archives every required component via rebel save into a complete offline bundle. In the second stage, a new clean Dockerfile is generated that copies this pre-built archive and reconstructs the environment via rebel apply using only local files, with no network access at build time. Because the second stage relies exclusively on archived assets, the resulting image is fully reproducible: rebuilding it at any future time, on any machine, will produce an identical environment. The final image can be executed with standard Docker commands, distributed via public container registries such as Docker Hub or GitHub Container Registry, or archived alongside a scientific publication as a permanent and self-contained record of the computational environment.

## Results

To evaluate REBEL under rigorous and reproducible conditions, we developed a custom test-harness that performs large-scale installation benchmarks in fully isolated environments [https://github.com/Rebel-Project-Core/Test_harness2.0]. We randomly sampled 1,000 R packages compatible with the R version installed in our Docker container (ubuntu:24.04) and tested each one in a pristine container that was destroyed and recreated for every individual attempt, ensuring that no residual state from previous installations could influence subsequent results. For the baseline condition, referred to throughout as Bare, we used the standard BiocManager::install command without any additional dependency resolution. For the REBEL condition, we applied the complete Discovery, Save, and Apply workflow and validated success by verifying that each package could be loaded correctly.

Of the 1,000 packages tested, 672 (67.2%) succeeded under Bare installation, while 756 (75.6%) succeeded under REBEL (Fig. 1C). These global figures should however be interpreted in light of the composition of the sample: the majority of CRAN packages are well-maintained and installed correctly under both conditions, meaning that a random sample is by construction dominated by packages that present no challenge to either approach. The practically relevant metric is therefore the recovery rate among the 328 packages that failed under Bare installation, which represents the population of packages that a researcher would actually turn to REBEL to resolve. Through its three heuristics, REBEL resolved 84 of these failures (25.6%) without requiring any manual intervention.

Crucially, every successful REBEL installation was verified to be deterministically rebuildable: rebuilding the archived environment across different machines with the same architecture consistently produced identical environments, confirming that the offline archive is a fully self-contained and stable record of the computational context.

The AI-driven harness was then applied to the 244 packages that remained unresolved after REBEL’s three heuristics. The harness analyzed each installation log via TF-IDF ranking to extract the most informative error fragments and used them to identify the missing system dependencies. Once validated and integrated into the knowledge base, these mappings resolved an additional 65 packages (26.6% of the previously unresolved cases). In total, REBEL recovered 149 of the 328 original Bare failures, corresponding to a cumulative recovery rate of 45.4%.. A full cyclical run of the AI harness across all failed packages in CRAN, Bioconductor, and PyPI is currently ongoing, and we expect the knowledge base to continue expanding as new resolution rules are validated and integrated.

## Conclusions

Reproducible computational science in bioinformatics faces two structural problems that existing tools leave unresolved: the fragility of long-term reproducibility, as standard package managers depend on live repositories that drift and disappear over time, and the inaccessibility of environment construction, which requires expertise in containerization and system package management that the majority of researchers do not possess. REBEL addresses both directly. To solve long-term reproducibility, it identifies every component that a package needs to run, down to the lowest-level system library, archives them locally, and uses exclusively those archived assets to reconstruct the environment at any future time, guaranteeing an identical result regardless of when or where the build is performed. To solve accessibility, DockerBuilder automates this entire process: the researcher provides a plain text file listing the software they need, and the tool produces a fully reproducible Docker image without requiring any expertise in containers or package management.

The practical consequence is a fundamental change in when reproducibility enters the research workflow. Rather than attempting to reconstruct an environment after analysis is complete, researchers can work inside a fully reproducible container from the first day of a project, producing a permanent and self-contained record of the computational environment that can be shared via public registries, archived alongside publications, and rebuilt indefinitely without dependence on upstream repositories.

REBEL is fully open-source and community-driven, and its AI-driven expansion system ensures that the knowledge base grows continuously as new failure patterns emerge, so that the tool remains effective as the bioinformatics software ecosystem evolves. Together, these properties provide the practical foundation that genuinely FAIR-compliant computational research requires.

## Methods

### Context Optimization for AI-Driven Log Analysis

A core design challenge in the AI-driven harness is that installation logs produced by complex build systems can span hundreds of kilobytes, containing thousands of lines of successful dependency resolutions, compiler warnings, and progress indicators that carry no diagnostic value. While modern large language models support large context windows, submitting raw logs in their entirety is counterproductive for two reasons. First, the signal-to-noise ratio is extremely low, as the informative content, typically a handful of error messages indicating a missing library or an incompatible version, is buried within a large volume of irrelevant output. Second, research has demonstrated that LLM performance follows a U-shaped curve with respect to information placement in the input context: models retrieve and use information located at the beginning or end of the input significantly more reliably than information positioned in the middle, a phenomenon known as the “Lost in the Middle” effect [13]. When the diagnostic signal is embedded in the middle of a large log, model reasoning degrades accordingly.

To address this, we implemented a preprocessing pipeline that reduces each installation log to its most informative fragments before submission to the language model. The log is first segmented into overlapping 10-line chunks, which preserves local context around each error event. Each chunk is then represented as a TF-IDF vector, where Term Frequency weights terms that appear frequently within a given chunk and Inverse Document Frequency downweights terms that appear across many chunks and therefore carry little discriminative value. Common log noise such as “building”, “at”, or “installing” receives low IDF weight, while rare and diagnostically relevant terms such as “SegmentationFault”, “undefined reference”, or “package not found” receive high weight. Chunks are then ranked by cosine similarity against a fixed query (“error failure exception traceback”), chosen for its linguistic universality across build systems including C, C++, R, Python, and Fortran, and the top K chunks are selected for submission to the language model.

The optimal value of K was determined empirically by analyzing the distribution of relevance scores across 245 failure logs (Fig. 2). The 90th percentile of chunk relevance scores shows a sharp decay after rank 10, crossing the 0.05 relevance threshold, while the cumulative utility curve, measured as the sum of mean relevance scores, flattens substantially beyond K=10, indicating diminishing returns from additional chunks. A relevance heatmap across all 245 logs confirms that the vast majority of high-relevance signal is captured within the first 10 ranked chunks. Based on this analysis, K=10 was selected as the operating parameter for all experiments reported in this paper.

**Figure 2.**
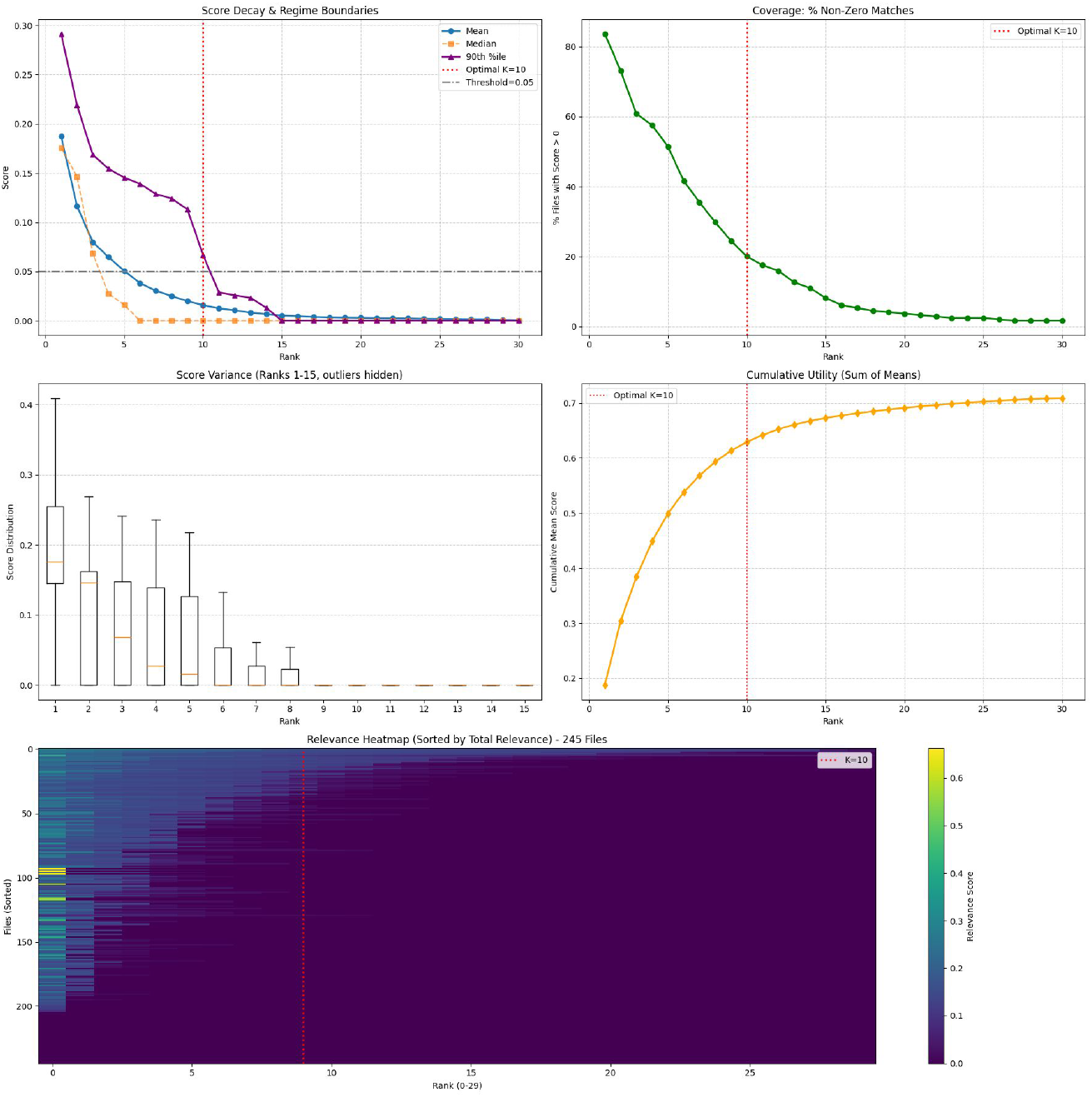
Empirical optimization of the TF-IDF chunk selection parameter K across 245 failure logs. (A) Score decay showing mean, median, and 90th percentile of chunk relevance scores by rank; the 90th percentile crosses the 0.05 threshold at K=10 (red dotted line). (B) Coverage: percentage of logs with at least one non-zero relevance score at each rank. (C) Score variance across ranks 1-15, showing near-zero distributions beyond rank 8. (D) Cumulative utility (sum of mean scores), flattening beyond K=10. Together, these analyses identify K=10 as the optimal cutoff beyond which additional chunks contribute more noise than signal.

### Package Sampling Strategy

The set of candidate packages was first filtered for compatibility with the R version installed in the test container (ubuntu:24.04) by querying the CRAN repository via available.packages() and excluding any package declaring a minimum R version requirement exceeding that of the container, using the script generate_cran_list.R [https://github.com/Rebel-Project-Core/Test_harness2.0 ]. From the resulting list of compatible packages, a single batch of 1,000 packages was drawn by uniform random sampling without replacement using sampleBatches.sh. The exact list of sampled packages is provided in cran_packages_sampled.txt in the test-harness repository, ensuring that all results reported in this paper can be independently reproduced on the identical set of packages. The sampled list was fixed prior to any testing and used identically for both the Bare and REBEL conditions.

## Supporting information

Harness Test

Seurat Network HTML

Seurat Dependencies

## References

1. Wilkinson MD, Dumontier M, Aalbersberg IJJ, Appleton G, Axton M, Baak A,et al. The FAIR Guiding Principles for scientific data management and stewardship. Sci Data. 3:1600182016;

2. Ioannidis JPA. Why most published research findings are false. PLoS Med. 2:e1242005;

3. Baker M. 1,500 scientists lift the lid on reproducibility. Nature. 533:452–42016;

4. Mangul S, Mosqueiro T, Abdill RJ, Duong D, Mitchell K, Sarwal V, et al. Challenges and recommendations to improve the installability and archival stability of omics computational tools. PLoS Biol. 17:e30003332019;

5. Hao Y, Stuart T, Kowalski MH, Choudhary S, Hoffman P, Hartman A, et al. Dictionary learning for integrative, multimodal and scalable single-cell analysis. Nat Biotechnol. 42:293–3042024;

6. Hao Y, Hao S, Andersen-Nissen E, Mauck WM 3rd, Zheng S, Butler A, et al. Integrated analysis of multimodal single-cell data. Cell. 184:3573–87.e292021;

7. Grüning B, Dale R, Sjödin A, Chapman BA, Rowe J, Tomkins-Tinch CH, et al. Bioconda: sustainable and comprehensive software distribution for the life sciences. Nat Methods. 15:475–62018;

8. Merkel D. Docker: lightweight Linux containers for consistent development and deployment. Linux Journal. 2014:22014;

9. Sochat VV, Prybol CJ, Kurtzer GM. Enhancing reproducibility in scientific computing: Metrics and registry for Singularity containers. PLoS One. 12:e01885112017;

10. Alessandri S, Ratto ML, Rabellino S, Piacenti G, Contaldo SG, Pernice S, et al. CREDO: a friendly Customizable, REproducible, DOcker file generator for bioinformatics applications. BMC Bioinformatics. Springer Science and Business Media LLC; 25:1102024;

11. Alessandrì L, Cordero F, Beccuti M, Arigoni M, Olivero M, Romano G, et al. rCASC:reproducible classification analysis of single-cell sequencing data. Gigascience. Oxford University Press (OUP); 2019; doi:10.1093/gigascience/giz105.

12. Kulkarni N, Alessandrì L, Panero R, Arigoni M, Olivero M, Ferrero G, et al. Reproducible bioinformatics project: a community for reproducible bioinformatics analysis pipelines. BMC Bioinformatics. Springer Science and Business Media LLC; 19:3492018;

13. Liu NF, Lin K, Hewitt J, Paranjape A, Bevilacqua M, Petroni F, et al. Lost in the Middle: How Language Models Use Long Contexts.

