## Supplementary material for "REBEL, Reproducible Environment Builder for Explicit Library resolution": Seurat Network HTML: SupplementaryTable2S.html

Seurat — Dependency Network


### 🧬 Seurat — Dependency Network

Seurat (root)

CRAN/BiocManager: 139

APT: 292

⏸ Pausa
🔭 Fit
⭐ Seurat
APT only
CRAN/Bio only
Tutti

Calcolo layout con 431 nodi e 1350 archi…

**📦 431 pacchetti** · **1350 dipendenze**  
Scroll → zoom · Drag → sposta · Click → dettagli

✕
