## Supplementary material for "REBEL, Reproducible Environment Builder for Explicit Library resolution": Seurat Dependencies: SupplementaryTable1S.yaml

```
# !!! WARNING !!!
# This file is automatically generated by CREDO.
# CREDO is a tool designed to create reproducible environments.
---
apt:
    - name: r-base
      dependencies:
        - name: r-api-4.0
          optional: true
        - name: liblapack.so.3
          optional: true
        - name: libhogweed6
          optional: true
        - name: libgnutls30
          optional: true
        - name: libblas.so.3
          optional: true
        - name: libglib2.0-0
          optional: true
        - name: libreadline8
          optional: true
        - name: libnettle8
          optional: true
        - name: debconf-2.0
          optional: true
        - name: x11-common
        - name: xfonts-encodings
        - name: zlib1g
        - name: libc6
        - name: libfontenc1
        - name: libdb5.3t64
        - name: fonts-noto-mono
        - name: libfreetype6
        - name: xfonts-utils
        - name: fonts-dejavu-mono
        - name: libacl1
        - name: sysvinit-utils
        - name: lsb-base
        - name: libsasl2-modules-db
        - name: libtasn1-6
        - name: libffi8
        - name: libp11-kit0
        - name: libkeyutils1
        - name: fonts-noto-core
        - name: fonts-texgyre
        - name: fonts-urw-base35
        - name: fonts-freefont-ttf
        - name: fonts-freefont-otf
        - name: fonts-croscore
        - name: fonts-liberation
        - name: fonts-dejavu-core
        - name: libselinux1
        - name: tar
        - name: libmd0
        - name: libtextwrap1
        - name: libslang2
        - name: libnewt0.52
        - name: libdebian-installer4
        - name: libuuid1
        - name: libbsd0
        - name: libsharpyuv0
        - name: libdatrie1
        - name: libthai-data
        - name: libgraphite2-3
        - name: libblkid1
        - name: libnettle8t64
        - name: libgmp10
        - name: libhogweed6t64
        - name: libssl3t64
        - name: libsasl2-2
        - name: libunistring5
        - name: libidn2-0
        - name: libgnutls30t64
        - name: libkrb5support0
        - name: libk5crypto3
        - name: libcom-err2
        - name: libkrb5-3
        - name: libxdmcp6
        - name: libxau6
        - name: fontconfig-config
        - name: libexpat1
        - name: libzstd1
        - name: liblzma5
        - name: libbz2-1.0
        - name: dpkg
        - name: openssl
        - name: sensible-utils
        - name: debconf
        - name: libreadline8t64
        - name: cdebconf
        - name: libice6
        - name: libsm6
        - name: libx11-data
        - name: libxext6
        - name: libx11-6
        - name: libxss1
        - name: libxrender1
        - name: libfontconfig1
        - name: libxft2
        - name: libtirpc-common
        - name: libwebp7
        - name: libstdc++6
        - name: libgcc-s1
        - name: liblerc4
        - name: libjbig0
        - name: libdeflate0
        - name: tzdata
        - name: libtinfo6
        - name: readline-common
        - name: libpango-1.0-0
        - name: libharfbuzz0b
        - name: libglib2.0-0t64
        - name: libpangoft2-1.0-0
        - name: libthai0
        - name: libfribidi0
        - name: fontconfig
        - name: libjpeg-turbo8
        - name: gcc-14-base
        - name: libpcre2-8-0
        - name: libmount1
        - name: libgssapi-krb5-2
        - name: libssh-4
        - name: librtmp1
        - name: libpsl5t64
        - name: libnghttp2-14
        - name: libldap2
        - name: libbrotli1
        - name: libxcb1
        - name: libxcb-shm0
        - name: libxcb-render0
        - name: libpixman-1-0
        - name: libpng16-16t64
        - name: libgfortran5
        - name: ucf
        - name: libpaper1
        - name: r-cran-lattice
        - name: r-base-core
        - name: libopenblas0-serial
        - name: libopenblas0-pthread
        - name: libopenblas0-openmp
        - name: liblapack3
        - name: libatlas3-base
        - name: libblis4-serial
        - name: libblis4-pthread
        - name: libblis4-openmp
        - name: libblas3
        - name: r-cran-matrix
        - name: r-cran-codetools
        - name: r-cran-spatial
        - name: r-cran-nnet
        - name: r-cran-mass
        - name: r-cran-class
        - name: r-cran-survival
        - name: r-cran-rpart
        - name: r-cran-nlme
        - name: libgomp1
        - name: r-cran-mgcv
        - name: r-cran-kernsmooth
        - name: r-cran-foreign
        - name: r-cran-cluster
        - name: r-cran-boot
        - name: ca-certificates
        - name: libxt6t64
        - name: libtcl8.6
        - name: libtk8.6
        - name: libtirpc3t64
        - name: libjpeg8
        - name: libtiff6
        - name: libcairo2
        - name: libpangocairo-1.0-0
        - name: libicu74
        - name: libcurl4t64
        - name: xdg-utils
        - name: libpaper-utils
        - name: unzip
        - name: zip
        - name: r-recommended
    - name: r-base-dev
      dependencies:
        - name: liblapack.so.3
          optional: true
        - name: libhogweed6
          optional: true
        - name: libgnutls30
          optional: true
        - name: libblas.so.3
          optional: true
        - name: libblas.so
          optional: true
        - name: libglib2.0-0
          optional: true
        - name: libreadline8
          optional: true
        - name: libc-dev
          optional: true
        - name: libnettle8
          optional: true
        - name: perl:any
          optional: true
        - name: debconf-2.0
          optional: true
        - name: x11-common
        - name: xfonts-encodings
        - name: zlib1g
        - name: libc6
        - name: libfontenc1
        - name: fonts-noto-mono
        - name: libfreetype6
        - name: xfonts-utils
        - name: fonts-dejavu-mono
        - name: gcc-14-base
        - name: libquadmath0
        - name: libgcc-s1
        - name: libhwasan0
        - name: libstdc++6
        - name: libubsan1
        - name: libtsan2
        - name: liblsan0
        - name: libasan8
        - name: libatomic1
        - name: libitm1
        - name: libjansson4
        - name: libbinutils
        - name: libgprofng0
        - name: libctf0
        - name: libctf-nobfd0
        - name: libsframe1
        - name: libgdbm6t64
        - name: libgdbm-compat4t64
        - name: libdb5.3t64
        - name: sysvinit-utils
        - name: lsb-base
        - name: libsasl2-modules-db
        - name: libtasn1-6
        - name: libffi8
        - name: libp11-kit0
        - name: libkeyutils1
        - name: fonts-noto-core
        - name: fonts-texgyre
        - name: fonts-urw-base35
        - name: fonts-freefont-ttf
        - name: fonts-freefont-otf
        - name: fonts-croscore
        - name: fonts-liberation
        - name: fonts-dejavu-core
        - name: libmd0
        - name: libgfortran5
        - name: libgcc-13-dev
        - name: gcc-13-base
        - name: libgfortran-13-dev
        - name: libc6-dev
        - name: libstdc++-13-dev
        - name: libgomp1
        - name: libcc1-0
        - name: libgmp10
        - name: libmpfr6
        - name: libmpc3
        - name: libisl23
        - name: libzstd1
        - name: binutils-common
        - name: binutils-x86-64-linux-gnu
        - name: libacl1
        - name: perl-modules-5.38
        - name: libcrypt1
        - name: libbz2-1.0
        - name: libperl5.38t64
        - name: perl-base
        - name: dpkg
        - name: libgc1
        - name: libtextwrap1
        - name: libslang2
        - name: libnewt0.52
        - name: libdebian-installer4
        - name: libuuid1
        - name: libbsd0
        - name: libsharpyuv0
        - name: libdatrie1
        - name: libthai-data
        - name: libgraphite2-3
        - name: libblkid1
        - name: libnettle8t64
        - name: libhogweed6t64
        - name: libssl3t64
        - name: libsasl2-2
        - name: libunistring5
        - name: libidn2-0
        - name: libgnutls30t64
        - name: libkrb5support0
        - name: libk5crypto3
        - name: libcom-err2
        - name: libkrb5-3
        - name: libxdmcp6
        - name: fontconfig-config
        - name: libexpat1
        - name: tar
        - name: libselinux1
        - name: liblzma5
        - name: libpkgconf3
        - name: gcc-13-x86-64-linux-gnu
        - name: gfortran-13-x86-64-linux-gnu
        - name: g++-13-x86-64-linux-gnu
        - name: cpp-13-x86-64-linux-gnu
        - name: cpp-13
        - name: lto-disabled-list
        - name: binutils
        - name: patch
        - name: xz-utils
        - name: bzip2
        - name: perl
        - name: libdpkg-perl
        - name: libreadline8t64
        - name: guile-3.0-libs
        - name: rpcsvc-proto
        - name: libcrypt-dev
        - name: linux-libc-dev
        - name: libc-dev-bin
        - name: openssl
        - name: sensible-utils
        - name: debconf
        - name: cdebconf
        - name: libice6
        - name: libsm6
        - name: libx11-data
        - name: libxext6
        - name: libx11-6
        - name: libxss1
        - name: libxrender1
        - name: libfontconfig1
        - name: libxft2
        - name: libtirpc-common
        - name: libwebp7
        - name: liblerc4
        - name: libjbig0
        - name: libdeflate0
        - name: tzdata
        - name: readline-common
        - name: libpango-1.0-0
        - name: libharfbuzz0b
        - name: libglib2.0-0t64
        - name: libpangoft2-1.0-0
        - name: libthai0
        - name: libfribidi0
        - name: fontconfig
        - name: libpcre2-8-0
        - name: libmount1
        - name: libgssapi-krb5-2
        - name: libssh-4
        - name: librtmp1
        - name: libpsl5t64
        - name: libnghttp2-14
        - name: libldap2
        - name: libbrotli1
        - name: libxau6
        - name: libxcb1
        - name: libxcb-shm0
        - name: libxcb-render0
        - name: libpixman-1-0
        - name: libpng16-16t64
        - name: ucf
        - name: libpaper1
        - name: pkgconf-bin
        - name: libxmuu1
        - name: libicu74
        - name: icu-devtools
        - name: libpcre2-posix3
        - name: libpcre2-32-0
        - name: libpcre2-16-0
        - name: libjpeg-turbo8
        - name: libjpeg8
        - name: libjpeg-turbo8-dev
        - name: libjpeg8-dev
        - name: libtinfo6
        - name: libncursesw6
        - name: libncurses6
        - name: libopenblas0-serial
        - name: libopenblas-serial-dev
        - name: libopenblas0-pthread
        - name: libopenblas-pthread-dev
        - name: libopenblas0-openmp
        - name: libopenblas-openmp-dev
        - name: libblis4-serial
        - name: libblis-serial-dev
        - name: libblis4-pthread
        - name: libblis-pthread-dev
        - name: libblis4-openmp
        - name: libblis-openmp-dev
        - name: gcc-x86-64-linux-gnu
        - name: cpp-x86-64-linux-gnu
        - name: gfortran-x86-64-linux-gnu
        - name: gcc-13
        - name: gfortran-13
        - name: g++-x86-64-linux-gnu
        - name: g++-13
        - name: cpp
        - name: make-guile
        - name: make
        - name: dpkg-dev
        - name: ca-certificates
        - name: libxt6t64
        - name: libtcl8.6
        - name: libtk8.6
        - name: libtirpc3t64
        - name: libtiff6
        - name: libcairo2
        - name: libpangocairo-1.0-0
        - name: libblas3
        - name: libatlas3-base
        - name: liblapack3
        - name: libcurl4t64
        - name: xdg-utils
        - name: libpaper-utils
        - name: unzip
        - name: zip
        - name: pkgconf
        - name: pkg-config
        - name: xauth
        - name: libicu-dev
        - name: liblzma-dev
        - name: libbz2-dev
        - name: zlib1g-dev
        - name: libpng-dev
        - name: libpcre2-dev
        - name: libjpeg-dev
        - name: libncurses-dev
        - name: libreadline-dev
        - name: libblas-dev
        - name: libatlas-base-dev
        - name: liblapack-dev
        - name: gcc
        - name: gfortran
        - name: g++
        - name: build-essential
        - name: r-base-core
    - name: r-base
cran:
    - package_name: BiocManager
      package_path: BiocManager_1.30.27.tar.gz
    - package_name: Seurat
      package_path: Seurat_5.4.0.tar.gz
      bioconductor: true
      dependencies:
        - package_name: cluster
          package_path: cluster_2.1.8.2.tar.gz
          bioconductor: true
        - package_name: fitdistrplus
          package_path: fitdistrplus_1.2-6.tar.gz
          bioconductor: true
          dependencies:
            - package_name: rlang
              package_path: rlang_1.1.7.tar.gz
              bioconductor: true
            - package_name: survival
              package_path: survival_3.8-6.tar.gz
              bioconductor: true
        - package_name: future
          package_path: future_1.69.0.tar.gz
          bioconductor: true
          dependencies:
            - package_name: digest
              package_path: digest_0.6.39.tar.gz
              bioconductor: true
            - package_name: codetools
              package_path: codetools_0.2-20.tar.gz
              bioconductor: true
        - package_name: future.apply
          package_path: future.apply_1.20.2.tar.gz
          bioconductor: true
          dependencies:
            - package_name: globals
              package_path: globals_0.19.0.tar.gz
              bioconductor: true
              dependencies:
                - package_name: codetools
                  package_path: codetools_0.2-20.tar.gz
                  bioconductor: true
            - package_name: digest
              package_path: digest_0.6.39.tar.gz
              bioconductor: true
            - package_name: listenv
              package_path: listenv_0.10.1.tar.gz
              bioconductor: true
            - package_name: parallelly
              package_path: parallelly_1.46.1.tar.gz
              bioconductor: true
            - package_name: codetools
              package_path: codetools_0.2-20.tar.gz
              bioconductor: true
            - package_name: future
              package_path: future_1.69.0.tar.gz
              bioconductor: true
              dependencies:
                - package_name: digest
                  package_path: digest_0.6.39.tar.gz
                  bioconductor: true
                - package_name: codetools
                  package_path: codetools_0.2-20.tar.gz
                  bioconductor: true
        - package_name: generics
          package_path: generics_0.1.4.tar.gz
          bioconductor: true
        - package_name: SeuratObject
          package_path: SeuratObject_5.3.0.tar.gz
          bioconductor: true
          dependencies:
            - package_name: generics
              package_path: generics_0.1.4.tar.gz
              bioconductor: true
            - package_name: future
              package_path: future_1.69.0.tar.gz
              bioconductor: true
              dependencies:
                - package_name: digest
                  package_path: digest_0.6.39.tar.gz
                  bioconductor: true
                - package_name: codetools
                  package_path: codetools_0.2-20.tar.gz
                  bioconductor: true
            - package_name: sp
              package_path: sp_2.2-1.tar.gz
              bioconductor: true
              dependencies:
                - package_name: lattice
                  package_path: lattice_0.22-9.tar.gz
                  bioconductor: true
            - package_name: lifecycle
              package_path: lifecycle_1.0.5.tar.gz
              bioconductor: true
              dependencies:
                - package_name: cli
                  package_path: cli_3.6.5.tar.gz
                  bioconductor: true
                - package_name: rlang
                  package_path: rlang_1.1.7.tar.gz
                  bioconductor: true
            - package_name: rlang
              package_path: rlang_1.1.7.tar.gz
              bioconductor: true
            - package_name: progressr
              package_path: progressr_0.18.0.tar.gz
              bioconductor: true
              dependencies:
                - package_name: digest
                  package_path: digest_0.6.39.tar.gz
                  bioconductor: true
            - package_name: future.apply
              package_path: future.apply_1.20.2.tar.gz
              bioconductor: true
              dependencies:
                - package_name: listenv
                  package_path: listenv_0.10.1.tar.gz
                  bioconductor: true
                - package_name: codetools
                  package_path: codetools_0.2-20.tar.gz
                  bioconductor: true
                - package_name: parallelly
                  package_path: parallelly_1.46.1.tar.gz
                  bioconductor: true
                - package_name: future
                  package_path: future_1.69.0.tar.gz
                  bioconductor: true
                  dependencies:
                    - package_name: listenv
                      package_path: listenv_0.10.1.tar.gz
                      bioconductor: true
                    - package_name: globals
                      package_path: globals_0.19.0.tar.gz
                      bioconductor: true
                      dependencies:
                        - package_name: codetools
                          package_path: codetools_0.2-20.tar.gz
                          bioconductor: true
                    - package_name: codetools
                      package_path: codetools_0.2-20.tar.gz
                      bioconductor: true
                    - package_name: digest
                      package_path: digest_0.6.39.tar.gz
                      bioconductor: true
                    - package_name: parallelly
                      package_path: parallelly_1.46.1.tar.gz
                      bioconductor: true
            - package_name: Rcpp
              package_path: Rcpp_1.1.1.tar.gz
              bioconductor: true
            - package_name: digest
              package_path: digest_0.6.39.tar.gz
              bioconductor: true
            - package_name: globals
              package_path: globals_0.19.0.tar.gz
              bioconductor: true
              dependencies:
                - package_name: codetools
                  package_path: codetools_0.2-20.tar.gz
                  bioconductor: true
            - package_name: listenv
              package_path: listenv_0.10.1.tar.gz
              bioconductor: true
            - package_name: parallelly
              package_path: parallelly_1.46.1.tar.gz
              bioconductor: true
            - package_name: cli
              package_path: cli_3.6.5.tar.gz
              bioconductor: true
            - package_name: lattice
              package_path: lattice_0.22-9.tar.gz
              bioconductor: true
            - package_name: codetools
              package_path: codetools_0.2-20.tar.gz
              bioconductor: true
            - package_name: dotCall64
              package_path: dotCall64_1.2.tar.gz
              bioconductor: true
            - package_name: spam
              package_path: spam_2.11-3.tar.gz
              bioconductor: true
              dependencies:
                - package_name: Rcpp
                  package_path: Rcpp_1.1.1.tar.gz
                  bioconductor: true
                - package_name: dotCall64
                  package_path: dotCall64_1.2.tar.gz
                  bioconductor: true
            - package_name: RcppEigen
              package_path: RcppEigen_0.3.4.0.2.tar.gz
              bioconductor: true
              dependencies:
                - package_name: Rcpp
                  package_path: Rcpp_1.1.1.tar.gz
                  bioconductor: true
        - package_name: cowplot
          package_path: cowplot_1.2.0.tar.gz
          bioconductor: true
          dependencies:
            - package_name: gtable
              package_path: gtable_0.3.6.tar.gz
              bioconductor: true
              dependencies:
                - package_name: rlang
                  package_path: rlang_1.1.7.tar.gz
                  bioconductor: true
            - package_name: lifecycle
              package_path: lifecycle_1.0.5.tar.gz
              bioconductor: true
              dependencies:
                - package_name: rlang
                  package_path: rlang_1.1.7.tar.gz
                  bioconductor: true
            - package_name: S7
              package_path: S7_0.2.1.tar.gz
              bioconductor: true
            - package_name: isoband
              package_path: isoband_0.3.0.tar.gz
              bioconductor: true
              dependencies:
                - package_name: rlang
                  package_path: rlang_1.1.7.tar.gz
                  bioconductor: true
                - package_name: cpp11
                  package_path: cpp11_0.5.3.tar.gz
                  bioconductor: true
            - package_name: glue
              package_path: glue_1.8.0.tar.gz
              bioconductor: true
            - package_name: vctrs
              package_path: vctrs_0.7.1.tar.gz
              bioconductor: true
              dependencies:
                - package_name: rlang
                  package_path: rlang_1.1.7.tar.gz
                  bioconductor: true
                - package_name: glue
                  package_path: glue_1.8.0.tar.gz
                  bioconductor: true
                - package_name: lifecycle
                  package_path: lifecycle_1.0.5.tar.gz
                  bioconductor: true
                  dependencies:
                    - package_name: cli
                      package_path: cli_3.6.5.tar.gz
                      bioconductor: true
                - package_name: cli
                  package_path: cli_3.6.5.tar.gz
                  bioconductor: true
            - package_name: withr
              package_path: withr_3.0.2.tar.gz
              bioconductor: true
            - package_name: farver
              package_path: farver_2.1.2.tar.gz
              bioconductor: true
            - package_name: RColorBrewer
              package_path: RColorBrewer_1.1-3.tar.gz
              bioconductor: true
            - package_name: labeling
              package_path: labeling_0.4.3.tar.gz
              bioconductor: true
            - package_name: cpp11
              package_path: cpp11_0.5.3.tar.gz
              bioconductor: true
            - package_name: R6
              package_path: R6_2.6.1.tar.gz
              bioconductor: true
            - package_name: viridisLite
              package_path: viridisLite_0.4.3.tar.gz
              bioconductor: true
            - package_name: ggplot2
              package_path: ggplot2_4.0.2.tar.gz
              bioconductor: true
              dependencies:
                - package_name: lifecycle
                  package_path: lifecycle_1.0.5.tar.gz
                  bioconductor: true
                  dependencies:
                    - package_name: cli
                      package_path: cli_3.6.5.tar.gz
                      bioconductor: true
                - package_name: gtable
                  package_path: gtable_0.3.6.tar.gz
                  bioconductor: true
                  dependencies:
                    - package_name: glue
                      package_path: glue_1.8.0.tar.gz
                      bioconductor: true
                    - package_name: rlang
                      package_path: rlang_1.1.7.tar.gz
                      bioconductor: true
                    - package_name: lifecycle
                      package_path: lifecycle_1.0.5.tar.gz
                      bioconductor: true
                - package_name: rlang
                  package_path: rlang_1.1.7.tar.gz
                  bioconductor: true
                - package_name: S7
                  package_path: S7_0.2.1.tar.gz
                  bioconductor: true
                - package_name: isoband
                  package_path: isoband_0.3.0.tar.gz
                  bioconductor: true
                  dependencies:
                    - package_name: rlang
                      package_path: rlang_1.1.7.tar.gz
                      bioconductor: true
                    - package_name: cpp11
                      package_path: cpp11_0.5.3.tar.gz
                      bioconductor: true
                - package_name: glue
                  package_path: glue_1.8.0.tar.gz
                  bioconductor: true
                - package_name: cpp11
                  package_path: cpp11_0.5.3.tar.gz
                  bioconductor: true
                - package_name: vctrs
                  package_path: vctrs_0.7.1.tar.gz
                  bioconductor: true
                  dependencies:
                    - package_name: rlang
                      package_path: rlang_1.1.7.tar.gz
                      bioconductor: true
                    - package_name: cli
                      package_path: cli_3.6.5.tar.gz
                      bioconductor: true
                    - package_name: lifecycle
                      package_path: lifecycle_1.0.5.tar.gz
                      bioconductor: true
                      dependencies:
                        - package_name: cli
                          package_path: cli_3.6.5.tar.gz
                          bioconductor: true
                    - package_name: glue
                      package_path: glue_1.8.0.tar.gz
                      bioconductor: true
                - package_name: farver
                  package_path: farver_2.1.2.tar.gz
                  bioconductor: true
                - package_name: RColorBrewer
                  package_path: RColorBrewer_1.1-3.tar.gz
                  bioconductor: true
                - package_name: labeling
                  package_path: labeling_0.4.3.tar.gz
                  bioconductor: true
                - package_name: scales
                  package_path: scales_1.4.0.tar.gz
                  bioconductor: true
                  dependencies:
                    - package_name: glue
                      package_path: glue_1.8.0.tar.gz
                      bioconductor: true
                    - package_name: lifecycle
                      package_path: lifecycle_1.0.5.tar.gz
                      bioconductor: true
                      dependencies:
                        - package_name: cli
                          package_path: cli_3.6.5.tar.gz
                          bioconductor: true
                    - package_name: cli
                      package_path: cli_3.6.5.tar.gz
                      bioconductor: true
                    - package_name: RColorBrewer
                      package_path: RColorBrewer_1.1-3.tar.gz
                      bioconductor: true
                    - package_name: rlang
                      package_path: rlang_1.1.7.tar.gz
                      bioconductor: true
                    - package_name: farver
                      package_path: farver_2.1.2.tar.gz
                      bioconductor: true
                    - package_name: viridisLite
                      package_path: viridisLite_0.4.3.tar.gz
                      bioconductor: true
                    - package_name: labeling
                      package_path: labeling_0.4.3.tar.gz
                      bioconductor: true
                - package_name: R6
                  package_path: R6_2.6.1.tar.gz
                  bioconductor: true
                - package_name: viridisLite
                  package_path: viridisLite_0.4.3.tar.gz
                  bioconductor: true
        - package_name: fastDummies
          package_path: fastDummies_1.7.5.tar.gz
          bioconductor: true
          dependencies:
            - package_name: data.table
              package_path: data.table_1.18.2.1.tar.gz
              bioconductor: true
            - package_name: glue
              package_path: glue_1.8.0.tar.gz
              bioconductor: true
            - package_name: magrittr
              package_path: magrittr_2.0.4.tar.gz
              bioconductor: true
            - package_name: rlang
              package_path: rlang_1.1.7.tar.gz
              bioconductor: true
            - package_name: lifecycle
              package_path: lifecycle_1.0.5.tar.gz
              bioconductor: true
              dependencies:
                - package_name: cli
                  package_path: cli_3.6.5.tar.gz
                  bioconductor: true
                - package_name: rlang
                  package_path: rlang_1.1.7.tar.gz
                  bioconductor: true
            - package_name: stringr
              package_path: stringr_1.6.0.tar.gz
              bioconductor: true
              dependencies:
                - package_name: magrittr
                  package_path: magrittr_2.0.4.tar.gz
                  bioconductor: true
                - package_name: glue
                  package_path: glue_1.8.0.tar.gz
                  bioconductor: true
                - package_name: lifecycle
                  package_path: lifecycle_1.0.5.tar.gz
                  bioconductor: true
                  dependencies:
                    - package_name: cli
                      package_path: cli_3.6.5.tar.gz
                      bioconductor: true
                - package_name: stringi
                  package_path: stringi_1.8.7.tar.gz
                  bioconductor: true
                  external_dependencies:
                    apt:
                        - name: libicu-dev
                          dependencies:
                            - name: libc-dev
                              optional: true
                            - name: libc6
                            - name: libcrypt1
                            - name: gcc-14-base
                            - name: rpcsvc-proto
                            - name: libcrypt-dev
                            - name: linux-libc-dev
                            - name: libc-dev-bin
                            - name: libgcc-s1
                            - name: libstdc++6
                            - name: libc6-dev
                            - name: libicu74
                            - name: icu-devtools
                - package_name: rlang
                  package_path: rlang_1.1.7.tar.gz
                  bioconductor: true
            - package_name: tibble
              package_path: tibble_3.3.1.tar.gz
              bioconductor: true
              dependencies:
                - package_name: magrittr
                  package_path: magrittr_2.0.4.tar.gz
                  bioconductor: true
                - package_name: rlang
                  package_path: rlang_1.1.7.tar.gz
                  bioconductor: true
                - package_name: pkgconfig
                  package_path: pkgconfig_2.0.3.tar.gz
                  bioconductor: true
                - package_name: lifecycle
                  package_path: lifecycle_1.0.5.tar.gz
                  bioconductor: true
                - package_name: glue
                  package_path: glue_1.8.0.tar.gz
                  bioconductor: true
                - package_name: pillar
                  package_path: pillar_1.11.1.tar.gz
                  bioconductor: true
                  dependencies:
                    - package_name: rlang
                      package_path: rlang_1.1.7.tar.gz
                      bioconductor: true
                    - package_name: glue
                      package_path: glue_1.8.0.tar.gz
                      bioconductor: true
                    - package_name: utf8
                      package_path: utf8_1.2.6.tar.gz
                      bioconductor: true
                    - package_name: lifecycle
                      package_path: lifecycle_1.0.5.tar.gz
                      bioconductor: true
                      dependencies:
                        - package_name: rlang
                          package_path: rlang_1.1.7.tar.gz
                          bioconductor: true
                        - package_name: cli
                          package_path: cli_3.6.5.tar.gz
                          bioconductor: true
                - package_name: utf8
                  package_path: utf8_1.2.6.tar.gz
                  bioconductor: true
                - package_name: vctrs
                  package_path: vctrs_0.7.1.tar.gz
                  bioconductor: true
                  dependencies:
                    - package_name: rlang
                      package_path: rlang_1.1.7.tar.gz
                      bioconductor: true
                    - package_name: glue
                      package_path: glue_1.8.0.tar.gz
                      bioconductor: true
                    - package_name: cli
                      package_path: cli_3.6.5.tar.gz
                      bioconductor: true
                    - package_name: lifecycle
                      package_path: lifecycle_1.0.5.tar.gz
                      bioconductor: true
                      dependencies:
                        - package_name: cli
                          package_path: cli_3.6.5.tar.gz
                          bioconductor: true
            - package_name: pkgconfig
              package_path: pkgconfig_2.0.3.tar.gz
              bioconductor: true
            - package_name: vctrs
              package_path: vctrs_0.7.1.tar.gz
              bioconductor: true
              dependencies:
                - package_name: rlang
                  package_path: rlang_1.1.7.tar.gz
                  bioconductor: true
                - package_name: glue
                  package_path: glue_1.8.0.tar.gz
                  bioconductor: true
                - package_name: cli
                  package_path: cli_3.6.5.tar.gz
                  bioconductor: true
                - package_name: lifecycle
                  package_path: lifecycle_1.0.5.tar.gz
                  bioconductor: true
                  dependencies:
                    - package_name: cli
                      package_path: cli_3.6.5.tar.gz
                      bioconductor: true
            - package_name: utf8
              package_path: utf8_1.2.6.tar.gz
              bioconductor: true
            - package_name: stringi
              package_path: stringi_1.8.7.tar.gz
              bioconductor: true
              external_dependencies:
                apt:
                    - name: libicu-dev
                      dependencies:
                        - name: libc-dev
                          optional: true
                        - name: libc6
                        - name: libcrypt1
                        - name: gcc-14-base
                        - name: rpcsvc-proto
                        - name: libcrypt-dev
                        - name: linux-libc-dev
                        - name: libc-dev-bin
                        - name: libgcc-s1
                        - name: libstdc++6
                        - name: libc6-dev
                        - name: libicu74
                        - name: icu-devtools
            - package_name: pillar
              package_path: pillar_1.11.1.tar.gz
              bioconductor: true
              dependencies:
                - package_name: rlang
                  package_path: rlang_1.1.7.tar.gz
                  bioconductor: true
                - package_name: utf8
                  package_path: utf8_1.2.6.tar.gz
                  bioconductor: true
                - package_name: cli
                  package_path: cli_3.6.5.tar.gz
                  bioconductor: true
                - package_name: lifecycle
                  package_path: lifecycle_1.0.5.tar.gz
                  bioconductor: true
                  dependencies:
                    - package_name: cli
                      package_path: cli_3.6.5.tar.gz
                      bioconductor: true
                - package_name: glue
                  package_path: glue_1.8.0.tar.gz
                  bioconductor: true
                - package_name: vctrs
                  package_path: vctrs_0.7.1.tar.gz
                  bioconductor: true
                  dependencies:
                    - package_name: rlang
                      package_path: rlang_1.1.7.tar.gz
                      bioconductor: true
                    - package_name: glue
                      package_path: glue_1.8.0.tar.gz
                      bioconductor: true
                    - package_name: lifecycle
                      package_path: lifecycle_1.0.5.tar.gz
                      bioconductor: true
                      dependencies:
                        - package_name: cli
                          package_path: cli_3.6.5.tar.gz
                          bioconductor: true
                    - package_name: cli
                      package_path: cli_3.6.5.tar.gz
                      bioconductor: true
        - package_name: ggplot2
          package_path: ggplot2_4.0.2.tar.gz
          bioconductor: true
          dependencies:
            - package_name: gtable
              package_path: gtable_0.3.6.tar.gz
              bioconductor: true
              dependencies:
                - package_name: rlang
                  package_path: rlang_1.1.7.tar.gz
                  bioconductor: true
            - package_name: isoband
              package_path: isoband_0.3.0.tar.gz
              bioconductor: true
              dependencies:
                - package_name: rlang
                  package_path: rlang_1.1.7.tar.gz
                  bioconductor: true
                - package_name: cpp11
                  package_path: cpp11_0.5.3.tar.gz
                  bioconductor: true
            - package_name: cli
              package_path: cli_3.6.5.tar.gz
              bioconductor: true
            - package_name: rlang
              package_path: rlang_1.1.7.tar.gz
              bioconductor: true
            - package_name: S7
              package_path: S7_0.2.1.tar.gz
              bioconductor: true
            - package_name: lifecycle
              package_path: lifecycle_1.0.5.tar.gz
              bioconductor: true
              dependencies:
                - package_name: cli
                  package_path: cli_3.6.5.tar.gz
                  bioconductor: true
            - package_name: vctrs
              package_path: vctrs_0.7.1.tar.gz
              bioconductor: true
              dependencies:
                - package_name: rlang
                  package_path: rlang_1.1.7.tar.gz
                  bioconductor: true
                - package_name: glue
                  package_path: glue_1.8.0.tar.gz
                  bioconductor: true
                - package_name: cli
                  package_path: cli_3.6.5.tar.gz
                  bioconductor: true
                - package_name: lifecycle
                  package_path: lifecycle_1.0.5.tar.gz
                  bioconductor: true
                  dependencies:
                    - package_name: cli
                      package_path: cli_3.6.5.tar.gz
                      bioconductor: true
            - package_name: withr
              package_path: withr_3.0.2.tar.gz
              bioconductor: true
            - package_name: glue
              package_path: glue_1.8.0.tar.gz
              bioconductor: true
            - package_name: cpp11
              package_path: cpp11_0.5.3.tar.gz
              bioconductor: true
            - package_name: farver
              package_path: farver_2.1.2.tar.gz
              bioconductor: true
            - package_name: labeling
              package_path: labeling_0.4.3.tar.gz
              bioconductor: true
            - package_name: R6
              package_path: R6_2.6.1.tar.gz
              bioconductor: true
            - package_name: RColorBrewer
              package_path: RColorBrewer_1.1-3.tar.gz
              bioconductor: true
            - package_name: viridisLite
              package_path: viridisLite_0.4.3.tar.gz
              bioconductor: true
            - package_name: scales
              package_path: scales_1.4.0.tar.gz
              bioconductor: true
              dependencies:
                - package_name: labeling
                  package_path: labeling_0.4.3.tar.gz
                  bioconductor: true
                - package_name: lifecycle
                  package_path: lifecycle_1.0.5.tar.gz
                  bioconductor: true
                  dependencies:
                    - package_name: cli
                      package_path: cli_3.6.5.tar.gz
                      bioconductor: true
                - package_name: cli
                  package_path: cli_3.6.5.tar.gz
                  bioconductor: true
                - package_name: RColorBrewer
                  package_path: RColorBrewer_1.1-3.tar.gz
                  bioconductor: true
                - package_name: rlang
                  package_path: rlang_1.1.7.tar.gz
                  bioconductor: true
                - package_name: viridisLite
                  package_path: viridisLite_0.4.3.tar.gz
                  bioconductor: true
                - package_name: glue
                  package_path: glue_1.8.0.tar.gz
                  bioconductor: true
                - package_name: farver
                  package_path: farver_2.1.2.tar.gz
                  bioconductor: true
                - package_name: R6
                  package_path: R6_2.6.1.tar.gz
                  bioconductor: true
        - package_name: ggridges
          package_path: ggridges_0.5.7.tar.gz
          bioconductor: true
          dependencies:
            - package_name: withr
              package_path: withr_3.0.2.tar.gz
              bioconductor: true
            - package_name: cli
              package_path: cli_3.6.5.tar.gz
              bioconductor: true
            - package_name: ggplot2
              package_path: ggplot2_4.0.2.tar.gz
              bioconductor: true
              dependencies:
                - package_name: lifecycle
                  package_path: lifecycle_1.0.5.tar.gz
                  bioconductor: true
                  dependencies:
                    - package_name: cli
                      package_path: cli_3.6.5.tar.gz
                      bioconductor: true
                - package_name: gtable
                  package_path: gtable_0.3.6.tar.gz
                  bioconductor: true
                  dependencies:
                    - package_name: glue
                      package_path: glue_1.8.0.tar.gz
                      bioconductor: true
                    - package_name: rlang
                      package_path: rlang_1.1.7.tar.gz
                      bioconductor: true
                    - package_name: lifecycle
                      package_path: lifecycle_1.0.5.tar.gz
                      bioconductor: true
                - package_name: rlang
                  package_path: rlang_1.1.7.tar.gz
                  bioconductor: true
                - package_name: S7
                  package_path: S7_0.2.1.tar.gz
                  bioconductor: true
                - package_name: isoband
                  package_path: isoband_0.3.0.tar.gz
                  bioconductor: true
                  dependencies:
                    - package_name: rlang
                      package_path: rlang_1.1.7.tar.gz
                      bioconductor: true
                    - package_name: cpp11
                      package_path: cpp11_0.5.3.tar.gz
                      bioconductor: true
                - package_name: glue
                  package_path: glue_1.8.0.tar.gz
                  bioconductor: true
                - package_name: cpp11
                  package_path: cpp11_0.5.3.tar.gz
                  bioconductor: true
                - package_name: vctrs
                  package_path: vctrs_0.7.1.tar.gz
                  bioconductor: true
                  dependencies:
                    - package_name: rlang
                      package_path: rlang_1.1.7.tar.gz
                      bioconductor: true
                    - package_name: cli
                      package_path: cli_3.6.5.tar.gz
                      bioconductor: true
                    - package_name: lifecycle
                      package_path: lifecycle_1.0.5.tar.gz
                      bioconductor: true
                      dependencies:
                        - package_name: cli
                          package_path: cli_3.6.5.tar.gz
                          bioconductor: true
                    - package_name: glue
                      package_path: glue_1.8.0.tar.gz
                      bioconductor: true
                - package_name: farver
                  package_path: farver_2.1.2.tar.gz
                  bioconductor: true
                - package_name: RColorBrewer
                  package_path: RColorBrewer_1.1-3.tar.gz
                  bioconductor: true
                - package_name: labeling
                  package_path: labeling_0.4.3.tar.gz
                  bioconductor: true
                - package_name: scales
                  package_path: scales_1.4.0.tar.gz
                  bioconductor: true
                  dependencies:
                    - package_name: glue
                      package_path: glue_1.8.0.tar.gz
                      bioconductor: true
                    - package_name: lifecycle
                      package_path: lifecycle_1.0.5.tar.gz
                      bioconductor: true
                      dependencies:
                        - package_name: cli
                          package_path: cli_3.6.5.tar.gz
                          bioconductor: true
                    - package_name: cli
                      package_path: cli_3.6.5.tar.gz
                      bioconductor: true
                    - package_name: RColorBrewer
                      package_path: RColorBrewer_1.1-3.tar.gz
                      bioconductor: true
                    - package_name: rlang
                      package_path: rlang_1.1.7.tar.gz
                      bioconductor: true
                    - package_name: farver
                      package_path: farver_2.1.2.tar.gz
                      bioconductor: true
                    - package_name: viridisLite
                      package_path: viridisLite_0.4.3.tar.gz
                      bioconductor: true
                    - package_name: labeling
                      package_path: labeling_0.4.3.tar.gz
                      bioconductor: true
                - package_name: R6
                  package_path: R6_2.6.1.tar.gz
                  bioconductor: true
                - package_name: viridisLite
                  package_path: viridisLite_0.4.3.tar.gz
                  bioconductor: true
            - package_name: gtable
              package_path: gtable_0.3.6.tar.gz
              bioconductor: true
              dependencies:
                - package_name: rlang
                  package_path: rlang_1.1.7.tar.gz
                  bioconductor: true
            - package_name: isoband
              package_path: isoband_0.3.0.tar.gz
              bioconductor: true
              dependencies:
                - package_name: rlang
                  package_path: rlang_1.1.7.tar.gz
                  bioconductor: true
                - package_name: cpp11
                  package_path: cpp11_0.5.3.tar.gz
                  bioconductor: true
            - package_name: lifecycle
              package_path: lifecycle_1.0.5.tar.gz
              bioconductor: true
              dependencies:
                - package_name: cli
                  package_path: cli_3.6.5.tar.gz
                  bioconductor: true
            - package_name: rlang
              package_path: rlang_1.1.7.tar.gz
              bioconductor: true
            - package_name: S7
              package_path: S7_0.2.1.tar.gz
              bioconductor: true
            - package_name: scales
              package_path: scales_1.4.0.tar.gz
              bioconductor: true
              dependencies:
                - package_name: glue
                  package_path: glue_1.8.0.tar.gz
                  bioconductor: true
                - package_name: lifecycle
                  package_path: lifecycle_1.0.5.tar.gz
                  bioconductor: true
                  dependencies:
                    - package_name: cli
                      package_path: cli_3.6.5.tar.gz
                      bioconductor: true
                - package_name: cli
                  package_path: cli_3.6.5.tar.gz
                  bioconductor: true
                - package_name: RColorBrewer
                  package_path: RColorBrewer_1.1-3.tar.gz
                  bioconductor: true
                - package_name: rlang
                  package_path: rlang_1.1.7.tar.gz
                  bioconductor: true
                - package_name: farver
                  package_path: farver_2.1.2.tar.gz
                  bioconductor: true
                - package_name: viridisLite
                  package_path: viridisLite_0.4.3.tar.gz
                  bioconductor: true
                - package_name: labeling
                  package_path: labeling_0.4.3.tar.gz
                  bioconductor: true
            - package_name: vctrs
              package_path: vctrs_0.7.1.tar.gz
              bioconductor: true
              dependencies:
                - package_name: rlang
                  package_path: rlang_1.1.7.tar.gz
                  bioconductor: true
                - package_name: glue
                  package_path: glue_1.8.0.tar.gz
                  bioconductor: true
                - package_name: cli
                  package_path: cli_3.6.5.tar.gz
                  bioconductor: true
                - package_name: lifecycle
                  package_path: lifecycle_1.0.5.tar.gz
                  bioconductor: true
                  dependencies:
                    - package_name: cli
                      package_path: cli_3.6.5.tar.gz
                      bioconductor: true
            - package_name: farver
              package_path: farver_2.1.2.tar.gz
              bioconductor: true
            - package_name: glue
              package_path: glue_1.8.0.tar.gz
              bioconductor: true
            - package_name: labeling
              package_path: labeling_0.4.3.tar.gz
              bioconductor: true
            - package_name: R6
              package_path: R6_2.6.1.tar.gz
              bioconductor: true
            - package_name: RColorBrewer
              package_path: RColorBrewer_1.1-3.tar.gz
              bioconductor: true
            - package_name: viridisLite
              package_path: viridisLite_0.4.3.tar.gz
              bioconductor: true
            - package_name: cpp11
              package_path: cpp11_0.5.3.tar.gz
              bioconductor: true
        - package_name: ica
          package_path: ica_1.0-3.tar.gz
          bioconductor: true
        - package_name: irlba
          package_path: irlba_2.3.7.tar.gz
          bioconductor: true
        - package_name: jsonlite
          package_path: jsonlite_2.0.0.tar.gz
          bioconductor: true
        - package_name: lifecycle
          package_path: lifecycle_1.0.5.tar.gz
          bioconductor: true
          dependencies:
            - package_name: cli
              package_path: cli_3.6.5.tar.gz
              bioconductor: true
        - package_name: KernSmooth
          package_path: KernSmooth_2.23-26.tar.gz
          bioconductor: true
        - package_name: igraph
          package_path: igraph_2.2.2.tar.gz
          bioconductor: true
          dependencies:
            - package_name: cli
              package_path: cli_3.6.5.tar.gz
              bioconductor: true
            - package_name: lifecycle
              package_path: lifecycle_1.0.5.tar.gz
              bioconductor: true
              dependencies:
                - package_name: cli
                  package_path: cli_3.6.5.tar.gz
                  bioconductor: true
            - package_name: magrittr
              package_path: magrittr_2.0.4.tar.gz
              bioconductor: true
            - package_name: pkgconfig
              package_path: pkgconfig_2.0.3.tar.gz
              bioconductor: true
            - package_name: rlang
              package_path: rlang_1.1.7.tar.gz
              bioconductor: true
            - package_name: vctrs
              package_path: vctrs_0.7.1.tar.gz
              bioconductor: true
              dependencies:
                - package_name: rlang
                  package_path: rlang_1.1.7.tar.gz
                  bioconductor: true
                - package_name: glue
                  package_path: glue_1.8.0.tar.gz
                  bioconductor: true
                - package_name: cli
                  package_path: cli_3.6.5.tar.gz
                  bioconductor: true
                - package_name: lifecycle
                  package_path: lifecycle_1.0.5.tar.gz
                  bioconductor: true
                  dependencies:
                    - package_name: cli
                      package_path: cli_3.6.5.tar.gz
                      bioconductor: true
            - package_name: cpp11
              package_path: cpp11_0.5.3.tar.gz
              bioconductor: true
            - package_name: glue
              package_path: glue_1.8.0.tar.gz
              bioconductor: true
          external_dependencies:
            apt:
                - name: libxml2-dev
                  dependencies:
                    - name: libc-dev
                      optional: true
                    - name: libc6
                    - name: libcrypt1
                    - name: gcc-14-base
                    - name: rpcsvc-proto
                    - name: libcrypt-dev
                    - name: linux-libc-dev
                    - name: libc-dev-bin
                    - name: libgcc-s1
                    - name: libstdc++6
                    - name: zlib1g
                    - name: liblzma5
                    - name: libc6-dev
                    - name: libicu74
                    - name: icu-devtools
                    - name: libxml2
                    - name: libicu-dev
                - name: libglpk-dev
                  dependencies:
                    - name: liblapack.so.3
                      optional: true
                    - name: liblapack.so
                      optional: true
                    - name: libblas.so.3
                      optional: true
                    - name: libblas.so
                      optional: true
                    - name: libc-dev
                      optional: true
                    - name: libgcc-s1
                    - name: libc6
                    - name: gcc-14-base
                    - name: libgfortran5
                    - name: libcrypt1
                    - name: libgmp10
                    - name: libmpfr6
                    - name: libopenblas0-serial
                    - name: libopenblas0-pthread
                    - name: libopenblas0-openmp
                    - name: libblis4-serial
                    - name: libblis4-pthread
                    - name: libblis4-openmp
                    - name: libblas3
                    - name: libatlas3-base
                    - name: liblapack3
                    - name: libgomp1
                    - name: rpcsvc-proto
                    - name: libcrypt-dev
                    - name: linux-libc-dev
                    - name: libc-dev-bin
                    - name: libstdc++6
                    - name: libopenblas-serial-dev
                    - name: libopenblas-pthread-dev
                    - name: libopenblas-openmp-dev
                    - name: libblis-serial-dev
                    - name: libblis-pthread-dev
                    - name: libblis-openmp-dev
                    - name: libblas-dev
                    - name: libatlas-base-dev
                    - name: liblapack-dev
                    - name: libsuitesparseconfig7
                    - name: libumfpack6
                    - name: libparu0
                    - name: libcolamd3
                    - name: libamd3
                    - name: libspex2
                    - name: libcholmod5
                    - name: libspqr4
                    - name: librbio4
                    - name: libsuitesparse-mongoose3
                    - name: libldl3
                    - name: libbtf2
                    - name: libklu2
                    - name: libcxsparse4
                    - name: libccolamd3
                    - name: libcamd3
                    - name: libc6-dev
                    - name: libgmpxx4ldbl
                    - name: zlib1g
                    - name: libltdl7
                    - name: libsuitesparse-dev
                    - name: zlib1g-dev
                    - name: libgmp-dev
                    - name: libglpk40
        - package_name: httr
          package_path: httr_1.4.8.tar.gz
          bioconductor: true
          dependencies:
            - package_name: mime
              package_path: mime_0.13.tar.gz
              bioconductor: true
            - package_name: R6
              package_path: R6_2.6.1.tar.gz
              bioconductor: true
            - package_name: jsonlite
              package_path: jsonlite_2.0.0.tar.gz
              bioconductor: true
            - package_name: sys
              package_path: sys_3.4.3.tar.gz
              bioconductor: true
            - package_name: askpass
              package_path: askpass_1.2.1.tar.gz
              bioconductor: true
              dependencies:
                - package_name: sys
                  package_path: sys_3.4.3.tar.gz
                  bioconductor: true
            - package_name: curl
              package_path: curl_7.0.0.tar.gz
              bioconductor: true
              external_dependencies:
                apt:
                    - name: libcurl4-openssl-dev
                      dependencies:
                        - name: libhogweed6
                          optional: true
                        - name: libgnutls30
                          optional: true
                        - name: libnettle8
                          optional: true
                        - name: libc6
                        - name: libdb5.3t64
                        - name: libffi8
                        - name: libsasl2-modules-db
                        - name: libtasn1-6
                        - name: libp11-kit0
                        - name: libkeyutils1
                        - name: gcc-14-base
                        - name: libnettle8t64
                        - name: libgmp10
                        - name: libhogweed6t64
                        - name: libssl3t64
                        - name: libsasl2-2
                        - name: libunistring5
                        - name: libidn2-0
                        - name: libgnutls30t64
                        - name: libkrb5support0
                        - name: libk5crypto3
                        - name: libcom-err2
                        - name: libkrb5-3
                        - name: libgcc-s1
                        - name: zlib1g
                        - name: libzstd1
                        - name: libgssapi-krb5-2
                        - name: libssh-4
                        - name: librtmp1
                        - name: libpsl5t64
                        - name: libnghttp2-14
                        - name: libldap2
                        - name: libbrotli1
                        - name: libcurl4t64
            - package_name: openssl
              package_path: openssl_2.3.5.tar.gz
              bioconductor: true
              dependencies:
                - package_name: sys
                  package_path: sys_3.4.3.tar.gz
                  bioconductor: true
                - package_name: askpass
                  package_path: askpass_1.2.1.tar.gz
                  bioconductor: true
                  dependencies:
                    - package_name: sys
                      package_path: sys_3.4.3.tar.gz
                      bioconductor: true
              external_dependencies:
                apt:
                    - name: libssl-dev
                      dependencies:
                        - name: gcc-14-base
                        - name: libc6
                        - name: libgcc-s1
                        - name: libssl3t64
        - package_name: matrixStats
          package_path: matrixStats_1.5.0.tar.gz
          bioconductor: true
        - package_name: lmtest
          package_path: lmtest_0.9-40.tar.gz
          bioconductor: true
          dependencies:
            - package_name: lattice
              package_path: lattice_0.22-9.tar.gz
              bioconductor: true
            - package_name: zoo
              package_path: zoo_1.8-15.tar.gz
              bioconductor: true
              dependencies:
                - package_name: lattice
                  package_path: lattice_0.22-9.tar.gz
                  bioconductor: true
        - package_name: pbapply
          package_path: pbapply_1.7-4.tar.gz
          bioconductor: true
        - package_name: patchwork
          package_path: patchwork_1.3.2.tar.gz
          bioconductor: true
          dependencies:
            - package_name: gtable
              package_path: gtable_0.3.6.tar.gz
              bioconductor: true
              dependencies:
                - package_name: rlang
                  package_path: rlang_1.1.7.tar.gz
                  bioconductor: true
            - package_name: ggplot2
              package_path: ggplot2_4.0.2.tar.gz
              bioconductor: true
              dependencies:
                - package_name: lifecycle
                  package_path: lifecycle_1.0.5.tar.gz
                  bioconductor: true
                  dependencies:
                    - package_name: cli
                      package_path: cli_3.6.5.tar.gz
                      bioconductor: true
                - package_name: gtable
                  package_path: gtable_0.3.6.tar.gz
                  bioconductor: true
                  dependencies:
                    - package_name: glue
                      package_path: glue_1.8.0.tar.gz
                      bioconductor: true
                    - package_name: rlang
                      package_path: rlang_1.1.7.tar.gz
                      bioconductor: true
                    - package_name: lifecycle
                      package_path: lifecycle_1.0.5.tar.gz
                      bioconductor: true
                - package_name: rlang
                  package_path: rlang_1.1.7.tar.gz
                  bioconductor: true
                - package_name: S7
                  package_path: S7_0.2.1.tar.gz
                  bioconductor: true
                - package_name: isoband
                  package_path: isoband_0.3.0.tar.gz
                  bioconductor: true
                  dependencies:
                    - package_name: rlang
                      package_path: rlang_1.1.7.tar.gz
                      bioconductor: true
                    - package_name: cpp11
                      package_path: cpp11_0.5.3.tar.gz
                      bioconductor: true
                - package_name: glue
                  package_path: glue_1.8.0.tar.gz
                  bioconductor: true
                - package_name: cpp11
                  package_path: cpp11_0.5.3.tar.gz
                  bioconductor: true
                - package_name: vctrs
                  package_path: vctrs_0.7.1.tar.gz
                  bioconductor: true
                  dependencies:
                    - package_name: rlang
                      package_path: rlang_1.1.7.tar.gz
                      bioconductor: true
                    - package_name: cli
                      package_path: cli_3.6.5.tar.gz
                      bioconductor: true
                    - package_name: lifecycle
                      package_path: lifecycle_1.0.5.tar.gz
                      bioconductor: true
                      dependencies:
                        - package_name: cli
                          package_path: cli_3.6.5.tar.gz
                          bioconductor: true
                    - package_name: glue
                      package_path: glue_1.8.0.tar.gz
                      bioconductor: true
                - package_name: farver
                  package_path: farver_2.1.2.tar.gz
                  bioconductor: true
                - package_name: RColorBrewer
                  package_path: RColorBrewer_1.1-3.tar.gz
                  bioconductor: true
                - package_name: labeling
                  package_path: labeling_0.4.3.tar.gz
                  bioconductor: true
                - package_name: scales
                  package_path: scales_1.4.0.tar.gz
                  bioconductor: true
                  dependencies:
                    - package_name: glue
                      package_path: glue_1.8.0.tar.gz
                      bioconductor: true
                    - package_name: lifecycle
                      package_path: lifecycle_1.0.5.tar.gz
                      bioconductor: true
                      dependencies:
                        - package_name: cli
                          package_path: cli_3.6.5.tar.gz
                          bioconductor: true
                    - package_name: cli
                      package_path: cli_3.6.5.tar.gz
                      bioconductor: true
                    - package_name: RColorBrewer
                      package_path: RColorBrewer_1.1-3.tar.gz
                      bioconductor: true
                    - package_name: rlang
                      package_path: rlang_1.1.7.tar.gz
                      bioconductor: true
                    - package_name: farver
                      package_path: farver_2.1.2.tar.gz
                      bioconductor: true
                    - package_name: viridisLite
                      package_path: viridisLite_0.4.3.tar.gz
                      bioconductor: true
                    - package_name: labeling
                      package_path: labeling_0.4.3.tar.gz
                      bioconductor: true
                - package_name: R6
                  package_path: R6_2.6.1.tar.gz
                  bioconductor: true
                - package_name: viridisLite
                  package_path: viridisLite_0.4.3.tar.gz
                  bioconductor: true
            - package_name: rlang
              package_path: rlang_1.1.7.tar.gz
              bioconductor: true
            - package_name: cli
              package_path: cli_3.6.5.tar.gz
              bioconductor: true
            - package_name: farver
              package_path: farver_2.1.2.tar.gz
              bioconductor: true
            - package_name: isoband
              package_path: isoband_0.3.0.tar.gz
              bioconductor: true
              dependencies:
                - package_name: rlang
                  package_path: rlang_1.1.7.tar.gz
                  bioconductor: true
                - package_name: cpp11
                  package_path: cpp11_0.5.3.tar.gz
                  bioconductor: true
            - package_name: lifecycle
              package_path: lifecycle_1.0.5.tar.gz
              bioconductor: true
              dependencies:
                - package_name: cli
                  package_path: cli_3.6.5.tar.gz
                  bioconductor: true
            - package_name: S7
              package_path: S7_0.2.1.tar.gz
              bioconductor: true
            - package_name: scales
              package_path: scales_1.4.0.tar.gz
              bioconductor: true
              dependencies:
                - package_name: glue
                  package_path: glue_1.8.0.tar.gz
                  bioconductor: true
                - package_name: lifecycle
                  package_path: lifecycle_1.0.5.tar.gz
                  bioconductor: true
                  dependencies:
                    - package_name: cli
                      package_path: cli_3.6.5.tar.gz
                      bioconductor: true
                - package_name: cli
                  package_path: cli_3.6.5.tar.gz
                  bioconductor: true
                - package_name: RColorBrewer
                  package_path: RColorBrewer_1.1-3.tar.gz
                  bioconductor: true
                - package_name: rlang
                  package_path: rlang_1.1.7.tar.gz
                  bioconductor: true
                - package_name: farver
                  package_path: farver_2.1.2.tar.gz
                  bioconductor: true
                - package_name: viridisLite
                  package_path: viridisLite_0.4.3.tar.gz
                  bioconductor: true
                - package_name: labeling
                  package_path: labeling_0.4.3.tar.gz
                  bioconductor: true
            - package_name: vctrs
              package_path: vctrs_0.7.1.tar.gz
              bioconductor: true
              dependencies:
                - package_name: rlang
                  package_path: rlang_1.1.7.tar.gz
                  bioconductor: true
                - package_name: glue
                  package_path: glue_1.8.0.tar.gz
                  bioconductor: true
                - package_name: cli
                  package_path: cli_3.6.5.tar.gz
                  bioconductor: true
                - package_name: lifecycle
                  package_path: lifecycle_1.0.5.tar.gz
                  bioconductor: true
                  dependencies:
                    - package_name: cli
                      package_path: cli_3.6.5.tar.gz
                      bioconductor: true
            - package_name: withr
              package_path: withr_3.0.2.tar.gz
              bioconductor: true
            - package_name: glue
              package_path: glue_1.8.0.tar.gz
              bioconductor: true
            - package_name: cpp11
              package_path: cpp11_0.5.3.tar.gz
              bioconductor: true
            - package_name: labeling
              package_path: labeling_0.4.3.tar.gz
              bioconductor: true
            - package_name: R6
              package_path: R6_2.6.1.tar.gz
              bioconductor: true
            - package_name: RColorBrewer
              package_path: RColorBrewer_1.1-3.tar.gz
              bioconductor: true
            - package_name: viridisLite
              package_path: viridisLite_0.4.3.tar.gz
              bioconductor: true
        - package_name: progressr
          package_path: progressr_0.18.0.tar.gz
          bioconductor: true
          dependencies:
            - package_name: digest
              package_path: digest_0.6.39.tar.gz
              bioconductor: true
        - package_name: RANN
          package_path: RANN_2.6.2.tar.gz
          bioconductor: true
        - package_name: RColorBrewer
          package_path: RColorBrewer_1.1-3.tar.gz
          bioconductor: true
        - package_name: Rcpp
          package_path: Rcpp_1.1.1.tar.gz
          bioconductor: true
        - package_name: png
          package_path: png_0.1-8.tar.gz
          bioconductor: true
          external_dependencies:
            apt:
                - name: libpng-dev
                  dependencies:
                    - name: libc-dev
                      optional: true
                    - name: libc6
                    - name: libcrypt1
                    - name: gcc-14-base
                    - name: rpcsvc-proto
                    - name: libcrypt-dev
                    - name: linux-libc-dev
                    - name: libc-dev-bin
                    - name: libgcc-s1
                    - name: libc6-dev
                    - name: zlib1g
                    - name: zlib1g-dev
                    - name: libpng16-16t64
        - package_name: RcppAnnoy
          package_path: RcppAnnoy_0.0.23.tar.gz
          bioconductor: true
          dependencies:
            - package_name: Rcpp
              package_path: Rcpp_1.1.1.tar.gz
              bioconductor: true
        - package_name: RcppHNSW
          package_path: RcppHNSW_0.6.0.tar.gz
          bioconductor: true
          dependencies:
            - package_name: Rcpp
              package_path: Rcpp_1.1.1.tar.gz
              bioconductor: true
        - package_name: rlang
          package_path: rlang_1.1.7.tar.gz
          bioconductor: true
        - package_name: reticulate
          package_path: reticulate_1.45.0.tar.gz
          bioconductor: true
          dependencies:
            - package_name: Rcpp
              package_path: Rcpp_1.1.1.tar.gz
              bioconductor: true
            - package_name: jsonlite
              package_path: jsonlite_2.0.0.tar.gz
              bioconductor: true
            - package_name: png
              package_path: png_0.1-8.tar.gz
              bioconductor: true
              external_dependencies:
                apt:
                    - name: libpng-dev
                      dependencies:
                        - name: libc-dev
                          optional: true
                        - name: libc6
                        - name: libcrypt1
                        - name: gcc-14-base
                        - name: rpcsvc-proto
                        - name: libcrypt-dev
                        - name: linux-libc-dev
                        - name: libc-dev-bin
                        - name: libgcc-s1
                        - name: libc6-dev
                        - name: zlib1g
                        - name: zlib1g-dev
                        - name: libpng16-16t64
            - package_name: rappdirs
              package_path: rappdirs_0.3.4.tar.gz
              bioconductor: true
            - package_name: rlang
              package_path: rlang_1.1.7.tar.gz
              bioconductor: true
            - package_name: withr
              package_path: withr_3.0.2.tar.gz
              bioconductor: true
            - package_name: rprojroot
              package_path: rprojroot_2.1.1.tar.gz
              bioconductor: true
            - package_name: RcppTOML
              package_path: RcppTOML_0.2.3.tar.gz
              bioconductor: true
              dependencies:
                - package_name: Rcpp
                  package_path: Rcpp_1.1.1.tar.gz
                  bioconductor: true
            - package_name: here
              package_path: here_1.0.2.tar.gz
              bioconductor: true
              dependencies:
                - package_name: rprojroot
                  package_path: rprojroot_2.1.1.tar.gz
                  bioconductor: true
        - package_name: ROCR
          package_path: ROCR_1.0-12.tar.gz
          bioconductor: true
          dependencies:
            - package_name: KernSmooth
              package_path: KernSmooth_2.23-26.tar.gz
              bioconductor: true
            - package_name: bitops
              package_path: bitops_1.0-9.tar.gz
              bioconductor: true
            - package_name: gtools
              package_path: gtools_3.9.5.tar.gz
              bioconductor: true
            - package_name: caTools
              package_path: caTools_1.18.3.tar.gz
              bioconductor: true
              dependencies:
                - package_name: bitops
                  package_path: bitops_1.0-9.tar.gz
                  bioconductor: true
            - package_name: gplots
              package_path: gplots_3.3.0.tar.gz
              bioconductor: true
              dependencies:
                - package_name: KernSmooth
                  package_path: KernSmooth_2.23-26.tar.gz
                  bioconductor: true
                - package_name: caTools
                  package_path: caTools_1.18.3.tar.gz
                  bioconductor: true
                  dependencies:
                    - package_name: bitops
                      package_path: bitops_1.0-9.tar.gz
                      bioconductor: true
                - package_name: bitops
                  package_path: bitops_1.0-9.tar.gz
                  bioconductor: true
                - package_name: gtools
                  package_path: gtools_3.9.5.tar.gz
                  bioconductor: true
        - package_name: miniUI
          package_path: miniUI_0.1.2.tar.gz
          bioconductor: true
          dependencies:
            - package_name: digest
              package_path: digest_0.6.39.tar.gz
              bioconductor: true
            - package_name: base64enc
              package_path: base64enc_0.1-6.tar.gz
              bioconductor: true
            - package_name: fastmap
              package_path: fastmap_1.2.0.tar.gz
              bioconductor: true
            - package_name: rlang
              package_path: rlang_1.1.7.tar.gz
              bioconductor: true
            - package_name: htmltools
              package_path: htmltools_0.5.9.tar.gz
              bioconductor: true
              dependencies:
                - package_name: digest
                  package_path: digest_0.6.39.tar.gz
                  bioconductor: true
                - package_name: rlang
                  package_path: rlang_1.1.7.tar.gz
                  bioconductor: true
                - package_name: base64enc
                  package_path: base64enc_0.1-6.tar.gz
                  bioconductor: true
                - package_name: fastmap
                  package_path: fastmap_1.2.0.tar.gz
                  bioconductor: true
            - package_name: cli
              package_path: cli_3.6.5.tar.gz
              bioconductor: true
            - package_name: commonmark
              package_path: commonmark_2.0.0.tar.gz
              bioconductor: true
            - package_name: cachem
              package_path: cachem_1.1.0.tar.gz
              bioconductor: true
              dependencies:
                - package_name: rlang
                  package_path: rlang_1.1.7.tar.gz
                  bioconductor: true
                - package_name: fastmap
                  package_path: fastmap_1.2.0.tar.gz
                  bioconductor: true
            - package_name: glue
              package_path: glue_1.8.0.tar.gz
              bioconductor: true
            - package_name: fontawesome
              package_path: fontawesome_0.5.3.tar.gz
              bioconductor: true
              dependencies:
                - package_name: digest
                  package_path: digest_0.6.39.tar.gz
                  bioconductor: true
                - package_name: fastmap
                  package_path: fastmap_1.2.0.tar.gz
                  bioconductor: true
                - package_name: htmltools
                  package_path: htmltools_0.5.9.tar.gz
                  bioconductor: true
                  dependencies:
                    - package_name: digest
                      package_path: digest_0.6.39.tar.gz
                      bioconductor: true
                    - package_name: rlang
                      package_path: rlang_1.1.7.tar.gz
                      bioconductor: true
                    - package_name: base64enc
                      package_path: base64enc_0.1-6.tar.gz
                      bioconductor: true
                    - package_name: fastmap
                      package_path: fastmap_1.2.0.tar.gz
                      bioconductor: true
                - package_name: base64enc
                  package_path: base64enc_0.1-6.tar.gz
                  bioconductor: true
                - package_name: rlang
                  package_path: rlang_1.1.7.tar.gz
                  bioconductor: true
            - package_name: jsonlite
              package_path: jsonlite_2.0.0.tar.gz
              bioconductor: true
            - package_name: later
              package_path: later_1.4.8.tar.gz
              bioconductor: true
              dependencies:
                - package_name: rlang
                  package_path: rlang_1.1.7.tar.gz
                  bioconductor: true
                - package_name: Rcpp
                  package_path: Rcpp_1.1.1.tar.gz
                  bioconductor: true
            - package_name: lifecycle
              package_path: lifecycle_1.0.5.tar.gz
              bioconductor: true
              dependencies:
                - package_name: cli
                  package_path: cli_3.6.5.tar.gz
                  bioconductor: true
            - package_name: mime
              package_path: mime_0.13.tar.gz
              bioconductor: true
            - package_name: otel
              package_path: otel_0.2.0.tar.gz
              bioconductor: true
            - package_name: bslib
              package_path: bslib_0.10.0.tar.gz
              bioconductor: true
              dependencies:
                - package_name: fastmap
                  package_path: fastmap_1.2.0.tar.gz
                  bioconductor: true
                - package_name: htmltools
                  package_path: htmltools_0.5.9.tar.gz
                  bioconductor: true
                  dependencies:
                    - package_name: digest
                      package_path: digest_0.6.39.tar.gz
                      bioconductor: true
                    - package_name: rlang
                      package_path: rlang_1.1.7.tar.gz
                      bioconductor: true
                    - package_name: base64enc
                      package_path: base64enc_0.1-6.tar.gz
                      bioconductor: true
                    - package_name: fastmap
                      package_path: fastmap_1.2.0.tar.gz
                      bioconductor: true
                - package_name: base64enc
                  package_path: base64enc_0.1-6.tar.gz
                  bioconductor: true
                - package_name: jsonlite
                  package_path: jsonlite_2.0.0.tar.gz
                  bioconductor: true
                - package_name: lifecycle
                  package_path: lifecycle_1.0.5.tar.gz
                  bioconductor: true
                  dependencies:
                    - package_name: cli
                      package_path: cli_3.6.5.tar.gz
                      bioconductor: true
                - package_name: cachem
                  package_path: cachem_1.1.0.tar.gz
                  bioconductor: true
                  dependencies:
                    - package_name: rlang
                      package_path: rlang_1.1.7.tar.gz
                      bioconductor: true
                    - package_name: fastmap
                      package_path: fastmap_1.2.0.tar.gz
                      bioconductor: true
                - package_name: mime
                  package_path: mime_0.13.tar.gz
                  bioconductor: true
                - package_name: rlang
                  package_path: rlang_1.1.7.tar.gz
                  bioconductor: true
                - package_name: digest
                  package_path: digest_0.6.39.tar.gz
                  bioconductor: true
                - package_name: cli
                  package_path: cli_3.6.5.tar.gz
                  bioconductor: true
                - package_name: memoise
                  package_path: memoise_2.0.1.tar.gz
                  bioconductor: true
                  dependencies:
                    - package_name: cachem
                      package_path: cachem_1.1.0.tar.gz
                      bioconductor: true
                      dependencies:
                        - package_name: rlang
                          package_path: rlang_1.1.7.tar.gz
                          bioconductor: true
                        - package_name: fastmap
                          package_path: fastmap_1.2.0.tar.gz
                          bioconductor: true
                    - package_name: rlang
                      package_path: rlang_1.1.7.tar.gz
                      bioconductor: true
                    - package_name: fastmap
                      package_path: fastmap_1.2.0.tar.gz
                      bioconductor: true
                - package_name: R6
                  package_path: R6_2.6.1.tar.gz
                  bioconductor: true
                - package_name: jquerylib
                  package_path: jquerylib_0.1.4.tar.gz
                  bioconductor: true
                  dependencies:
                    - package_name: base64enc
                      package_path: base64enc_0.1-6.tar.gz
                      bioconductor: true
                    - package_name: fastmap
                      package_path: fastmap_1.2.0.tar.gz
                      bioconductor: true
                    - package_name: rlang
                      package_path: rlang_1.1.7.tar.gz
                      bioconductor: true
                    - package_name: htmltools
                      package_path: htmltools_0.5.9.tar.gz
                      bioconductor: true
                      dependencies:
                        - package_name: digest
                          package_path: digest_0.6.39.tar.gz
                          bioconductor: true
                        - package_name: rlang
                          package_path: rlang_1.1.7.tar.gz
                          bioconductor: true
                        - package_name: base64enc
                          package_path: base64enc_0.1-6.tar.gz
                          bioconductor: true
                        - package_name: fastmap
                          package_path: fastmap_1.2.0.tar.gz
                          bioconductor: true
                    - package_name: digest
                      package_path: digest_0.6.39.tar.gz
                      bioconductor: true
                - package_name: sass
                  package_path: sass_0.4.10.tar.gz
                  bioconductor: true
                  dependencies:
                    - package_name: R6
                      package_path: R6_2.6.1.tar.gz
                      bioconductor: true
                    - package_name: htmltools
                      package_path: htmltools_0.5.9.tar.gz
                      bioconductor: true
                      dependencies:
                        - package_name: digest
                          package_path: digest_0.6.39.tar.gz
                          bioconductor: true
                        - package_name: rlang
                          package_path: rlang_1.1.7.tar.gz
                          bioconductor: true
                        - package_name: base64enc
                          package_path: base64enc_0.1-6.tar.gz
                          bioconductor: true
                        - package_name: fastmap
                          package_path: fastmap_1.2.0.tar.gz
                          bioconductor: true
                    - package_name: rlang
                      package_path: rlang_1.1.7.tar.gz
                      bioconductor: true
                    - package_name: base64enc
                      package_path: base64enc_0.1-6.tar.gz
                      bioconductor: true
                    - package_name: digest
                      package_path: digest_0.6.39.tar.gz
                      bioconductor: true
                    - package_name: fastmap
                      package_path: fastmap_1.2.0.tar.gz
                      bioconductor: true
                    - package_name: rappdirs
                      package_path: rappdirs_0.3.4.tar.gz
                      bioconductor: true
                  external_dependencies:
                    apt:
                        - name: make
                          dependencies:
                            - name: gcc-14-base
                            - name: libc6
                            - name: libgcc-s1
                - package_name: rappdirs
                  package_path: rappdirs_0.3.4.tar.gz
                  bioconductor: true
            - package_name: R6
              package_path: R6_2.6.1.tar.gz
              bioconductor: true
            - package_name: sourcetools
              package_path: sourcetools_0.1.7-1.tar.gz
              bioconductor: true
            - package_name: withr
              package_path: withr_3.0.2.tar.gz
              bioconductor: true
            - package_name: httpuv
              package_path: httpuv_1.6.16.tar.gz
              bioconductor: true
              dependencies:
                - package_name: Rcpp
                  package_path: Rcpp_1.1.1.tar.gz
                  bioconductor: true
                - package_name: later
                  package_path: later_1.4.8.tar.gz
                  bioconductor: true
                  dependencies:
                    - package_name: rlang
                      package_path: rlang_1.1.7.tar.gz
                      bioconductor: true
                    - package_name: Rcpp
                      package_path: Rcpp_1.1.1.tar.gz
                      bioconductor: true
                - package_name: rlang
                  package_path: rlang_1.1.7.tar.gz
                  bioconductor: true
                - package_name: fastmap
                  package_path: fastmap_1.2.0.tar.gz
                  bioconductor: true
                - package_name: lifecycle
                  package_path: lifecycle_1.0.5.tar.gz
                  bioconductor: true
                  dependencies:
                    - package_name: cli
                      package_path: cli_3.6.5.tar.gz
                      bioconductor: true
                - package_name: magrittr
                  package_path: magrittr_2.0.4.tar.gz
                  bioconductor: true
                - package_name: R6
                  package_path: R6_2.6.1.tar.gz
                  bioconductor: true
                - package_name: cli
                  package_path: cli_3.6.5.tar.gz
                  bioconductor: true
                - package_name: otel
                  package_path: otel_0.2.0.tar.gz
                  bioconductor: true
                - package_name: promises
                  package_path: promises_1.5.0.tar.gz
                  bioconductor: true
                  dependencies:
                    - package_name: magrittr
                      package_path: magrittr_2.0.4.tar.gz
                      bioconductor: true
                    - package_name: otel
                      package_path: otel_0.2.0.tar.gz
                      bioconductor: true
                    - package_name: R6
                      package_path: R6_2.6.1.tar.gz
                      bioconductor: true
                    - package_name: rlang
                      package_path: rlang_1.1.7.tar.gz
                      bioconductor: true
                    - package_name: Rcpp
                      package_path: Rcpp_1.1.1.tar.gz
                      bioconductor: true
                    - package_name: later
                      package_path: later_1.4.8.tar.gz
                      bioconductor: true
                      dependencies:
                        - package_name: rlang
                          package_path: rlang_1.1.7.tar.gz
                          bioconductor: true
                        - package_name: Rcpp
                          package_path: Rcpp_1.1.1.tar.gz
                          bioconductor: true
                    - package_name: lifecycle
                      package_path: lifecycle_1.0.5.tar.gz
                      bioconductor: true
                      dependencies:
                        - package_name: cli
                          package_path: cli_3.6.5.tar.gz
                          bioconductor: true
                    - package_name: fastmap
                      package_path: fastmap_1.2.0.tar.gz
                      bioconductor: true
                    - package_name: cli
                      package_path: cli_3.6.5.tar.gz
                      bioconductor: true
              external_dependencies:
                apt:
                    - name: make
                      dependencies:
                        - name: gcc-14-base
                        - name: libc6
                        - name: libgcc-s1
                    - name: zlib1g-dev
                      dependencies:
                        - name: libc-dev
                          optional: true
                        - name: gcc-14-base
                        - name: libc6
                        - name: libcrypt1
                        - name: libgcc-s1
                        - name: rpcsvc-proto
                        - name: libcrypt-dev
                        - name: linux-libc-dev
                        - name: libc-dev-bin
                        - name: libc6-dev
                        - name: zlib1g
            - package_name: jquerylib
              package_path: jquerylib_0.1.4.tar.gz
              bioconductor: true
              dependencies:
                - package_name: fastmap
                  package_path: fastmap_1.2.0.tar.gz
                  bioconductor: true
                - package_name: base64enc
                  package_path: base64enc_0.1-6.tar.gz
                  bioconductor: true
                - package_name: rlang
                  package_path: rlang_1.1.7.tar.gz
                  bioconductor: true
                - package_name: htmltools
                  package_path: htmltools_0.5.9.tar.gz
                  bioconductor: true
                  dependencies:
                    - package_name: digest
                      package_path: digest_0.6.39.tar.gz
                      bioconductor: true
                    - package_name: rlang
                      package_path: rlang_1.1.7.tar.gz
                      bioconductor: true
                    - package_name: base64enc
                      package_path: base64enc_0.1-6.tar.gz
                      bioconductor: true
                    - package_name: fastmap
                      package_path: fastmap_1.2.0.tar.gz
                      bioconductor: true
                - package_name: digest
                  package_path: digest_0.6.39.tar.gz
                  bioconductor: true
            - package_name: memoise
              package_path: memoise_2.0.1.tar.gz
              bioconductor: true
              dependencies:
                - package_name: cachem
                  package_path: cachem_1.1.0.tar.gz
                  bioconductor: true
                  dependencies:
                    - package_name: rlang
                      package_path: rlang_1.1.7.tar.gz
                      bioconductor: true
                    - package_name: fastmap
                      package_path: fastmap_1.2.0.tar.gz
                      bioconductor: true
                - package_name: fastmap
                  package_path: fastmap_1.2.0.tar.gz
                  bioconductor: true
                - package_name: rlang
                  package_path: rlang_1.1.7.tar.gz
                  bioconductor: true
            - package_name: sass
              package_path: sass_0.4.10.tar.gz
              bioconductor: true
              dependencies:
                - package_name: R6
                  package_path: R6_2.6.1.tar.gz
                  bioconductor: true
                - package_name: htmltools
                  package_path: htmltools_0.5.9.tar.gz
                  bioconductor: true
                  dependencies:
                    - package_name: digest
                      package_path: digest_0.6.39.tar.gz
                      bioconductor: true
                    - package_name: rlang
                      package_path: rlang_1.1.7.tar.gz
                      bioconductor: true
                    - package_name: base64enc
                      package_path: base64enc_0.1-6.tar.gz
                      bioconductor: true
                    - package_name: fastmap
                      package_path: fastmap_1.2.0.tar.gz
                      bioconductor: true
                - package_name: rlang
                  package_path: rlang_1.1.7.tar.gz
                  bioconductor: true
                - package_name: base64enc
                  package_path: base64enc_0.1-6.tar.gz
                  bioconductor: true
                - package_name: digest
                  package_path: digest_0.6.39.tar.gz
                  bioconductor: true
                - package_name: fastmap
                  package_path: fastmap_1.2.0.tar.gz
                  bioconductor: true
                - package_name: rappdirs
                  package_path: rappdirs_0.3.4.tar.gz
                  bioconductor: true
              external_dependencies:
                apt:
                    - name: make
                      dependencies:
                        - name: gcc-14-base
                        - name: libc6
                        - name: libgcc-s1
            - package_name: Rcpp
              package_path: Rcpp_1.1.1.tar.gz
              bioconductor: true
            - package_name: magrittr
              package_path: magrittr_2.0.4.tar.gz
              bioconductor: true
            - package_name: shiny
              package_path: shiny_1.13.0.tar.gz
              bioconductor: true
              dependencies:
                - package_name: cli
                  package_path: cli_3.6.5.tar.gz
                  bioconductor: true
                - package_name: fastmap
                  package_path: fastmap_1.2.0.tar.gz
                  bioconductor: true
                - package_name: commonmark
                  package_path: commonmark_2.0.0.tar.gz
                  bioconductor: true
                - package_name: glue
                  package_path: glue_1.8.0.tar.gz
                  bioconductor: true
                - package_name: cachem
                  package_path: cachem_1.1.0.tar.gz
                  bioconductor: true
                  dependencies:
                    - package_name: rlang
                      package_path: rlang_1.1.7.tar.gz
                      bioconductor: true
                    - package_name: fastmap
                      package_path: fastmap_1.2.0.tar.gz
                      bioconductor: true
                - package_name: htmltools
                  package_path: htmltools_0.5.9.tar.gz
                  bioconductor: true
                  dependencies:
                    - package_name: digest
                      package_path: digest_0.6.39.tar.gz
                      bioconductor: true
                    - package_name: rlang
                      package_path: rlang_1.1.7.tar.gz
                      bioconductor: true
                    - package_name: base64enc
                      package_path: base64enc_0.1-6.tar.gz
                      bioconductor: true
                    - package_name: fastmap
                      package_path: fastmap_1.2.0.tar.gz
                      bioconductor: true
                - package_name: jsonlite
                  package_path: jsonlite_2.0.0.tar.gz
                  bioconductor: true
                - package_name: fontawesome
                  package_path: fontawesome_0.5.3.tar.gz
                  bioconductor: true
                  dependencies:
                    - package_name: digest
                      package_path: digest_0.6.39.tar.gz
                      bioconductor: true
                    - package_name: fastmap
                      package_path: fastmap_1.2.0.tar.gz
                      bioconductor: true
                    - package_name: htmltools
                      package_path: htmltools_0.5.9.tar.gz
                      bioconductor: true
                      dependencies:
                        - package_name: digest
                          package_path: digest_0.6.39.tar.gz
                          bioconductor: true
                        - package_name: rlang
                          package_path: rlang_1.1.7.tar.gz
                          bioconductor: true
                        - package_name: base64enc
                          package_path: base64enc_0.1-6.tar.gz
                          bioconductor: true
                        - package_name: fastmap
                          package_path: fastmap_1.2.0.tar.gz
                          bioconductor: true
                    - package_name: rlang
                      package_path: rlang_1.1.7.tar.gz
                      bioconductor: true
                    - package_name: base64enc
                      package_path: base64enc_0.1-6.tar.gz
                      bioconductor: true
                - package_name: lifecycle
                  package_path: lifecycle_1.0.5.tar.gz
                  bioconductor: true
                  dependencies:
                    - package_name: cli
                      package_path: cli_3.6.5.tar.gz
                      bioconductor: true
                - package_name: mime
                  package_path: mime_0.13.tar.gz
                  bioconductor: true
                - package_name: later
                  package_path: later_1.4.8.tar.gz
                  bioconductor: true
                  dependencies:
                    - package_name: rlang
                      package_path: rlang_1.1.7.tar.gz
                      bioconductor: true
                    - package_name: Rcpp
                      package_path: Rcpp_1.1.1.tar.gz
                      bioconductor: true
                - package_name: R6
                  package_path: R6_2.6.1.tar.gz
                  bioconductor: true
                - package_name: rlang
                  package_path: rlang_1.1.7.tar.gz
                  bioconductor: true
                - package_name: withr
                  package_path: withr_3.0.2.tar.gz
                  bioconductor: true
                - package_name: base64enc
                  package_path: base64enc_0.1-6.tar.gz
                  bioconductor: true
                - package_name: jquerylib
                  package_path: jquerylib_0.1.4.tar.gz
                  bioconductor: true
                  dependencies:
                    - package_name: fastmap
                      package_path: fastmap_1.2.0.tar.gz
                      bioconductor: true
                    - package_name: base64enc
                      package_path: base64enc_0.1-6.tar.gz
                      bioconductor: true
                    - package_name: rlang
                      package_path: rlang_1.1.7.tar.gz
                      bioconductor: true
                    - package_name: htmltools
                      package_path: htmltools_0.5.9.tar.gz
                      bioconductor: true
                      dependencies:
                        - package_name: digest
                          package_path: digest_0.6.39.tar.gz
                          bioconductor: true
                        - package_name: rlang
                          package_path: rlang_1.1.7.tar.gz
                          bioconductor: true
                        - package_name: base64enc
                          package_path: base64enc_0.1-6.tar.gz
                          bioconductor: true
                        - package_name: fastmap
                          package_path: fastmap_1.2.0.tar.gz
                          bioconductor: true
                    - package_name: digest
                      package_path: digest_0.6.39.tar.gz
                      bioconductor: true
                - package_name: memoise
                  package_path: memoise_2.0.1.tar.gz
                  bioconductor: true
                  dependencies:
                    - package_name: cachem
                      package_path: cachem_1.1.0.tar.gz
                      bioconductor: true
                      dependencies:
                        - package_name: rlang
                          package_path: rlang_1.1.7.tar.gz
                          bioconductor: true
                        - package_name: fastmap
                          package_path: fastmap_1.2.0.tar.gz
                          bioconductor: true
                    - package_name: fastmap
                      package_path: fastmap_1.2.0.tar.gz
                      bioconductor: true
                    - package_name: rlang
                      package_path: rlang_1.1.7.tar.gz
                      bioconductor: true
                - package_name: sass
                  package_path: sass_0.4.10.tar.gz
                  bioconductor: true
                  dependencies:
                    - package_name: R6
                      package_path: R6_2.6.1.tar.gz
                      bioconductor: true
                    - package_name: htmltools
                      package_path: htmltools_0.5.9.tar.gz
                      bioconductor: true
                      dependencies:
                        - package_name: digest
                          package_path: digest_0.6.39.tar.gz
                          bioconductor: true
                        - package_name: rlang
                          package_path: rlang_1.1.7.tar.gz
                          bioconductor: true
                        - package_name: base64enc
                          package_path: base64enc_0.1-6.tar.gz
                          bioconductor: true
                        - package_name: fastmap
                          package_path: fastmap_1.2.0.tar.gz
                          bioconductor: true
                    - package_name: rlang
                      package_path: rlang_1.1.7.tar.gz
                      bioconductor: true
                    - package_name: base64enc
                      package_path: base64enc_0.1-6.tar.gz
                      bioconductor: true
                    - package_name: digest
                      package_path: digest_0.6.39.tar.gz
                      bioconductor: true
                    - package_name: fastmap
                      package_path: fastmap_1.2.0.tar.gz
                      bioconductor: true
                    - package_name: rappdirs
                      package_path: rappdirs_0.3.4.tar.gz
                      bioconductor: true
                  external_dependencies:
                    apt:
                        - name: make
                          dependencies:
                            - name: gcc-14-base
                            - name: libc6
                            - name: libgcc-s1
                - package_name: digest
                  package_path: digest_0.6.39.tar.gz
                  bioconductor: true
                - package_name: Rcpp
                  package_path: Rcpp_1.1.1.tar.gz
                  bioconductor: true
                - package_name: magrittr
                  package_path: magrittr_2.0.4.tar.gz
                  bioconductor: true
                - package_name: sourcetools
                  package_path: sourcetools_0.1.7-1.tar.gz
                  bioconductor: true
                - package_name: rappdirs
                  package_path: rappdirs_0.3.4.tar.gz
                  bioconductor: true
                - package_name: httpuv
                  package_path: httpuv_1.6.16.tar.gz
                  bioconductor: true
                  dependencies:
                    - package_name: Rcpp
                      package_path: Rcpp_1.1.1.tar.gz
                      bioconductor: true
                    - package_name: R6
                      package_path: R6_2.6.1.tar.gz
                      bioconductor: true
                    - package_name: rlang
                      package_path: rlang_1.1.7.tar.gz
                      bioconductor: true
                    - package_name: fastmap
                      package_path: fastmap_1.2.0.tar.gz
                      bioconductor: true
                    - package_name: lifecycle
                      package_path: lifecycle_1.0.5.tar.gz
                      bioconductor: true
                      dependencies:
                        - package_name: cli
                          package_path: cli_3.6.5.tar.gz
                          bioconductor: true
                    - package_name: magrittr
                      package_path: magrittr_2.0.4.tar.gz
                      bioconductor: true
                    - package_name: cli
                      package_path: cli_3.6.5.tar.gz
                      bioconductor: true
                    - package_name: otel
                      package_path: otel_0.2.0.tar.gz
                      bioconductor: true
                    - package_name: promises
                      package_path: promises_1.5.0.tar.gz
                      bioconductor: true
                      dependencies:
                        - package_name: magrittr
                          package_path: magrittr_2.0.4.tar.gz
                          bioconductor: true
                        - package_name: later
                          package_path: later_1.4.8.tar.gz
                          bioconductor: true
                          dependencies:
                            - package_name: rlang
                              package_path: rlang_1.1.7.tar.gz
                              bioconductor: true
                            - package_name: Rcpp
                              package_path: Rcpp_1.1.1.tar.gz
                              bioconductor: true
                        - package_name: R6
                          package_path: R6_2.6.1.tar.gz
                          bioconductor: true
                        - package_name: rlang
                          package_path: rlang_1.1.7.tar.gz
                          bioconductor: true
                        - package_name: Rcpp
                          package_path: Rcpp_1.1.1.tar.gz
                          bioconductor: true
                        - package_name: fastmap
                          package_path: fastmap_1.2.0.tar.gz
                          bioconductor: true
                        - package_name: lifecycle
                          package_path: lifecycle_1.0.5.tar.gz
                          bioconductor: true
                          dependencies:
                            - package_name: cli
                              package_path: cli_3.6.5.tar.gz
                              bioconductor: true
                        - package_name: cli
                          package_path: cli_3.6.5.tar.gz
                          bioconductor: true
                        - package_name: otel
                          package_path: otel_0.2.0.tar.gz
                          bioconductor: true
                    - package_name: later
                      package_path: later_1.4.8.tar.gz
                      bioconductor: true
                      dependencies:
                        - package_name: Rcpp
                          package_path: Rcpp_1.1.1.tar.gz
                          bioconductor: true
                        - package_name: rlang
                          package_path: rlang_1.1.7.tar.gz
                          bioconductor: true
                  external_dependencies:
                    apt:
                        - name: make
                          dependencies:
                            - name: gcc-14-base
                            - name: libc6
                            - name: libgcc-s1
                        - name: zlib1g-dev
                          dependencies:
                            - name: libc-dev
                              optional: true
                            - name: gcc-14-base
                            - name: libc6
                            - name: libcrypt1
                            - name: libgcc-s1
                            - name: rpcsvc-proto
                            - name: libcrypt-dev
                            - name: linux-libc-dev
                            - name: libc-dev-bin
                            - name: libc6-dev
                            - name: zlib1g
                - package_name: xtable
                  package_path: xtable_1.8-8.tar.gz
                  bioconductor: true
            - package_name: rappdirs
              package_path: rappdirs_0.3.4.tar.gz
              bioconductor: true
            - package_name: promises
              package_path: promises_1.5.0.tar.gz
              bioconductor: true
              dependencies:
                - package_name: magrittr
                  package_path: magrittr_2.0.4.tar.gz
                  bioconductor: true
                - package_name: later
                  package_path: later_1.4.8.tar.gz
                  bioconductor: true
                  dependencies:
                    - package_name: rlang
                      package_path: rlang_1.1.7.tar.gz
                      bioconductor: true
                    - package_name: Rcpp
                      package_path: Rcpp_1.1.1.tar.gz
                      bioconductor: true
                - package_name: R6
                  package_path: R6_2.6.1.tar.gz
                  bioconductor: true
                - package_name: rlang
                  package_path: rlang_1.1.7.tar.gz
                  bioconductor: true
                - package_name: Rcpp
                  package_path: Rcpp_1.1.1.tar.gz
                  bioconductor: true
                - package_name: cli
                  package_path: cli_3.6.5.tar.gz
                  bioconductor: true
                - package_name: lifecycle
                  package_path: lifecycle_1.0.5.tar.gz
                  bioconductor: true
                  dependencies:
                    - package_name: cli
                      package_path: cli_3.6.5.tar.gz
                      bioconductor: true
                - package_name: otel
                  package_path: otel_0.2.0.tar.gz
                  bioconductor: true
                - package_name: fastmap
                  package_path: fastmap_1.2.0.tar.gz
                  bioconductor: true
            - package_name: xtable
              package_path: xtable_1.8-8.tar.gz
              bioconductor: true
            - package_name: fs
              package_path: fs_1.6.7.tar.gz
              bioconductor: true
              external_dependencies:
                apt:
                    - name: make
                      dependencies:
                        - name: gcc-14-base
                        - name: libc6
                        - name: libgcc-s1
        - package_name: scales
          package_path: scales_1.4.0.tar.gz
          bioconductor: true
          dependencies:
            - package_name: glue
              package_path: glue_1.8.0.tar.gz
              bioconductor: true
            - package_name: lifecycle
              package_path: lifecycle_1.0.5.tar.gz
              bioconductor: true
              dependencies:
                - package_name: cli
                  package_path: cli_3.6.5.tar.gz
                  bioconductor: true
            - package_name: cli
              package_path: cli_3.6.5.tar.gz
              bioconductor: true
            - package_name: RColorBrewer
              package_path: RColorBrewer_1.1-3.tar.gz
              bioconductor: true
            - package_name: rlang
              package_path: rlang_1.1.7.tar.gz
              bioconductor: true
            - package_name: farver
              package_path: farver_2.1.2.tar.gz
              bioconductor: true
            - package_name: viridisLite
              package_path: viridisLite_0.4.3.tar.gz
              bioconductor: true
            - package_name: labeling
              package_path: labeling_0.4.3.tar.gz
              bioconductor: true
        - package_name: RSpectra
          package_path: RSpectra_0.16-2.tar.gz
          bioconductor: true
          dependencies:
            - package_name: RcppEigen
              package_path: RcppEigen_0.3.4.0.2.tar.gz
              bioconductor: true
              dependencies:
                - package_name: Rcpp
                  package_path: Rcpp_1.1.1.tar.gz
                  bioconductor: true
            - package_name: Rcpp
              package_path: Rcpp_1.1.1.tar.gz
              bioconductor: true
        - package_name: plotly
          package_path: plotly_4.12.0.tar.gz
          bioconductor: true
          dependencies:
            - package_name: httr
              package_path: httr_1.4.8.tar.gz
              bioconductor: true
              dependencies:
                - package_name: mime
                  package_path: mime_0.13.tar.gz
                  bioconductor: true
                - package_name: R6
                  package_path: R6_2.6.1.tar.gz
                  bioconductor: true
                - package_name: jsonlite
                  package_path: jsonlite_2.0.0.tar.gz
                  bioconductor: true
                - package_name: sys
                  package_path: sys_3.4.3.tar.gz
                  bioconductor: true
                - package_name: askpass
                  package_path: askpass_1.2.1.tar.gz
                  bioconductor: true
                  dependencies:
                    - package_name: sys
                      package_path: sys_3.4.3.tar.gz
                      bioconductor: true
                - package_name: curl
                  package_path: curl_7.0.0.tar.gz
                  bioconductor: true
                  external_dependencies:
                    apt:
                        - name: libcurl4-openssl-dev
                          dependencies:
                            - name: libhogweed6
                              optional: true
                            - name: libgnutls30
                              optional: true
                            - name: libnettle8
                              optional: true
                            - name: libc6
                            - name: libdb5.3t64
                            - name: libffi8
                            - name: libsasl2-modules-db
                            - name: libtasn1-6
                            - name: libp11-kit0
                            - name: libkeyutils1
                            - name: gcc-14-base
                            - name: libnettle8t64
                            - name: libgmp10
                            - name: libhogweed6t64
                            - name: libssl3t64
                            - name: libsasl2-2
                            - name: libunistring5
                            - name: libidn2-0
                            - name: libgnutls30t64
                            - name: libkrb5support0
                            - name: libk5crypto3
                            - name: libcom-err2
                            - name: libkrb5-3
                            - name: libgcc-s1
                            - name: zlib1g
                            - name: libzstd1
                            - name: libgssapi-krb5-2
                            - name: libssh-4
                            - name: librtmp1
                            - name: libpsl5t64
                            - name: libnghttp2-14
                            - name: libldap2
                            - name: libbrotli1
                            - name: libcurl4t64
                - package_name: openssl
                  package_path: openssl_2.3.5.tar.gz
                  bioconductor: true
                  dependencies:
                    - package_name: sys
                      package_path: sys_3.4.3.tar.gz
                      bioconductor: true
                    - package_name: askpass
                      package_path: askpass_1.2.1.tar.gz
                      bioconductor: true
                      dependencies:
                        - package_name: sys
                          package_path: sys_3.4.3.tar.gz
                          bioconductor: true
                  external_dependencies:
                    apt:
                        - name: libssl-dev
                          dependencies:
                            - name: gcc-14-base
                            - name: libc6
                            - name: libgcc-s1
                            - name: libssl3t64
            - package_name: ggplot2
              package_path: ggplot2_4.0.2.tar.gz
              bioconductor: true
              dependencies:
                - package_name: lifecycle
                  package_path: lifecycle_1.0.5.tar.gz
                  bioconductor: true
                  dependencies:
                    - package_name: cli
                      package_path: cli_3.6.5.tar.gz
                      bioconductor: true
                - package_name: gtable
                  package_path: gtable_0.3.6.tar.gz
                  bioconductor: true
                  dependencies:
                    - package_name: glue
                      package_path: glue_1.8.0.tar.gz
                      bioconductor: true
                    - package_name: rlang
                      package_path: rlang_1.1.7.tar.gz
                      bioconductor: true
                    - package_name: lifecycle
                      package_path: lifecycle_1.0.5.tar.gz
                      bioconductor: true
                - package_name: rlang
                  package_path: rlang_1.1.7.tar.gz
                  bioconductor: true
                - package_name: S7
                  package_path: S7_0.2.1.tar.gz
                  bioconductor: true
                - package_name: isoband
                  package_path: isoband_0.3.0.tar.gz
                  bioconductor: true
                  dependencies:
                    - package_name: rlang
                      package_path: rlang_1.1.7.tar.gz
                      bioconductor: true
                    - package_name: cpp11
                      package_path: cpp11_0.5.3.tar.gz
                      bioconductor: true
                - package_name: glue
                  package_path: glue_1.8.0.tar.gz
                  bioconductor: true
                - package_name: cpp11
                  package_path: cpp11_0.5.3.tar.gz
                  bioconductor: true
                - package_name: vctrs
                  package_path: vctrs_0.7.1.tar.gz
                  bioconductor: true
                  dependencies:
                    - package_name: rlang
                      package_path: rlang_1.1.7.tar.gz
                      bioconductor: true
                    - package_name: cli
                      package_path: cli_3.6.5.tar.gz
                      bioconductor: true
                    - package_name: lifecycle
                      package_path: lifecycle_1.0.5.tar.gz
                      bioconductor: true
                      dependencies:
                        - package_name: cli
                          package_path: cli_3.6.5.tar.gz
                          bioconductor: true
                    - package_name: glue
                      package_path: glue_1.8.0.tar.gz
                      bioconductor: true
                - package_name: farver
                  package_path: farver_2.1.2.tar.gz
                  bioconductor: true
                - package_name: RColorBrewer
                  package_path: RColorBrewer_1.1-3.tar.gz
                  bioconductor: true
                - package_name: labeling
                  package_path: labeling_0.4.3.tar.gz
                  bioconductor: true
                - package_name: scales
                  package_path: scales_1.4.0.tar.gz
                  bioconductor: true
                  dependencies:
                    - package_name: glue
                      package_path: glue_1.8.0.tar.gz
                      bioconductor: true
                    - package_name: lifecycle
                      package_path: lifecycle_1.0.5.tar.gz
                      bioconductor: true
                      dependencies:
                        - package_name: cli
                          package_path: cli_3.6.5.tar.gz
                          bioconductor: true
                    - package_name: cli
                      package_path: cli_3.6.5.tar.gz
                      bioconductor: true
                    - package_name: RColorBrewer
                      package_path: RColorBrewer_1.1-3.tar.gz
                      bioconductor: true
                    - package_name: rlang
                      package_path: rlang_1.1.7.tar.gz
                      bioconductor: true
                    - package_name: farver
                      package_path: farver_2.1.2.tar.gz
                      bioconductor: true
                    - package_name: viridisLite
                      package_path: viridisLite_0.4.3.tar.gz
                      bioconductor: true
                    - package_name: labeling
                      package_path: labeling_0.4.3.tar.gz
                      bioconductor: true
                - package_name: R6
                  package_path: R6_2.6.1.tar.gz
                  bioconductor: true
                - package_name: viridisLite
                  package_path: viridisLite_0.4.3.tar.gz
                  bioconductor: true
            - package_name: magrittr
              package_path: magrittr_2.0.4.tar.gz
              bioconductor: true
            - package_name: jsonlite
              package_path: jsonlite_2.0.0.tar.gz
              bioconductor: true
            - package_name: viridisLite
              package_path: viridisLite_0.4.3.tar.gz
              bioconductor: true
            - package_name: digest
              package_path: digest_0.6.39.tar.gz
              bioconductor: true
            - package_name: scales
              package_path: scales_1.4.0.tar.gz
              bioconductor: true
              dependencies:
                - package_name: glue
                  package_path: glue_1.8.0.tar.gz
                  bioconductor: true
                - package_name: lifecycle
                  package_path: lifecycle_1.0.5.tar.gz
                  bioconductor: true
                  dependencies:
                    - package_name: cli
                      package_path: cli_3.6.5.tar.gz
                      bioconductor: true
                - package_name: cli
                  package_path: cli_3.6.5.tar.gz
                  bioconductor: true
                - package_name: RColorBrewer
                  package_path: RColorBrewer_1.1-3.tar.gz
                  bioconductor: true
                - package_name: rlang
                  package_path: rlang_1.1.7.tar.gz
                  bioconductor: true
                - package_name: farver
                  package_path: farver_2.1.2.tar.gz
                  bioconductor: true
                - package_name: viridisLite
                  package_path: viridisLite_0.4.3.tar.gz
                  bioconductor: true
                - package_name: labeling
                  package_path: labeling_0.4.3.tar.gz
                  bioconductor: true
            - package_name: base64enc
              package_path: base64enc_0.1-6.tar.gz
              bioconductor: true
            - package_name: RColorBrewer
              package_path: RColorBrewer_1.1-3.tar.gz
              bioconductor: true
            - package_name: htmltools
              package_path: htmltools_0.5.9.tar.gz
              bioconductor: true
              dependencies:
                - package_name: digest
                  package_path: digest_0.6.39.tar.gz
                  bioconductor: true
                - package_name: rlang
                  package_path: rlang_1.1.7.tar.gz
                  bioconductor: true
                - package_name: fastmap
                  package_path: fastmap_1.2.0.tar.gz
                  bioconductor: true
                - package_name: base64enc
                  package_path: base64enc_0.1-6.tar.gz
                  bioconductor: true
            - package_name: vctrs
              package_path: vctrs_0.7.1.tar.gz
              bioconductor: true
              dependencies:
                - package_name: rlang
                  package_path: rlang_1.1.7.tar.gz
                  bioconductor: true
                - package_name: glue
                  package_path: glue_1.8.0.tar.gz
                  bioconductor: true
                - package_name: cli
                  package_path: cli_3.6.5.tar.gz
                  bioconductor: true
                - package_name: lifecycle
                  package_path: lifecycle_1.0.5.tar.gz
                  bioconductor: true
                  dependencies:
                    - package_name: cli
                      package_path: cli_3.6.5.tar.gz
                      bioconductor: true
            - package_name: tibble
              package_path: tibble_3.3.1.tar.gz
              bioconductor: true
              dependencies:
                - package_name: magrittr
                  package_path: magrittr_2.0.4.tar.gz
                  bioconductor: true
                - package_name: rlang
                  package_path: rlang_1.1.7.tar.gz
                  bioconductor: true
                - package_name: pkgconfig
                  package_path: pkgconfig_2.0.3.tar.gz
                  bioconductor: true
                - package_name: lifecycle
                  package_path: lifecycle_1.0.5.tar.gz
                  bioconductor: true
                - package_name: glue
                  package_path: glue_1.8.0.tar.gz
                  bioconductor: true
                - package_name: pillar
                  package_path: pillar_1.11.1.tar.gz
                  bioconductor: true
                  dependencies:
                    - package_name: rlang
                      package_path: rlang_1.1.7.tar.gz
                      bioconductor: true
                    - package_name: glue
                      package_path: glue_1.8.0.tar.gz
                      bioconductor: true
                    - package_name: utf8
                      package_path: utf8_1.2.6.tar.gz
                      bioconductor: true
                    - package_name: lifecycle
                      package_path: lifecycle_1.0.5.tar.gz
                      bioconductor: true
                      dependencies:
                        - package_name: rlang
                          package_path: rlang_1.1.7.tar.gz
                          bioconductor: true
                        - package_name: cli
                          package_path: cli_3.6.5.tar.gz
                          bioconductor: true
                - package_name: utf8
                  package_path: utf8_1.2.6.tar.gz
                  bioconductor: true
                - package_name: vctrs
                  package_path: vctrs_0.7.1.tar.gz
                  bioconductor: true
                  dependencies:
                    - package_name: rlang
                      package_path: rlang_1.1.7.tar.gz
                      bioconductor: true
                    - package_name: glue
                      package_path: glue_1.8.0.tar.gz
                      bioconductor: true
                    - package_name: cli
                      package_path: cli_3.6.5.tar.gz
                      bioconductor: true
                    - package_name: lifecycle
                      package_path: lifecycle_1.0.5.tar.gz
                      bioconductor: true
                      dependencies:
                        - package_name: cli
                          package_path: cli_3.6.5.tar.gz
                          bioconductor: true
            - package_name: lazyeval
              package_path: lazyeval_0.2.2.tar.gz
              bioconductor: true
            - package_name: rlang
              package_path: rlang_1.1.7.tar.gz
              bioconductor: true
            - package_name: dplyr
              package_path: dplyr_1.2.0.tar.gz
              bioconductor: true
              dependencies:
                - package_name: lifecycle
                  package_path: lifecycle_1.0.5.tar.gz
                  bioconductor: true
                  dependencies:
                    - package_name: cli
                      package_path: cli_3.6.5.tar.gz
                      bioconductor: true
                - package_name: generics
                  package_path: generics_0.1.4.tar.gz
                  bioconductor: true
                - package_name: cli
                  package_path: cli_3.6.5.tar.gz
                  bioconductor: true
                - package_name: pillar
                  package_path: pillar_1.11.1.tar.gz
                  bioconductor: true
                  dependencies:
                    - package_name: rlang
                      package_path: rlang_1.1.7.tar.gz
                      bioconductor: true
                    - package_name: glue
                      package_path: glue_1.8.0.tar.gz
                      bioconductor: true
                    - package_name: utf8
                      package_path: utf8_1.2.6.tar.gz
                      bioconductor: true
                    - package_name: lifecycle
                      package_path: lifecycle_1.0.5.tar.gz
                      bioconductor: true
                      dependencies:
                        - package_name: rlang
                          package_path: rlang_1.1.7.tar.gz
                          bioconductor: true
                        - package_name: cli
                          package_path: cli_3.6.5.tar.gz
                          bioconductor: true
                - package_name: R6
                  package_path: R6_2.6.1.tar.gz
                  bioconductor: true
                - package_name: rlang
                  package_path: rlang_1.1.7.tar.gz
                  bioconductor: true
                - package_name: tibble
                  package_path: tibble_3.3.1.tar.gz
                  bioconductor: true
                  dependencies:
                    - package_name: magrittr
                      package_path: magrittr_2.0.4.tar.gz
                      bioconductor: true
                    - package_name: rlang
                      package_path: rlang_1.1.7.tar.gz
                      bioconductor: true
                    - package_name: pkgconfig
                      package_path: pkgconfig_2.0.3.tar.gz
                      bioconductor: true
                    - package_name: lifecycle
                      package_path: lifecycle_1.0.5.tar.gz
                      bioconductor: true
                    - package_name: glue
                      package_path: glue_1.8.0.tar.gz
                      bioconductor: true
                    - package_name: pillar
                      package_path: pillar_1.11.1.tar.gz
                      bioconductor: true
                      dependencies:
                        - package_name: rlang
                          package_path: rlang_1.1.7.tar.gz
                          bioconductor: true
                        - package_name: glue
                          package_path: glue_1.8.0.tar.gz
                          bioconductor: true
                        - package_name: utf8
                          package_path: utf8_1.2.6.tar.gz
                          bioconductor: true
                        - package_name: lifecycle
                          package_path: lifecycle_1.0.5.tar.gz
                          bioconductor: true
                          dependencies:
                            - package_name: rlang
                              package_path: rlang_1.1.7.tar.gz
                              bioconductor: true
                            - package_name: cli
                              package_path: cli_3.6.5.tar.gz
                              bioconductor: true
                    - package_name: utf8
                      package_path: utf8_1.2.6.tar.gz
                      bioconductor: true
                    - package_name: vctrs
                      package_path: vctrs_0.7.1.tar.gz
                      bioconductor: true
                      dependencies:
                        - package_name: rlang
                          package_path: rlang_1.1.7.tar.gz
                          bioconductor: true
                        - package_name: glue
                          package_path: glue_1.8.0.tar.gz
                          bioconductor: true
                        - package_name: cli
                          package_path: cli_3.6.5.tar.gz
                          bioconductor: true
                        - package_name: lifecycle
                          package_path: lifecycle_1.0.5.tar.gz
                          bioconductor: true
                          dependencies:
                            - package_name: cli
                              package_path: cli_3.6.5.tar.gz
                              bioconductor: true
                - package_name: glue
                  package_path: glue_1.8.0.tar.gz
                  bioconductor: true
                - package_name: magrittr
                  package_path: magrittr_2.0.4.tar.gz
                  bioconductor: true
                - package_name: vctrs
                  package_path: vctrs_0.7.1.tar.gz
                  bioconductor: true
                  dependencies:
                    - package_name: rlang
                      package_path: rlang_1.1.7.tar.gz
                      bioconductor: true
                    - package_name: glue
                      package_path: glue_1.8.0.tar.gz
                      bioconductor: true
                    - package_name: cli
                      package_path: cli_3.6.5.tar.gz
                      bioconductor: true
                    - package_name: lifecycle
                      package_path: lifecycle_1.0.5.tar.gz
                      bioconductor: true
                      dependencies:
                        - package_name: cli
                          package_path: cli_3.6.5.tar.gz
                          bioconductor: true
                - package_name: utf8
                  package_path: utf8_1.2.6.tar.gz
                  bioconductor: true
                - package_name: pkgconfig
                  package_path: pkgconfig_2.0.3.tar.gz
                  bioconductor: true
                - package_name: withr
                  package_path: withr_3.0.2.tar.gz
                  bioconductor: true
            - package_name: purrr
              package_path: purrr_1.2.1.tar.gz
              bioconductor: true
              dependencies:
                - package_name: rlang
                  package_path: rlang_1.1.7.tar.gz
                  bioconductor: true
                - package_name: vctrs
                  package_path: vctrs_0.7.1.tar.gz
                  bioconductor: true
                  dependencies:
                    - package_name: rlang
                      package_path: rlang_1.1.7.tar.gz
                      bioconductor: true
                    - package_name: glue
                      package_path: glue_1.8.0.tar.gz
                      bioconductor: true
                    - package_name: cli
                      package_path: cli_3.6.5.tar.gz
                      bioconductor: true
                    - package_name: lifecycle
                      package_path: lifecycle_1.0.5.tar.gz
                      bioconductor: true
                      dependencies:
                        - package_name: cli
                          package_path: cli_3.6.5.tar.gz
                          bioconductor: true
                - package_name: lifecycle
                  package_path: lifecycle_1.0.5.tar.gz
                  bioconductor: true
                  dependencies:
                    - package_name: cli
                      package_path: cli_3.6.5.tar.gz
                      bioconductor: true
                - package_name: glue
                  package_path: glue_1.8.0.tar.gz
                  bioconductor: true
                - package_name: cli
                  package_path: cli_3.6.5.tar.gz
                  bioconductor: true
                - package_name: magrittr
                  package_path: magrittr_2.0.4.tar.gz
                  bioconductor: true
            - package_name: data.table
              package_path: data.table_1.18.2.1.tar.gz
              bioconductor: true
            - package_name: tidyr
              package_path: tidyr_1.3.2.tar.gz
              bioconductor: true
              dependencies:
                - package_name: lifecycle
                  package_path: lifecycle_1.0.5.tar.gz
                  bioconductor: true
                  dependencies:
                    - package_name: cli
                      package_path: cli_3.6.5.tar.gz
                      bioconductor: true
                - package_name: magrittr
                  package_path: magrittr_2.0.4.tar.gz
                  bioconductor: true
                - package_name: glue
                  package_path: glue_1.8.0.tar.gz
                  bioconductor: true
                - package_name: rlang
                  package_path: rlang_1.1.7.tar.gz
                  bioconductor: true
                - package_name: stringr
                  package_path: stringr_1.6.0.tar.gz
                  bioconductor: true
                  dependencies:
                    - package_name: magrittr
                      package_path: magrittr_2.0.4.tar.gz
                      bioconductor: true
                    - package_name: glue
                      package_path: glue_1.8.0.tar.gz
                      bioconductor: true
                    - package_name: lifecycle
                      package_path: lifecycle_1.0.5.tar.gz
                      bioconductor: true
                      dependencies:
                        - package_name: cli
                          package_path: cli_3.6.5.tar.gz
                          bioconductor: true
                    - package_name: stringi
                      package_path: stringi_1.8.7.tar.gz
                      bioconductor: true
                      external_dependencies:
                        apt:
                            - name: libicu-dev
                              dependencies:
                                - name: libc-dev
                                  optional: true
                                - name: libc6
                                - name: libcrypt1
                                - name: gcc-14-base
                                - name: rpcsvc-proto
                                - name: libcrypt-dev
                                - name: linux-libc-dev
                                - name: libc-dev-bin
                                - name: libgcc-s1
                                - name: libstdc++6
                                - name: libc6-dev
                                - name: libicu74
                                - name: icu-devtools
                    - package_name: rlang
                      package_path: rlang_1.1.7.tar.gz
                      bioconductor: true
                - package_name: tibble
                  package_path: tibble_3.3.1.tar.gz
                  bioconductor: true
                  dependencies:
                    - package_name: magrittr
                      package_path: magrittr_2.0.4.tar.gz
                      bioconductor: true
                    - package_name: rlang
                      package_path: rlang_1.1.7.tar.gz
                      bioconductor: true
                    - package_name: pkgconfig
                      package_path: pkgconfig_2.0.3.tar.gz
                      bioconductor: true
                    - package_name: lifecycle
                      package_path: lifecycle_1.0.5.tar.gz
                      bioconductor: true
                    - package_name: glue
                      package_path: glue_1.8.0.tar.gz
                      bioconductor: true
                    - package_name: pillar
                      package_path: pillar_1.11.1.tar.gz
                      bioconductor: true
                      dependencies:
                        - package_name: rlang
                          package_path: rlang_1.1.7.tar.gz
                          bioconductor: true
                        - package_name: glue
                          package_path: glue_1.8.0.tar.gz
                          bioconductor: true
                        - package_name: utf8
                          package_path: utf8_1.2.6.tar.gz
                          bioconductor: true
                        - package_name: lifecycle
                          package_path: lifecycle_1.0.5.tar.gz
                          bioconductor: true
                          dependencies:
                            - package_name: rlang
                              package_path: rlang_1.1.7.tar.gz
                              bioconductor: true
                            - package_name: cli
                              package_path: cli_3.6.5.tar.gz
                              bioconductor: true
                    - package_name: utf8
                      package_path: utf8_1.2.6.tar.gz
                      bioconductor: true
                    - package_name: vctrs
                      package_path: vctrs_0.7.1.tar.gz
                      bioconductor: true
                      dependencies:
                        - package_name: rlang
                          package_path: rlang_1.1.7.tar.gz
                          bioconductor: true
                        - package_name: glue
                          package_path: glue_1.8.0.tar.gz
                          bioconductor: true
                        - package_name: cli
                          package_path: cli_3.6.5.tar.gz
                          bioconductor: true
                        - package_name: lifecycle
                          package_path: lifecycle_1.0.5.tar.gz
                          bioconductor: true
                          dependencies:
                            - package_name: cli
                              package_path: cli_3.6.5.tar.gz
                              bioconductor: true
                - package_name: cli
                  package_path: cli_3.6.5.tar.gz
                  bioconductor: true
                - package_name: vctrs
                  package_path: vctrs_0.7.1.tar.gz
                  bioconductor: true
                  dependencies:
                    - package_name: rlang
                      package_path: rlang_1.1.7.tar.gz
                      bioconductor: true
                    - package_name: glue
                      package_path: glue_1.8.0.tar.gz
                      bioconductor: true
                    - package_name: cli
                      package_path: cli_3.6.5.tar.gz
                      bioconductor: true
                    - package_name: lifecycle
                      package_path: lifecycle_1.0.5.tar.gz
                      bioconductor: true
                      dependencies:
                        - package_name: cli
                          package_path: cli_3.6.5.tar.gz
                          bioconductor: true
                - package_name: cpp11
                  package_path: cpp11_0.5.3.tar.gz
                  bioconductor: true
                - package_name: generics
                  package_path: generics_0.1.4.tar.gz
                  bioconductor: true
                - package_name: purrr
                  package_path: purrr_1.2.1.tar.gz
                  bioconductor: true
                  dependencies:
                    - package_name: rlang
                      package_path: rlang_1.1.7.tar.gz
                      bioconductor: true
                    - package_name: vctrs
                      package_path: vctrs_0.7.1.tar.gz
                      bioconductor: true
                      dependencies:
                        - package_name: rlang
                          package_path: rlang_1.1.7.tar.gz
                          bioconductor: true
                        - package_name: glue
                          package_path: glue_1.8.0.tar.gz
                          bioconductor: true
                        - package_name: cli
                          package_path: cli_3.6.5.tar.gz
                          bioconductor: true
                        - package_name: lifecycle
                          package_path: lifecycle_1.0.5.tar.gz
                          bioconductor: true
                          dependencies:
                            - package_name: cli
                              package_path: cli_3.6.5.tar.gz
                              bioconductor: true
                    - package_name: lifecycle
                      package_path: lifecycle_1.0.5.tar.gz
                      bioconductor: true
                      dependencies:
                        - package_name: cli
                          package_path: cli_3.6.5.tar.gz
                          bioconductor: true
                    - package_name: glue
                      package_path: glue_1.8.0.tar.gz
                      bioconductor: true
                    - package_name: cli
                      package_path: cli_3.6.5.tar.gz
                      bioconductor: true
                    - package_name: magrittr
                      package_path: magrittr_2.0.4.tar.gz
                      bioconductor: true
                - package_name: pillar
                  package_path: pillar_1.11.1.tar.gz
                  bioconductor: true
                  dependencies:
                    - package_name: rlang
                      package_path: rlang_1.1.7.tar.gz
                      bioconductor: true
                    - package_name: glue
                      package_path: glue_1.8.0.tar.gz
                      bioconductor: true
                    - package_name: utf8
                      package_path: utf8_1.2.6.tar.gz
                      bioconductor: true
                    - package_name: lifecycle
                      package_path: lifecycle_1.0.5.tar.gz
                      bioconductor: true
                      dependencies:
                        - package_name: rlang
                          package_path: rlang_1.1.7.tar.gz
                          bioconductor: true
                        - package_name: cli
                          package_path: cli_3.6.5.tar.gz
                          bioconductor: true
                - package_name: R6
                  package_path: R6_2.6.1.tar.gz
                  bioconductor: true
                - package_name: stringi
                  package_path: stringi_1.8.7.tar.gz
                  bioconductor: true
                  external_dependencies:
                    apt:
                        - name: libicu-dev
                          dependencies:
                            - name: libc-dev
                              optional: true
                            - name: libc6
                            - name: libcrypt1
                            - name: gcc-14-base
                            - name: rpcsvc-proto
                            - name: libcrypt-dev
                            - name: linux-libc-dev
                            - name: libc-dev-bin
                            - name: libgcc-s1
                            - name: libstdc++6
                            - name: libc6-dev
                            - name: libicu74
                            - name: icu-devtools
                - package_name: pkgconfig
                  package_path: pkgconfig_2.0.3.tar.gz
                  bioconductor: true
                - package_name: withr
                  package_path: withr_3.0.2.tar.gz
                  bioconductor: true
                - package_name: utf8
                  package_path: utf8_1.2.6.tar.gz
                  bioconductor: true
                - package_name: tidyselect
                  package_path: tidyselect_1.2.1.tar.gz
                  bioconductor: true
                  dependencies:
                    - package_name: rlang
                      package_path: rlang_1.1.7.tar.gz
                      bioconductor: true
                    - package_name: vctrs
                      package_path: vctrs_0.7.1.tar.gz
                      bioconductor: true
                      dependencies:
                        - package_name: rlang
                          package_path: rlang_1.1.7.tar.gz
                          bioconductor: true
                        - package_name: glue
                          package_path: glue_1.8.0.tar.gz
                          bioconductor: true
                        - package_name: cli
                          package_path: cli_3.6.5.tar.gz
                          bioconductor: true
                        - package_name: lifecycle
                          package_path: lifecycle_1.0.5.tar.gz
                          bioconductor: true
                          dependencies:
                            - package_name: cli
                              package_path: cli_3.6.5.tar.gz
                              bioconductor: true
                    - package_name: withr
                      package_path: withr_3.0.2.tar.gz
                      bioconductor: true
                    - package_name: glue
                      package_path: glue_1.8.0.tar.gz
                      bioconductor: true
                    - package_name: lifecycle
                      package_path: lifecycle_1.0.5.tar.gz
                      bioconductor: true
                      dependencies:
                        - package_name: cli
                          package_path: cli_3.6.5.tar.gz
                          bioconductor: true
                    - package_name: cli
                      package_path: cli_3.6.5.tar.gz
                      bioconductor: true
                - package_name: dplyr
                  package_path: dplyr_1.2.0.tar.gz
                  bioconductor: true
                  dependencies:
                    - package_name: lifecycle
                      package_path: lifecycle_1.0.5.tar.gz
                      bioconductor: true
                      dependencies:
                        - package_name: cli
                          package_path: cli_3.6.5.tar.gz
                          bioconductor: true
                    - package_name: magrittr
                      package_path: magrittr_2.0.4.tar.gz
                      bioconductor: true
                    - package_name: cli
                      package_path: cli_3.6.5.tar.gz
                      bioconductor: true
                    - package_name: pillar
                      package_path: pillar_1.11.1.tar.gz
                      bioconductor: true
                      dependencies:
                        - package_name: rlang
                          package_path: rlang_1.1.7.tar.gz
                          bioconductor: true
                        - package_name: glue
                          package_path: glue_1.8.0.tar.gz
                          bioconductor: true
                        - package_name: utf8
                          package_path: utf8_1.2.6.tar.gz
                          bioconductor: true
                        - package_name: lifecycle
                          package_path: lifecycle_1.0.5.tar.gz
                          bioconductor: true
                          dependencies:
                            - package_name: rlang
                              package_path: rlang_1.1.7.tar.gz
                              bioconductor: true
                            - package_name: cli
                              package_path: cli_3.6.5.tar.gz
                              bioconductor: true
                    - package_name: glue
                      package_path: glue_1.8.0.tar.gz
                      bioconductor: true
                    - package_name: rlang
                      package_path: rlang_1.1.7.tar.gz
                      bioconductor: true
                    - package_name: tibble
                      package_path: tibble_3.3.1.tar.gz
                      bioconductor: true
                      dependencies:
                        - package_name: magrittr
                          package_path: magrittr_2.0.4.tar.gz
                          bioconductor: true
                        - package_name: rlang
                          package_path: rlang_1.1.7.tar.gz
                          bioconductor: true
                        - package_name: pkgconfig
                          package_path: pkgconfig_2.0.3.tar.gz
                          bioconductor: true
                        - package_name: lifecycle
                          package_path: lifecycle_1.0.5.tar.gz
                          bioconductor: true
                        - package_name: glue
                          package_path: glue_1.8.0.tar.gz
                          bioconductor: true
                        - package_name: pillar
                          package_path: pillar_1.11.1.tar.gz
                          bioconductor: true
                          dependencies:
                            - package_name: rlang
                              package_path: rlang_1.1.7.tar.gz
                              bioconductor: true
                            - package_name: glue
                              package_path: glue_1.8.0.tar.gz
                              bioconductor: true
                            - package_name: utf8
                              package_path: utf8_1.2.6.tar.gz
                              bioconductor: true
                            - package_name: lifecycle
                              package_path: lifecycle_1.0.5.tar.gz
                              bioconductor: true
                              dependencies:
                                - package_name: rlang
                                  package_path: rlang_1.1.7.tar.gz
                                  bioconductor: true
                                - package_name: cli
                                  package_path: cli_3.6.5.tar.gz
                                  bioconductor: true
                        - package_name: utf8
                          package_path: utf8_1.2.6.tar.gz
                          bioconductor: true
                        - package_name: vctrs
                          package_path: vctrs_0.7.1.tar.gz
                          bioconductor: true
                          dependencies:
                            - package_name: rlang
                              package_path: rlang_1.1.7.tar.gz
                              bioconductor: true
                            - package_name: glue
                              package_path: glue_1.8.0.tar.gz
                              bioconductor: true
                            - package_name: cli
                              package_path: cli_3.6.5.tar.gz
                              bioconductor: true
                            - package_name: lifecycle
                              package_path: lifecycle_1.0.5.tar.gz
                              bioconductor: true
                              dependencies:
                                - package_name: cli
                                  package_path: cli_3.6.5.tar.gz
                                  bioconductor: true
                    - package_name: R6
                      package_path: R6_2.6.1.tar.gz
                      bioconductor: true
                    - package_name: generics
                      package_path: generics_0.1.4.tar.gz
                      bioconductor: true
                    - package_name: vctrs
                      package_path: vctrs_0.7.1.tar.gz
                      bioconductor: true
                      dependencies:
                        - package_name: rlang
                          package_path: rlang_1.1.7.tar.gz
                          bioconductor: true
                        - package_name: glue
                          package_path: glue_1.8.0.tar.gz
                          bioconductor: true
                        - package_name: cli
                          package_path: cli_3.6.5.tar.gz
                          bioconductor: true
                        - package_name: lifecycle
                          package_path: lifecycle_1.0.5.tar.gz
                          bioconductor: true
                          dependencies:
                            - package_name: cli
                              package_path: cli_3.6.5.tar.gz
                              bioconductor: true
                    - package_name: utf8
                      package_path: utf8_1.2.6.tar.gz
                      bioconductor: true
                    - package_name: pkgconfig
                      package_path: pkgconfig_2.0.3.tar.gz
                      bioconductor: true
                    - package_name: withr
                      package_path: withr_3.0.2.tar.gz
                      bioconductor: true
                    - package_name: tidyselect
                      package_path: tidyselect_1.2.1.tar.gz
                      bioconductor: true
                      dependencies:
                        - package_name: rlang
                          package_path: rlang_1.1.7.tar.gz
                          bioconductor: true
                        - package_name: vctrs
                          package_path: vctrs_0.7.1.tar.gz
                          bioconductor: true
                          dependencies:
                            - package_name: rlang
                              package_path: rlang_1.1.7.tar.gz
                              bioconductor: true
                            - package_name: glue
                              package_path: glue_1.8.0.tar.gz
                              bioconductor: true
                            - package_name: cli
                              package_path: cli_3.6.5.tar.gz
                              bioconductor: true
                            - package_name: lifecycle
                              package_path: lifecycle_1.0.5.tar.gz
                              bioconductor: true
                              dependencies:
                                - package_name: cli
                                  package_path: cli_3.6.5.tar.gz
                                  bioconductor: true
                        - package_name: withr
                          package_path: withr_3.0.2.tar.gz
                          bioconductor: true
                        - package_name: cli
                          package_path: cli_3.6.5.tar.gz
                          bioconductor: true
                        - package_name: glue
                          package_path: glue_1.8.0.tar.gz
                          bioconductor: true
                        - package_name: lifecycle
                          package_path: lifecycle_1.0.5.tar.gz
                          bioconductor: true
                          dependencies:
                            - package_name: cli
                              package_path: cli_3.6.5.tar.gz
                              bioconductor: true
            - package_name: R6
              package_path: R6_2.6.1.tar.gz
              bioconductor: true
            - package_name: crosstalk
              package_path: crosstalk_1.2.2.tar.gz
              bioconductor: true
              dependencies:
                - package_name: R6
                  package_path: R6_2.6.1.tar.gz
                  bioconductor: true
                - package_name: base64enc
                  package_path: base64enc_0.1-6.tar.gz
                  bioconductor: true
                - package_name: digest
                  package_path: digest_0.6.39.tar.gz
                  bioconductor: true
                - package_name: fastmap
                  package_path: fastmap_1.2.0.tar.gz
                  bioconductor: true
                - package_name: htmltools
                  package_path: htmltools_0.5.9.tar.gz
                  bioconductor: true
                  dependencies:
                    - package_name: digest
                      package_path: digest_0.6.39.tar.gz
                      bioconductor: true
                    - package_name: rlang
                      package_path: rlang_1.1.7.tar.gz
                      bioconductor: true
                    - package_name: base64enc
                      package_path: base64enc_0.1-6.tar.gz
                      bioconductor: true
                    - package_name: fastmap
                      package_path: fastmap_1.2.0.tar.gz
                      bioconductor: true
                - package_name: rlang
                  package_path: rlang_1.1.7.tar.gz
                  bioconductor: true
                - package_name: lazyeval
                  package_path: lazyeval_0.2.2.tar.gz
                  bioconductor: true
                - package_name: jsonlite
                  package_path: jsonlite_2.0.0.tar.gz
                  bioconductor: true
            - package_name: cli
              package_path: cli_3.6.5.tar.gz
              bioconductor: true
            - package_name: generics
              package_path: generics_0.1.4.tar.gz
              bioconductor: true
            - package_name: glue
              package_path: glue_1.8.0.tar.gz
              bioconductor: true
            - package_name: lifecycle
              package_path: lifecycle_1.0.5.tar.gz
              bioconductor: true
              dependencies:
                - package_name: cli
                  package_path: cli_3.6.5.tar.gz
                  bioconductor: true
            - package_name: pillar
              package_path: pillar_1.11.1.tar.gz
              bioconductor: true
              dependencies:
                - package_name: rlang
                  package_path: rlang_1.1.7.tar.gz
                  bioconductor: true
                - package_name: glue
                  package_path: glue_1.8.0.tar.gz
                  bioconductor: true
                - package_name: utf8
                  package_path: utf8_1.2.6.tar.gz
                  bioconductor: true
                - package_name: lifecycle
                  package_path: lifecycle_1.0.5.tar.gz
                  bioconductor: true
                  dependencies:
                    - package_name: rlang
                      package_path: rlang_1.1.7.tar.gz
                      bioconductor: true
                    - package_name: cli
                      package_path: cli_3.6.5.tar.gz
                      bioconductor: true
            - package_name: tidyselect
              package_path: tidyselect_1.2.1.tar.gz
              bioconductor: true
              dependencies:
                - package_name: rlang
                  package_path: rlang_1.1.7.tar.gz
                  bioconductor: true
                - package_name: vctrs
                  package_path: vctrs_0.7.1.tar.gz
                  bioconductor: true
                  dependencies:
                    - package_name: rlang
                      package_path: rlang_1.1.7.tar.gz
                      bioconductor: true
                    - package_name: glue
                      package_path: glue_1.8.0.tar.gz
                      bioconductor: true
                    - package_name: cli
                      package_path: cli_3.6.5.tar.gz
                      bioconductor: true
                    - package_name: lifecycle
                      package_path: lifecycle_1.0.5.tar.gz
                      bioconductor: true
                      dependencies:
                        - package_name: cli
                          package_path: cli_3.6.5.tar.gz
                          bioconductor: true
                - package_name: withr
                  package_path: withr_3.0.2.tar.gz
                  bioconductor: true
                - package_name: glue
                  package_path: glue_1.8.0.tar.gz
                  bioconductor: true
                - package_name: lifecycle
                  package_path: lifecycle_1.0.5.tar.gz
                  bioconductor: true
                  dependencies:
                    - package_name: cli
                      package_path: cli_3.6.5.tar.gz
                      bioconductor: true
                - package_name: cli
                  package_path: cli_3.6.5.tar.gz
                  bioconductor: true
            - package_name: gtable
              package_path: gtable_0.3.6.tar.gz
              bioconductor: true
              dependencies:
                - package_name: rlang
                  package_path: rlang_1.1.7.tar.gz
                  bioconductor: true
            - package_name: isoband
              package_path: isoband_0.3.0.tar.gz
              bioconductor: true
              dependencies:
                - package_name: rlang
                  package_path: rlang_1.1.7.tar.gz
                  bioconductor: true
                - package_name: cpp11
                  package_path: cpp11_0.5.3.tar.gz
                  bioconductor: true
            - package_name: S7
              package_path: S7_0.2.1.tar.gz
              bioconductor: true
            - package_name: promises
              package_path: promises_1.5.0.tar.gz
              bioconductor: true
              dependencies:
                - package_name: magrittr
                  package_path: magrittr_2.0.4.tar.gz
                  bioconductor: true
                - package_name: otel
                  package_path: otel_0.2.0.tar.gz
                  bioconductor: true
                - package_name: R6
                  package_path: R6_2.6.1.tar.gz
                  bioconductor: true
                - package_name: rlang
                  package_path: rlang_1.1.7.tar.gz
                  bioconductor: true
                - package_name: Rcpp
                  package_path: Rcpp_1.1.1.tar.gz
                  bioconductor: true
                - package_name: cli
                  package_path: cli_3.6.5.tar.gz
                  bioconductor: true
                - package_name: fastmap
                  package_path: fastmap_1.2.0.tar.gz
                  bioconductor: true
                - package_name: lifecycle
                  package_path: lifecycle_1.0.5.tar.gz
                  bioconductor: true
                  dependencies:
                    - package_name: cli
                      package_path: cli_3.6.5.tar.gz
                      bioconductor: true
                - package_name: later
                  package_path: later_1.4.8.tar.gz
                  bioconductor: true
                  dependencies:
                    - package_name: rlang
                      package_path: rlang_1.1.7.tar.gz
                      bioconductor: true
                    - package_name: Rcpp
                      package_path: Rcpp_1.1.1.tar.gz
                      bioconductor: true
            - package_name: withr
              package_path: withr_3.0.2.tar.gz
              bioconductor: true
            - package_name: fastmap
              package_path: fastmap_1.2.0.tar.gz
              bioconductor: true
            - package_name: yaml
              package_path: yaml_2.3.12.tar.gz
              bioconductor: true
            - package_name: curl
              package_path: curl_7.0.0.tar.gz
              bioconductor: true
              external_dependencies:
                apt:
                    - name: libcurl4-openssl-dev
                      dependencies:
                        - name: libhogweed6
                          optional: true
                        - name: libgnutls30
                          optional: true
                        - name: libnettle8
                          optional: true
                        - name: libc6
                        - name: libdb5.3t64
                        - name: libffi8
                        - name: libsasl2-modules-db
                        - name: libtasn1-6
                        - name: libp11-kit0
                        - name: libkeyutils1
                        - name: gcc-14-base
                        - name: libnettle8t64
                        - name: libgmp10
                        - name: libhogweed6t64
                        - name: libssl3t64
                        - name: libsasl2-2
                        - name: libunistring5
                        - name: libidn2-0
                        - name: libgnutls30t64
                        - name: libkrb5support0
                        - name: libk5crypto3
                        - name: libcom-err2
                        - name: libkrb5-3
                        - name: libgcc-s1
                        - name: zlib1g
                        - name: libzstd1
                        - name: libgssapi-krb5-2
                        - name: libssh-4
                        - name: librtmp1
                        - name: libpsl5t64
                        - name: libnghttp2-14
                        - name: libldap2
                        - name: libbrotli1
                        - name: libcurl4t64
            - package_name: mime
              package_path: mime_0.13.tar.gz
              bioconductor: true
            - package_name: openssl
              package_path: openssl_2.3.5.tar.gz
              bioconductor: true
              dependencies:
                - package_name: sys
                  package_path: sys_3.4.3.tar.gz
                  bioconductor: true
                - package_name: askpass
                  package_path: askpass_1.2.1.tar.gz
                  bioconductor: true
                  dependencies:
                    - package_name: sys
                      package_path: sys_3.4.3.tar.gz
                      bioconductor: true
              external_dependencies:
                apt:
                    - name: libssl-dev
                      dependencies:
                        - name: gcc-14-base
                        - name: libc6
                        - name: libgcc-s1
                        - name: libssl3t64
            - package_name: later
              package_path: later_1.4.8.tar.gz
              bioconductor: true
              dependencies:
                - package_name: rlang
                  package_path: rlang_1.1.7.tar.gz
                  bioconductor: true
                - package_name: Rcpp
                  package_path: Rcpp_1.1.1.tar.gz
                  bioconductor: true
            - package_name: otel
              package_path: otel_0.2.0.tar.gz
              bioconductor: true
            - package_name: farver
              package_path: farver_2.1.2.tar.gz
              bioconductor: true
            - package_name: labeling
              package_path: labeling_0.4.3.tar.gz
              bioconductor: true
            - package_name: pkgconfig
              package_path: pkgconfig_2.0.3.tar.gz
              bioconductor: true
            - package_name: stringr
              package_path: stringr_1.6.0.tar.gz
              bioconductor: true
              dependencies:
                - package_name: magrittr
                  package_path: magrittr_2.0.4.tar.gz
                  bioconductor: true
                - package_name: glue
                  package_path: glue_1.8.0.tar.gz
                  bioconductor: true
                - package_name: lifecycle
                  package_path: lifecycle_1.0.5.tar.gz
                  bioconductor: true
                  dependencies:
                    - package_name: cli
                      package_path: cli_3.6.5.tar.gz
                      bioconductor: true
                - package_name: stringi
                  package_path: stringi_1.8.7.tar.gz
                  bioconductor: true
                  external_dependencies:
                    apt:
                        - name: libicu-dev
                          dependencies:
                            - name: libc-dev
                              optional: true
                            - name: libc6
                            - name: libcrypt1
                            - name: gcc-14-base
                            - name: rpcsvc-proto
                            - name: libcrypt-dev
                            - name: linux-libc-dev
                            - name: libc-dev-bin
                            - name: libgcc-s1
                            - name: libstdc++6
                            - name: libc6-dev
                            - name: libicu74
                            - name: icu-devtools
                - package_name: rlang
                  package_path: rlang_1.1.7.tar.gz
                  bioconductor: true
            - package_name: cpp11
              package_path: cpp11_0.5.3.tar.gz
              bioconductor: true
            - package_name: evaluate
              package_path: evaluate_1.0.5.tar.gz
              bioconductor: true
            - package_name: highr
              package_path: highr_0.12.tar.gz
              bioconductor: true
              dependencies:
                - package_name: xfun
                  package_path: xfun_0.56.tar.gz
                  bioconductor: true
            - package_name: xfun
              package_path: xfun_0.56.tar.gz
              bioconductor: true
            - package_name: Rcpp
              package_path: Rcpp_1.1.1.tar.gz
              bioconductor: true
            - package_name: askpass
              package_path: askpass_1.2.1.tar.gz
              bioconductor: true
              dependencies:
                - package_name: sys
                  package_path: sys_3.4.3.tar.gz
                  bioconductor: true
            - package_name: utf8
              package_path: utf8_1.2.6.tar.gz
              bioconductor: true
            - package_name: bslib
              package_path: bslib_0.10.0.tar.gz
              bioconductor: true
              dependencies:
                - package_name: fastmap
                  package_path: fastmap_1.2.0.tar.gz
                  bioconductor: true
                - package_name: htmltools
                  package_path: htmltools_0.5.9.tar.gz
                  bioconductor: true
                  dependencies:
                    - package_name: digest
                      package_path: digest_0.6.39.tar.gz
                      bioconductor: true
                    - package_name: rlang
                      package_path: rlang_1.1.7.tar.gz
                      bioconductor: true
                    - package_name: base64enc
                      package_path: base64enc_0.1-6.tar.gz
                      bioconductor: true
                    - package_name: fastmap
                      package_path: fastmap_1.2.0.tar.gz
                      bioconductor: true
                - package_name: base64enc
                  package_path: base64enc_0.1-6.tar.gz
                  bioconductor: true
                - package_name: jsonlite
                  package_path: jsonlite_2.0.0.tar.gz
                  bioconductor: true
                - package_name: lifecycle
                  package_path: lifecycle_1.0.5.tar.gz
                  bioconductor: true
                  dependencies:
                    - package_name: cli
                      package_path: cli_3.6.5.tar.gz
                      bioconductor: true
                - package_name: cachem
                  package_path: cachem_1.1.0.tar.gz
                  bioconductor: true
                  dependencies:
                    - package_name: rlang
                      package_path: rlang_1.1.7.tar.gz
                      bioconductor: true
                    - package_name: fastmap
                      package_path: fastmap_1.2.0.tar.gz
                      bioconductor: true
                - package_name: mime
                  package_path: mime_0.13.tar.gz
                  bioconductor: true
                - package_name: rlang
                  package_path: rlang_1.1.7.tar.gz
                  bioconductor: true
                - package_name: digest
                  package_path: digest_0.6.39.tar.gz
                  bioconductor: true
                - package_name: cli
                  package_path: cli_3.6.5.tar.gz
                  bioconductor: true
                - package_name: memoise
                  package_path: memoise_2.0.1.tar.gz
                  bioconductor: true
                  dependencies:
                    - package_name: cachem
                      package_path: cachem_1.1.0.tar.gz
                      bioconductor: true
                      dependencies:
                        - package_name: rlang
                          package_path: rlang_1.1.7.tar.gz
                          bioconductor: true
                        - package_name: fastmap
                          package_path: fastmap_1.2.0.tar.gz
                          bioconductor: true
                    - package_name: rlang
                      package_path: rlang_1.1.7.tar.gz
                      bioconductor: true
                    - package_name: fastmap
                      package_path: fastmap_1.2.0.tar.gz
                      bioconductor: true
                - package_name: R6
                  package_path: R6_2.6.1.tar.gz
                  bioconductor: true
                - package_name: jquerylib
                  package_path: jquerylib_0.1.4.tar.gz
                  bioconductor: true
                  dependencies:
                    - package_name: base64enc
                      package_path: base64enc_0.1-6.tar.gz
                      bioconductor: true
                    - package_name: fastmap
                      package_path: fastmap_1.2.0.tar.gz
                      bioconductor: true
                    - package_name: rlang
                      package_path: rlang_1.1.7.tar.gz
                      bioconductor: true
                    - package_name: htmltools
                      package_path: htmltools_0.5.9.tar.gz
                      bioconductor: true
                      dependencies:
                        - package_name: digest
                          package_path: digest_0.6.39.tar.gz
                          bioconductor: true
                        - package_name: rlang
                          package_path: rlang_1.1.7.tar.gz
                          bioconductor: true
                        - package_name: base64enc
                          package_path: base64enc_0.1-6.tar.gz
                          bioconductor: true
                        - package_name: fastmap
                          package_path: fastmap_1.2.0.tar.gz
                          bioconductor: true
                    - package_name: digest
                      package_path: digest_0.6.39.tar.gz
                      bioconductor: true
                - package_name: sass
                  package_path: sass_0.4.10.tar.gz
                  bioconductor: true
                  dependencies:
                    - package_name: R6
                      package_path: R6_2.6.1.tar.gz
                      bioconductor: true
                    - package_name: htmltools
                      package_path: htmltools_0.5.9.tar.gz
                      bioconductor: true
                      dependencies:
                        - package_name: digest
                          package_path: digest_0.6.39.tar.gz
                          bioconductor: true
                        - package_name: rlang
                          package_path: rlang_1.1.7.tar.gz
                          bioconductor: true
                        - package_name: base64enc
                          package_path: base64enc_0.1-6.tar.gz
                          bioconductor: true
                        - package_name: fastmap
                          package_path: fastmap_1.2.0.tar.gz
                          bioconductor: true
                    - package_name: rlang
                      package_path: rlang_1.1.7.tar.gz
                      bioconductor: true
                    - package_name: base64enc
                      package_path: base64enc_0.1-6.tar.gz
                      bioconductor: true
                    - package_name: digest
                      package_path: digest_0.6.39.tar.gz
                      bioconductor: true
                    - package_name: fastmap
                      package_path: fastmap_1.2.0.tar.gz
                      bioconductor: true
                    - package_name: rappdirs
                      package_path: rappdirs_0.3.4.tar.gz
                      bioconductor: true
                  external_dependencies:
                    apt:
                        - name: make
                          dependencies:
                            - name: gcc-14-base
                            - name: libc6
                            - name: libgcc-s1
                - package_name: rappdirs
                  package_path: rappdirs_0.3.4.tar.gz
                  bioconductor: true
            - package_name: fontawesome
              package_path: fontawesome_0.5.3.tar.gz
              bioconductor: true
              dependencies:
                - package_name: digest
                  package_path: digest_0.6.39.tar.gz
                  bioconductor: true
                - package_name: fastmap
                  package_path: fastmap_1.2.0.tar.gz
                  bioconductor: true
                - package_name: htmltools
                  package_path: htmltools_0.5.9.tar.gz
                  bioconductor: true
                  dependencies:
                    - package_name: digest
                      package_path: digest_0.6.39.tar.gz
                      bioconductor: true
                    - package_name: rlang
                      package_path: rlang_1.1.7.tar.gz
                      bioconductor: true
                    - package_name: base64enc
                      package_path: base64enc_0.1-6.tar.gz
                      bioconductor: true
                    - package_name: fastmap
                      package_path: fastmap_1.2.0.tar.gz
                      bioconductor: true
                - package_name: rlang
                  package_path: rlang_1.1.7.tar.gz
                  bioconductor: true
                - package_name: base64enc
                  package_path: base64enc_0.1-6.tar.gz
                  bioconductor: true
            - package_name: jquerylib
              package_path: jquerylib_0.1.4.tar.gz
              bioconductor: true
              dependencies:
                - package_name: fastmap
                  package_path: fastmap_1.2.0.tar.gz
                  bioconductor: true
                - package_name: base64enc
                  package_path: base64enc_0.1-6.tar.gz
                  bioconductor: true
                - package_name: rlang
                  package_path: rlang_1.1.7.tar.gz
                  bioconductor: true
                - package_name: htmltools
                  package_path: htmltools_0.5.9.tar.gz
                  bioconductor: true
                  dependencies:
                    - package_name: digest
                      package_path: digest_0.6.39.tar.gz
                      bioconductor: true
                    - package_name: rlang
                      package_path: rlang_1.1.7.tar.gz
                      bioconductor: true
                    - package_name: base64enc
                      package_path: base64enc_0.1-6.tar.gz
                      bioconductor: true
                    - package_name: fastmap
                      package_path: fastmap_1.2.0.tar.gz
                      bioconductor: true
                - package_name: digest
                  package_path: digest_0.6.39.tar.gz
                  bioconductor: true
            - package_name: tinytex
              package_path: tinytex_0.58.tar.gz
              bioconductor: true
              dependencies:
                - package_name: xfun
                  package_path: xfun_0.56.tar.gz
                  bioconductor: true
            - package_name: stringi
              package_path: stringi_1.8.7.tar.gz
              bioconductor: true
              external_dependencies:
                apt:
                    - name: libicu-dev
                      dependencies:
                        - name: libc-dev
                          optional: true
                        - name: libc6
                        - name: libcrypt1
                        - name: gcc-14-base
                        - name: rpcsvc-proto
                        - name: libcrypt-dev
                        - name: linux-libc-dev
                        - name: libc-dev-bin
                        - name: libgcc-s1
                        - name: libstdc++6
                        - name: libc6-dev
                        - name: libicu74
                        - name: icu-devtools
            - package_name: sys
              package_path: sys_3.4.3.tar.gz
              bioconductor: true
            - package_name: cachem
              package_path: cachem_1.1.0.tar.gz
              bioconductor: true
              dependencies:
                - package_name: rlang
                  package_path: rlang_1.1.7.tar.gz
                  bioconductor: true
                - package_name: fastmap
                  package_path: fastmap_1.2.0.tar.gz
                  bioconductor: true
            - package_name: memoise
              package_path: memoise_2.0.1.tar.gz
              bioconductor: true
              dependencies:
                - package_name: cachem
                  package_path: cachem_1.1.0.tar.gz
                  bioconductor: true
                  dependencies:
                    - package_name: rlang
                      package_path: rlang_1.1.7.tar.gz
                      bioconductor: true
                    - package_name: fastmap
                      package_path: fastmap_1.2.0.tar.gz
                      bioconductor: true
                - package_name: fastmap
                  package_path: fastmap_1.2.0.tar.gz
                  bioconductor: true
                - package_name: rlang
                  package_path: rlang_1.1.7.tar.gz
                  bioconductor: true
            - package_name: sass
              package_path: sass_0.4.10.tar.gz
              bioconductor: true
              dependencies:
                - package_name: R6
                  package_path: R6_2.6.1.tar.gz
                  bioconductor: true
                - package_name: htmltools
                  package_path: htmltools_0.5.9.tar.gz
                  bioconductor: true
                  dependencies:
                    - package_name: digest
                      package_path: digest_0.6.39.tar.gz
                      bioconductor: true
                    - package_name: rlang
                      package_path: rlang_1.1.7.tar.gz
                      bioconductor: true
                    - package_name: base64enc
                      package_path: base64enc_0.1-6.tar.gz
                      bioconductor: true
                    - package_name: fastmap
                      package_path: fastmap_1.2.0.tar.gz
                      bioconductor: true
                - package_name: rlang
                  package_path: rlang_1.1.7.tar.gz
                  bioconductor: true
                - package_name: base64enc
                  package_path: base64enc_0.1-6.tar.gz
                  bioconductor: true
                - package_name: digest
                  package_path: digest_0.6.39.tar.gz
                  bioconductor: true
                - package_name: fastmap
                  package_path: fastmap_1.2.0.tar.gz
                  bioconductor: true
                - package_name: rappdirs
                  package_path: rappdirs_0.3.4.tar.gz
                  bioconductor: true
              external_dependencies:
                apt:
                    - name: make
                      dependencies:
                        - name: gcc-14-base
                        - name: libc6
                        - name: libgcc-s1
            - package_name: knitr
              package_path: knitr_1.51.tar.gz
              bioconductor: true
              dependencies:
                - package_name: highr
                  package_path: highr_0.12.tar.gz
                  bioconductor: true
                  dependencies:
                    - package_name: xfun
                      package_path: xfun_0.56.tar.gz
                      bioconductor: true
                - package_name: xfun
                  package_path: xfun_0.56.tar.gz
                  bioconductor: true
                - package_name: yaml
                  package_path: yaml_2.3.12.tar.gz
                  bioconductor: true
                - package_name: evaluate
                  package_path: evaluate_1.0.5.tar.gz
                  bioconductor: true
            - package_name: rappdirs
              package_path: rappdirs_0.3.4.tar.gz
              bioconductor: true
            - package_name: fs
              package_path: fs_1.6.7.tar.gz
              bioconductor: true
              external_dependencies:
                apt:
                    - name: make
                      dependencies:
                        - name: gcc-14-base
                        - name: libc6
                        - name: libgcc-s1
            - package_name: htmlwidgets
              package_path: htmlwidgets_1.6.4.tar.gz
              bioconductor: true
              dependencies:
                - package_name: jsonlite
                  package_path: jsonlite_2.0.0.tar.gz
                  bioconductor: true
                - package_name: htmltools
                  package_path: htmltools_0.5.9.tar.gz
                  bioconductor: true
                  dependencies:
                    - package_name: digest
                      package_path: digest_0.6.39.tar.gz
                      bioconductor: true
                    - package_name: rlang
                      package_path: rlang_1.1.7.tar.gz
                      bioconductor: true
                    - package_name: fastmap
                      package_path: fastmap_1.2.0.tar.gz
                      bioconductor: true
                    - package_name: base64enc
                      package_path: base64enc_0.1-6.tar.gz
                      bioconductor: true
                - package_name: base64enc
                  package_path: base64enc_0.1-6.tar.gz
                  bioconductor: true
                - package_name: digest
                  package_path: digest_0.6.39.tar.gz
                  bioconductor: true
                - package_name: fastmap
                  package_path: fastmap_1.2.0.tar.gz
                  bioconductor: true
                - package_name: rlang
                  package_path: rlang_1.1.7.tar.gz
                  bioconductor: true
                - package_name: yaml
                  package_path: yaml_2.3.12.tar.gz
                  bioconductor: true
                - package_name: highr
                  package_path: highr_0.12.tar.gz
                  bioconductor: true
                  dependencies:
                    - package_name: xfun
                      package_path: xfun_0.56.tar.gz
                      bioconductor: true
                - package_name: fontawesome
                  package_path: fontawesome_0.5.3.tar.gz
                  bioconductor: true
                  dependencies:
                    - package_name: digest
                      package_path: digest_0.6.39.tar.gz
                      bioconductor: true
                    - package_name: fastmap
                      package_path: fastmap_1.2.0.tar.gz
                      bioconductor: true
                    - package_name: htmltools
                      package_path: htmltools_0.5.9.tar.gz
                      bioconductor: true
                      dependencies:
                        - package_name: digest
                          package_path: digest_0.6.39.tar.gz
                          bioconductor: true
                        - package_name: rlang
                          package_path: rlang_1.1.7.tar.gz
                          bioconductor: true
                        - package_name: base64enc
                          package_path: base64enc_0.1-6.tar.gz
                          bioconductor: true
                        - package_name: fastmap
                          package_path: fastmap_1.2.0.tar.gz
                          bioconductor: true
                    - package_name: rlang
                      package_path: rlang_1.1.7.tar.gz
                      bioconductor: true
                    - package_name: base64enc
                      package_path: base64enc_0.1-6.tar.gz
                      bioconductor: true
                - package_name: jquerylib
                  package_path: jquerylib_0.1.4.tar.gz
                  bioconductor: true
                  dependencies:
                    - package_name: fastmap
                      package_path: fastmap_1.2.0.tar.gz
                      bioconductor: true
                    - package_name: base64enc
                      package_path: base64enc_0.1-6.tar.gz
                      bioconductor: true
                    - package_name: rlang
                      package_path: rlang_1.1.7.tar.gz
                      bioconductor: true
                    - package_name: htmltools
                      package_path: htmltools_0.5.9.tar.gz
                      bioconductor: true
                      dependencies:
                        - package_name: digest
                          package_path: digest_0.6.39.tar.gz
                          bioconductor: true
                        - package_name: rlang
                          package_path: rlang_1.1.7.tar.gz
                          bioconductor: true
                        - package_name: base64enc
                          package_path: base64enc_0.1-6.tar.gz
                          bioconductor: true
                        - package_name: fastmap
                          package_path: fastmap_1.2.0.tar.gz
                          bioconductor: true
                    - package_name: digest
                      package_path: digest_0.6.39.tar.gz
                      bioconductor: true
                - package_name: knitr
                  package_path: knitr_1.51.tar.gz
                  bioconductor: true
                  dependencies:
                    - package_name: yaml
                      package_path: yaml_2.3.12.tar.gz
                      bioconductor: true
                    - package_name: evaluate
                      package_path: evaluate_1.0.5.tar.gz
                      bioconductor: true
                    - package_name: xfun
                      package_path: xfun_0.56.tar.gz
                      bioconductor: true
                    - package_name: highr
                      package_path: highr_0.12.tar.gz
                      bioconductor: true
                      dependencies:
                        - package_name: xfun
                          package_path: xfun_0.56.tar.gz
                          bioconductor: true
                - package_name: cachem
                  package_path: cachem_1.1.0.tar.gz
                  bioconductor: true
                  dependencies:
                    - package_name: rlang
                      package_path: rlang_1.1.7.tar.gz
                      bioconductor: true
                    - package_name: fastmap
                      package_path: fastmap_1.2.0.tar.gz
                      bioconductor: true
                - package_name: lifecycle
                  package_path: lifecycle_1.0.5.tar.gz
                  bioconductor: true
                  dependencies:
                    - package_name: cli
                      package_path: cli_3.6.5.tar.gz
                      bioconductor: true
                - package_name: memoise
                  package_path: memoise_2.0.1.tar.gz
                  bioconductor: true
                  dependencies:
                    - package_name: cachem
                      package_path: cachem_1.1.0.tar.gz
                      bioconductor: true
                      dependencies:
                        - package_name: rlang
                          package_path: rlang_1.1.7.tar.gz
                          bioconductor: true
                        - package_name: fastmap
                          package_path: fastmap_1.2.0.tar.gz
                          bioconductor: true
                    - package_name: fastmap
                      package_path: fastmap_1.2.0.tar.gz
                      bioconductor: true
                    - package_name: rlang
                      package_path: rlang_1.1.7.tar.gz
                      bioconductor: true
                - package_name: mime
                  package_path: mime_0.13.tar.gz
                  bioconductor: true
                - package_name: sass
                  package_path: sass_0.4.10.tar.gz
                  bioconductor: true
                  dependencies:
                    - package_name: R6
                      package_path: R6_2.6.1.tar.gz
                      bioconductor: true
                    - package_name: htmltools
                      package_path: htmltools_0.5.9.tar.gz
                      bioconductor: true
                      dependencies:
                        - package_name: digest
                          package_path: digest_0.6.39.tar.gz
                          bioconductor: true
                        - package_name: rlang
                          package_path: rlang_1.1.7.tar.gz
                          bioconductor: true
                        - package_name: base64enc
                          package_path: base64enc_0.1-6.tar.gz
                          bioconductor: true
                        - package_name: fastmap
                          package_path: fastmap_1.2.0.tar.gz
                          bioconductor: true
                    - package_name: rlang
                      package_path: rlang_1.1.7.tar.gz
                      bioconductor: true
                    - package_name: base64enc
                      package_path: base64enc_0.1-6.tar.gz
                      bioconductor: true
                    - package_name: digest
                      package_path: digest_0.6.39.tar.gz
                      bioconductor: true
                    - package_name: fastmap
                      package_path: fastmap_1.2.0.tar.gz
                      bioconductor: true
                    - package_name: rappdirs
                      package_path: rappdirs_0.3.4.tar.gz
                      bioconductor: true
                  external_dependencies:
                    apt:
                        - name: make
                          dependencies:
                            - name: gcc-14-base
                            - name: libc6
                            - name: libgcc-s1
                - package_name: cli
                  package_path: cli_3.6.5.tar.gz
                  bioconductor: true
                - package_name: tinytex
                  package_path: tinytex_0.58.tar.gz
                  bioconductor: true
                  dependencies:
                    - package_name: xfun
                      package_path: xfun_0.56.tar.gz
                      bioconductor: true
                - package_name: R6
                  package_path: R6_2.6.1.tar.gz
                  bioconductor: true
                - package_name: rappdirs
                  package_path: rappdirs_0.3.4.tar.gz
                  bioconductor: true
                - package_name: rmarkdown
                  package_path: rmarkdown_2.30.tar.gz
                  bioconductor: true
                  dependencies:
                    - package_name: htmltools
                      package_path: htmltools_0.5.9.tar.gz
                      bioconductor: true
                      dependencies:
                        - package_name: digest
                          package_path: digest_0.6.39.tar.gz
                          bioconductor: true
                        - package_name: rlang
                          package_path: rlang_1.1.7.tar.gz
                          bioconductor: true
                        - package_name: base64enc
                          package_path: base64enc_0.1-6.tar.gz
                          bioconductor: true
                        - package_name: fastmap
                          package_path: fastmap_1.2.0.tar.gz
                          bioconductor: true
                    - package_name: fontawesome
                      package_path: fontawesome_0.5.3.tar.gz
                      bioconductor: true
                      dependencies:
                        - package_name: digest
                          package_path: digest_0.6.39.tar.gz
                          bioconductor: true
                        - package_name: fastmap
                          package_path: fastmap_1.2.0.tar.gz
                          bioconductor: true
                        - package_name: htmltools
                          package_path: htmltools_0.5.9.tar.gz
                          bioconductor: true
                          dependencies:
                            - package_name: digest
                              package_path: digest_0.6.39.tar.gz
                              bioconductor: true
                            - package_name: rlang
                              package_path: rlang_1.1.7.tar.gz
                              bioconductor: true
                            - package_name: base64enc
                              package_path: base64enc_0.1-6.tar.gz
                              bioconductor: true
                            - package_name: fastmap
                              package_path: fastmap_1.2.0.tar.gz
                              bioconductor: true
                        - package_name: rlang
                          package_path: rlang_1.1.7.tar.gz
                          bioconductor: true
                        - package_name: base64enc
                          package_path: base64enc_0.1-6.tar.gz
                          bioconductor: true
                    - package_name: jsonlite
                      package_path: jsonlite_2.0.0.tar.gz
                      bioconductor: true
                    - package_name: xfun
                      package_path: xfun_0.56.tar.gz
                      bioconductor: true
                    - package_name: yaml
                      package_path: yaml_2.3.12.tar.gz
                      bioconductor: true
                    - package_name: base64enc
                      package_path: base64enc_0.1-6.tar.gz
                      bioconductor: true
                    - package_name: cachem
                      package_path: cachem_1.1.0.tar.gz
                      bioconductor: true
                      dependencies:
                        - package_name: rlang
                          package_path: rlang_1.1.7.tar.gz
                          bioconductor: true
                        - package_name: fastmap
                          package_path: fastmap_1.2.0.tar.gz
                          bioconductor: true
                    - package_name: fastmap
                      package_path: fastmap_1.2.0.tar.gz
                      bioconductor: true
                    - package_name: lifecycle
                      package_path: lifecycle_1.0.5.tar.gz
                      bioconductor: true
                      dependencies:
                        - package_name: cli
                          package_path: cli_3.6.5.tar.gz
                          bioconductor: true
                    - package_name: memoise
                      package_path: memoise_2.0.1.tar.gz
                      bioconductor: true
                      dependencies:
                        - package_name: cachem
                          package_path: cachem_1.1.0.tar.gz
                          bioconductor: true
                          dependencies:
                            - package_name: rlang
                              package_path: rlang_1.1.7.tar.gz
                              bioconductor: true
                            - package_name: fastmap
                              package_path: fastmap_1.2.0.tar.gz
                              bioconductor: true
                        - package_name: fastmap
                          package_path: fastmap_1.2.0.tar.gz
                          bioconductor: true
                        - package_name: rlang
                          package_path: rlang_1.1.7.tar.gz
                          bioconductor: true
                    - package_name: mime
                      package_path: mime_0.13.tar.gz
                      bioconductor: true
                    - package_name: rlang
                      package_path: rlang_1.1.7.tar.gz
                      bioconductor: true
                    - package_name: jquerylib
                      package_path: jquerylib_0.1.4.tar.gz
                      bioconductor: true
                      dependencies:
                        - package_name: fastmap
                          package_path: fastmap_1.2.0.tar.gz
                          bioconductor: true
                        - package_name: htmltools
                          package_path: htmltools_0.5.9.tar.gz
                          bioconductor: true
                          dependencies:
                            - package_name: digest
                              package_path: digest_0.6.39.tar.gz
                              bioconductor: true
                            - package_name: rlang
                              package_path: rlang_1.1.7.tar.gz
                              bioconductor: true
                            - package_name: base64enc
                              package_path: base64enc_0.1-6.tar.gz
                              bioconductor: true
                            - package_name: fastmap
                              package_path: fastmap_1.2.0.tar.gz
                              bioconductor: true
                        - package_name: rlang
                          package_path: rlang_1.1.7.tar.gz
                          bioconductor: true
                        - package_name: base64enc
                          package_path: base64enc_0.1-6.tar.gz
                          bioconductor: true
                        - package_name: digest
                          package_path: digest_0.6.39.tar.gz
                          bioconductor: true
                    - package_name: digest
                      package_path: digest_0.6.39.tar.gz
                      bioconductor: true
                    - package_name: highr
                      package_path: highr_0.12.tar.gz
                      bioconductor: true
                      dependencies:
                        - package_name: xfun
                          package_path: xfun_0.56.tar.gz
                          bioconductor: true
                    - package_name: cli
                      package_path: cli_3.6.5.tar.gz
                      bioconductor: true
                    - package_name: R6
                      package_path: R6_2.6.1.tar.gz
                      bioconductor: true
                    - package_name: rappdirs
                      package_path: rappdirs_0.3.4.tar.gz
                      bioconductor: true
                    - package_name: tinytex
                      package_path: tinytex_0.58.tar.gz
                      bioconductor: true
                      dependencies:
                        - package_name: xfun
                          package_path: xfun_0.56.tar.gz
                          bioconductor: true
                    - package_name: fs
                      package_path: fs_1.6.7.tar.gz
                      bioconductor: true
                      external_dependencies:
                        apt:
                            - name: make
                              dependencies:
                                - name: gcc-14-base
                                - name: libc6
                                - name: libgcc-s1
                    - package_name: sass
                      package_path: sass_0.4.10.tar.gz
                      bioconductor: true
                      dependencies:
                        - package_name: R6
                          package_path: R6_2.6.1.tar.gz
                          bioconductor: true
                        - package_name: rappdirs
                          package_path: rappdirs_0.3.4.tar.gz
                          bioconductor: true
                        - package_name: rlang
                          package_path: rlang_1.1.7.tar.gz
                          bioconductor: true
                        - package_name: base64enc
                          package_path: base64enc_0.1-6.tar.gz
                          bioconductor: true
                        - package_name: digest
                          package_path: digest_0.6.39.tar.gz
                          bioconductor: true
                        - package_name: fastmap
                          package_path: fastmap_1.2.0.tar.gz
                          bioconductor: true
                        - package_name: htmltools
                          package_path: htmltools_0.5.9.tar.gz
                          bioconductor: true
                          dependencies:
                            - package_name: digest
                              package_path: digest_0.6.39.tar.gz
                              bioconductor: true
                            - package_name: rlang
                              package_path: rlang_1.1.7.tar.gz
                              bioconductor: true
                            - package_name: base64enc
                              package_path: base64enc_0.1-6.tar.gz
                              bioconductor: true
                            - package_name: fastmap
                              package_path: fastmap_1.2.0.tar.gz
                              bioconductor: true
                      external_dependencies:
                        apt:
                            - name: make
                              dependencies:
                                - name: gcc-14-base
                                - name: libc6
                                - name: libgcc-s1
                  external_dependencies:
                    apt:
                        - name: pandoc
                          dependencies:
                            - name: gcc-14-base
                            - name: libgcc-s1
                            - name: libc6
                            - name: libstdc++6
                            - name: zlib1g
                            - name: libyaml-0-2
                            - name: liblua5.4-0
                            - name: libgmp10
                            - name: libffi8
                            - name: pandoc-data
                - package_name: bslib
                  package_path: bslib_0.10.0.tar.gz
                  bioconductor: true
                  dependencies:
                    - package_name: cachem
                      package_path: cachem_1.1.0.tar.gz
                      bioconductor: true
                      dependencies:
                        - package_name: rlang
                          package_path: rlang_1.1.7.tar.gz
                          bioconductor: true
                        - package_name: fastmap
                          package_path: fastmap_1.2.0.tar.gz
                          bioconductor: true
                    - package_name: htmltools
                      package_path: htmltools_0.5.9.tar.gz
                      bioconductor: true
                      dependencies:
                        - package_name: digest
                          package_path: digest_0.6.39.tar.gz
                          bioconductor: true
                        - package_name: rlang
                          package_path: rlang_1.1.7.tar.gz
                          bioconductor: true
                        - package_name: base64enc
                          package_path: base64enc_0.1-6.tar.gz
                          bioconductor: true
                        - package_name: fastmap
                          package_path: fastmap_1.2.0.tar.gz
                          bioconductor: true
                    - package_name: jquerylib
                      package_path: jquerylib_0.1.4.tar.gz
                      bioconductor: true
                      dependencies:
                        - package_name: fastmap
                          package_path: fastmap_1.2.0.tar.gz
                          bioconductor: true
                        - package_name: base64enc
                          package_path: base64enc_0.1-6.tar.gz
                          bioconductor: true
                        - package_name: rlang
                          package_path: rlang_1.1.7.tar.gz
                          bioconductor: true
                        - package_name: htmltools
                          package_path: htmltools_0.5.9.tar.gz
                          bioconductor: true
                          dependencies:
                            - package_name: digest
                              package_path: digest_0.6.39.tar.gz
                              bioconductor: true
                            - package_name: rlang
                              package_path: rlang_1.1.7.tar.gz
                              bioconductor: true
                            - package_name: base64enc
                              package_path: base64enc_0.1-6.tar.gz
                              bioconductor: true
                            - package_name: fastmap
                              package_path: fastmap_1.2.0.tar.gz
                              bioconductor: true
                        - package_name: digest
                          package_path: digest_0.6.39.tar.gz
                          bioconductor: true
                    - package_name: jsonlite
                      package_path: jsonlite_2.0.0.tar.gz
                      bioconductor: true
                    - package_name: lifecycle
                      package_path: lifecycle_1.0.5.tar.gz
                      bioconductor: true
                      dependencies:
                        - package_name: cli
                          package_path: cli_3.6.5.tar.gz
                          bioconductor: true
                    - package_name: memoise
                      package_path: memoise_2.0.1.tar.gz
                      bioconductor: true
                      dependencies:
                        - package_name: cachem
                          package_path: cachem_1.1.0.tar.gz
                          bioconductor: true
                          dependencies:
                            - package_name: rlang
                              package_path: rlang_1.1.7.tar.gz
                              bioconductor: true
                            - package_name: fastmap
                              package_path: fastmap_1.2.0.tar.gz
                              bioconductor: true
                        - package_name: fastmap
                          package_path: fastmap_1.2.0.tar.gz
                          bioconductor: true
                        - package_name: rlang
                          package_path: rlang_1.1.7.tar.gz
                          bioconductor: true
                    - package_name: mime
                      package_path: mime_0.13.tar.gz
                      bioconductor: true
                    - package_name: rlang
                      package_path: rlang_1.1.7.tar.gz
                      bioconductor: true
                    - package_name: sass
                      package_path: sass_0.4.10.tar.gz
                      bioconductor: true
                      dependencies:
                        - package_name: R6
                          package_path: R6_2.6.1.tar.gz
                          bioconductor: true
                        - package_name: htmltools
                          package_path: htmltools_0.5.9.tar.gz
                          bioconductor: true
                          dependencies:
                            - package_name: digest
                              package_path: digest_0.6.39.tar.gz
                              bioconductor: true
                            - package_name: rlang
                              package_path: rlang_1.1.7.tar.gz
                              bioconductor: true
                            - package_name: base64enc
                              package_path: base64enc_0.1-6.tar.gz
                              bioconductor: true
                            - package_name: fastmap
                              package_path: fastmap_1.2.0.tar.gz
                              bioconductor: true
                        - package_name: rlang
                          package_path: rlang_1.1.7.tar.gz
                          bioconductor: true
                        - package_name: base64enc
                          package_path: base64enc_0.1-6.tar.gz
                          bioconductor: true
                        - package_name: digest
                          package_path: digest_0.6.39.tar.gz
                          bioconductor: true
                        - package_name: fastmap
                          package_path: fastmap_1.2.0.tar.gz
                          bioconductor: true
                        - package_name: rappdirs
                          package_path: rappdirs_0.3.4.tar.gz
                          bioconductor: true
                      external_dependencies:
                        apt:
                            - name: make
                              dependencies:
                                - name: gcc-14-base
                                - name: libc6
                                - name: libgcc-s1
                    - package_name: digest
                      package_path: digest_0.6.39.tar.gz
                      bioconductor: true
                    - package_name: base64enc
                      package_path: base64enc_0.1-6.tar.gz
                      bioconductor: true
                    - package_name: fastmap
                      package_path: fastmap_1.2.0.tar.gz
                      bioconductor: true
                    - package_name: cli
                      package_path: cli_3.6.5.tar.gz
                      bioconductor: true
                    - package_name: R6
                      package_path: R6_2.6.1.tar.gz
                      bioconductor: true
                    - package_name: rappdirs
                      package_path: rappdirs_0.3.4.tar.gz
                      bioconductor: true
                    - package_name: fs
                      package_path: fs_1.6.7.tar.gz
                      bioconductor: true
                      external_dependencies:
                        apt:
                            - name: make
                              dependencies:
                                - name: gcc-14-base
                                - name: libc6
                                - name: libgcc-s1
            - package_name: rmarkdown
              package_path: rmarkdown_2.30.tar.gz
              bioconductor: true
              dependencies:
                - package_name: htmltools
                  package_path: htmltools_0.5.9.tar.gz
                  bioconductor: true
                  dependencies:
                    - package_name: digest
                      package_path: digest_0.6.39.tar.gz
                      bioconductor: true
                    - package_name: rlang
                      package_path: rlang_1.1.7.tar.gz
                      bioconductor: true
                    - package_name: base64enc
                      package_path: base64enc_0.1-6.tar.gz
                      bioconductor: true
                    - package_name: fastmap
                      package_path: fastmap_1.2.0.tar.gz
                      bioconductor: true
                - package_name: jquerylib
                  package_path: jquerylib_0.1.4.tar.gz
                  bioconductor: true
                  dependencies:
                    - package_name: fastmap
                      package_path: fastmap_1.2.0.tar.gz
                      bioconductor: true
                    - package_name: base64enc
                      package_path: base64enc_0.1-6.tar.gz
                      bioconductor: true
                    - package_name: rlang
                      package_path: rlang_1.1.7.tar.gz
                      bioconductor: true
                    - package_name: htmltools
                      package_path: htmltools_0.5.9.tar.gz
                      bioconductor: true
                      dependencies:
                        - package_name: digest
                          package_path: digest_0.6.39.tar.gz
                          bioconductor: true
                        - package_name: rlang
                          package_path: rlang_1.1.7.tar.gz
                          bioconductor: true
                        - package_name: base64enc
                          package_path: base64enc_0.1-6.tar.gz
                          bioconductor: true
                        - package_name: fastmap
                          package_path: fastmap_1.2.0.tar.gz
                          bioconductor: true
                    - package_name: digest
                      package_path: digest_0.6.39.tar.gz
                      bioconductor: true
                - package_name: jsonlite
                  package_path: jsonlite_2.0.0.tar.gz
                  bioconductor: true
                - package_name: knitr
                  package_path: knitr_1.51.tar.gz
                  bioconductor: true
                  dependencies:
                    - package_name: yaml
                      package_path: yaml_2.3.12.tar.gz
                      bioconductor: true
                    - package_name: evaluate
                      package_path: evaluate_1.0.5.tar.gz
                      bioconductor: true
                    - package_name: xfun
                      package_path: xfun_0.56.tar.gz
                      bioconductor: true
                    - package_name: highr
                      package_path: highr_0.12.tar.gz
                      bioconductor: true
                      dependencies:
                        - package_name: xfun
                          package_path: xfun_0.56.tar.gz
                          bioconductor: true
                - package_name: fontawesome
                  package_path: fontawesome_0.5.3.tar.gz
                  bioconductor: true
                  dependencies:
                    - package_name: digest
                      package_path: digest_0.6.39.tar.gz
                      bioconductor: true
                    - package_name: fastmap
                      package_path: fastmap_1.2.0.tar.gz
                      bioconductor: true
                    - package_name: htmltools
                      package_path: htmltools_0.5.9.tar.gz
                      bioconductor: true
                      dependencies:
                        - package_name: digest
                          package_path: digest_0.6.39.tar.gz
                          bioconductor: true
                        - package_name: rlang
                          package_path: rlang_1.1.7.tar.gz
                          bioconductor: true
                        - package_name: base64enc
                          package_path: base64enc_0.1-6.tar.gz
                          bioconductor: true
                        - package_name: fastmap
                          package_path: fastmap_1.2.0.tar.gz
                          bioconductor: true
                    - package_name: rlang
                      package_path: rlang_1.1.7.tar.gz
                      bioconductor: true
                    - package_name: base64enc
                      package_path: base64enc_0.1-6.tar.gz
                      bioconductor: true
                - package_name: tinytex
                  package_path: tinytex_0.58.tar.gz
                  bioconductor: true
                  dependencies:
                    - package_name: xfun
                      package_path: xfun_0.56.tar.gz
                      bioconductor: true
                - package_name: bslib
                  package_path: bslib_0.10.0.tar.gz
                  bioconductor: true
                  dependencies:
                    - package_name: fastmap
                      package_path: fastmap_1.2.0.tar.gz
                      bioconductor: true
                    - package_name: htmltools
                      package_path: htmltools_0.5.9.tar.gz
                      bioconductor: true
                      dependencies:
                        - package_name: digest
                          package_path: digest_0.6.39.tar.gz
                          bioconductor: true
                        - package_name: rlang
                          package_path: rlang_1.1.7.tar.gz
                          bioconductor: true
                        - package_name: base64enc
                          package_path: base64enc_0.1-6.tar.gz
                          bioconductor: true
                        - package_name: fastmap
                          package_path: fastmap_1.2.0.tar.gz
                          bioconductor: true
                    - package_name: base64enc
                      package_path: base64enc_0.1-6.tar.gz
                      bioconductor: true
                    - package_name: jsonlite
                      package_path: jsonlite_2.0.0.tar.gz
                      bioconductor: true
                    - package_name: lifecycle
                      package_path: lifecycle_1.0.5.tar.gz
                      bioconductor: true
                      dependencies:
                        - package_name: cli
                          package_path: cli_3.6.5.tar.gz
                          bioconductor: true
                    - package_name: cachem
                      package_path: cachem_1.1.0.tar.gz
                      bioconductor: true
                      dependencies:
                        - package_name: rlang
                          package_path: rlang_1.1.7.tar.gz
                          bioconductor: true
                        - package_name: fastmap
                          package_path: fastmap_1.2.0.tar.gz
                          bioconductor: true
                    - package_name: mime
                      package_path: mime_0.13.tar.gz
                      bioconductor: true
                    - package_name: rlang
                      package_path: rlang_1.1.7.tar.gz
                      bioconductor: true
                    - package_name: digest
                      package_path: digest_0.6.39.tar.gz
                      bioconductor: true
                    - package_name: cli
                      package_path: cli_3.6.5.tar.gz
                      bioconductor: true
                    - package_name: memoise
                      package_path: memoise_2.0.1.tar.gz
                      bioconductor: true
                      dependencies:
                        - package_name: cachem
                          package_path: cachem_1.1.0.tar.gz
                          bioconductor: true
                          dependencies:
                            - package_name: rlang
                              package_path: rlang_1.1.7.tar.gz
                              bioconductor: true
                            - package_name: fastmap
                              package_path: fastmap_1.2.0.tar.gz
                              bioconductor: true
                        - package_name: rlang
                          package_path: rlang_1.1.7.tar.gz
                          bioconductor: true
                        - package_name: fastmap
                          package_path: fastmap_1.2.0.tar.gz
                          bioconductor: true
                    - package_name: R6
                      package_path: R6_2.6.1.tar.gz
                      bioconductor: true
                    - package_name: jquerylib
                      package_path: jquerylib_0.1.4.tar.gz
                      bioconductor: true
                      dependencies:
                        - package_name: base64enc
                          package_path: base64enc_0.1-6.tar.gz
                          bioconductor: true
                        - package_name: fastmap
                          package_path: fastmap_1.2.0.tar.gz
                          bioconductor: true
                        - package_name: rlang
                          package_path: rlang_1.1.7.tar.gz
                          bioconductor: true
                        - package_name: htmltools
                          package_path: htmltools_0.5.9.tar.gz
                          bioconductor: true
                          dependencies:
                            - package_name: digest
                              package_path: digest_0.6.39.tar.gz
                              bioconductor: true
                            - package_name: rlang
                              package_path: rlang_1.1.7.tar.gz
                              bioconductor: true
                            - package_name: base64enc
                              package_path: base64enc_0.1-6.tar.gz
                              bioconductor: true
                            - package_name: fastmap
                              package_path: fastmap_1.2.0.tar.gz
                              bioconductor: true
                        - package_name: digest
                          package_path: digest_0.6.39.tar.gz
                          bioconductor: true
                    - package_name: sass
                      package_path: sass_0.4.10.tar.gz
                      bioconductor: true
                      dependencies:
                        - package_name: R6
                          package_path: R6_2.6.1.tar.gz
                          bioconductor: true
                        - package_name: htmltools
                          package_path: htmltools_0.5.9.tar.gz
                          bioconductor: true
                          dependencies:
                            - package_name: digest
                              package_path: digest_0.6.39.tar.gz
                              bioconductor: true
                            - package_name: rlang
                              package_path: rlang_1.1.7.tar.gz
                              bioconductor: true
                            - package_name: base64enc
                              package_path: base64enc_0.1-6.tar.gz
                              bioconductor: true
                            - package_name: fastmap
                              package_path: fastmap_1.2.0.tar.gz
                              bioconductor: true
                        - package_name: rlang
                          package_path: rlang_1.1.7.tar.gz
                          bioconductor: true
                        - package_name: base64enc
                          package_path: base64enc_0.1-6.tar.gz
                          bioconductor: true
                        - package_name: digest
                          package_path: digest_0.6.39.tar.gz
                          bioconductor: true
                        - package_name: fastmap
                          package_path: fastmap_1.2.0.tar.gz
                          bioconductor: true
                        - package_name: rappdirs
                          package_path: rappdirs_0.3.4.tar.gz
                          bioconductor: true
                      external_dependencies:
                        apt:
                            - name: make
                              dependencies:
                                - name: gcc-14-base
                                - name: libc6
                                - name: libgcc-s1
                    - package_name: rappdirs
                      package_path: rappdirs_0.3.4.tar.gz
                      bioconductor: true
                - package_name: evaluate
                  package_path: evaluate_1.0.5.tar.gz
                  bioconductor: true
                - package_name: xfun
                  package_path: xfun_0.56.tar.gz
                  bioconductor: true
                - package_name: yaml
                  package_path: yaml_2.3.12.tar.gz
                  bioconductor: true
                - package_name: base64enc
                  package_path: base64enc_0.1-6.tar.gz
                  bioconductor: true
                - package_name: cachem
                  package_path: cachem_1.1.0.tar.gz
                  bioconductor: true
                  dependencies:
                    - package_name: rlang
                      package_path: rlang_1.1.7.tar.gz
                      bioconductor: true
                    - package_name: fastmap
                      package_path: fastmap_1.2.0.tar.gz
                      bioconductor: true
                - package_name: fastmap
                  package_path: fastmap_1.2.0.tar.gz
                  bioconductor: true
                - package_name: lifecycle
                  package_path: lifecycle_1.0.5.tar.gz
                  bioconductor: true
                  dependencies:
                    - package_name: cli
                      package_path: cli_3.6.5.tar.gz
                      bioconductor: true
                - package_name: memoise
                  package_path: memoise_2.0.1.tar.gz
                  bioconductor: true
                  dependencies:
                    - package_name: cachem
                      package_path: cachem_1.1.0.tar.gz
                      bioconductor: true
                      dependencies:
                        - package_name: rlang
                          package_path: rlang_1.1.7.tar.gz
                          bioconductor: true
                        - package_name: fastmap
                          package_path: fastmap_1.2.0.tar.gz
                          bioconductor: true
                    - package_name: fastmap
                      package_path: fastmap_1.2.0.tar.gz
                      bioconductor: true
                    - package_name: rlang
                      package_path: rlang_1.1.7.tar.gz
                      bioconductor: true
                - package_name: mime
                  package_path: mime_0.13.tar.gz
                  bioconductor: true
                - package_name: rlang
                  package_path: rlang_1.1.7.tar.gz
                  bioconductor: true
                - package_name: sass
                  package_path: sass_0.4.10.tar.gz
                  bioconductor: true
                  dependencies:
                    - package_name: R6
                      package_path: R6_2.6.1.tar.gz
                      bioconductor: true
                    - package_name: htmltools
                      package_path: htmltools_0.5.9.tar.gz
                      bioconductor: true
                      dependencies:
                        - package_name: digest
                          package_path: digest_0.6.39.tar.gz
                          bioconductor: true
                        - package_name: rlang
                          package_path: rlang_1.1.7.tar.gz
                          bioconductor: true
                        - package_name: base64enc
                          package_path: base64enc_0.1-6.tar.gz
                          bioconductor: true
                        - package_name: fastmap
                          package_path: fastmap_1.2.0.tar.gz
                          bioconductor: true
                    - package_name: rlang
                      package_path: rlang_1.1.7.tar.gz
                      bioconductor: true
                    - package_name: base64enc
                      package_path: base64enc_0.1-6.tar.gz
                      bioconductor: true
                    - package_name: digest
                      package_path: digest_0.6.39.tar.gz
                      bioconductor: true
                    - package_name: fastmap
                      package_path: fastmap_1.2.0.tar.gz
                      bioconductor: true
                    - package_name: rappdirs
                      package_path: rappdirs_0.3.4.tar.gz
                      bioconductor: true
                  external_dependencies:
                    apt:
                        - name: make
                          dependencies:
                            - name: gcc-14-base
                            - name: libc6
                            - name: libgcc-s1
                - package_name: digest
                  package_path: digest_0.6.39.tar.gz
                  bioconductor: true
                - package_name: highr
                  package_path: highr_0.12.tar.gz
                  bioconductor: true
                  dependencies:
                    - package_name: xfun
                      package_path: xfun_0.56.tar.gz
                      bioconductor: true
                - package_name: cli
                  package_path: cli_3.6.5.tar.gz
                  bioconductor: true
                - package_name: R6
                  package_path: R6_2.6.1.tar.gz
                  bioconductor: true
                - package_name: rappdirs
                  package_path: rappdirs_0.3.4.tar.gz
                  bioconductor: true
                - package_name: fs
                  package_path: fs_1.6.7.tar.gz
                  bioconductor: true
                  external_dependencies:
                    apt:
                        - name: make
                          dependencies:
                            - name: gcc-14-base
                            - name: libc6
                            - name: libgcc-s1
              external_dependencies:
                apt:
                    - name: pandoc
                      dependencies:
                        - name: gcc-14-base
                        - name: libgcc-s1
                        - name: libc6
                        - name: libstdc++6
                        - name: zlib1g
                        - name: libyaml-0-2
                        - name: liblua5.4-0
                        - name: libgmp10
                        - name: libffi8
                        - name: pandoc-data
        - package_name: shiny
          package_path: shiny_1.13.0.tar.gz
          bioconductor: true
          dependencies:
            - package_name: cli
              package_path: cli_3.6.5.tar.gz
              bioconductor: true
            - package_name: fastmap
              package_path: fastmap_1.2.0.tar.gz
              bioconductor: true
            - package_name: commonmark
              package_path: commonmark_2.0.0.tar.gz
              bioconductor: true
            - package_name: glue
              package_path: glue_1.8.0.tar.gz
              bioconductor: true
            - package_name: cachem
              package_path: cachem_1.1.0.tar.gz
              bioconductor: true
              dependencies:
                - package_name: rlang
                  package_path: rlang_1.1.7.tar.gz
                  bioconductor: true
                - package_name: fastmap
                  package_path: fastmap_1.2.0.tar.gz
                  bioconductor: true
            - package_name: htmltools
              package_path: htmltools_0.5.9.tar.gz
              bioconductor: true
              dependencies:
                - package_name: digest
                  package_path: digest_0.6.39.tar.gz
                  bioconductor: true
                - package_name: rlang
                  package_path: rlang_1.1.7.tar.gz
                  bioconductor: true
                - package_name: base64enc
                  package_path: base64enc_0.1-6.tar.gz
                  bioconductor: true
                - package_name: fastmap
                  package_path: fastmap_1.2.0.tar.gz
                  bioconductor: true
            - package_name: jsonlite
              package_path: jsonlite_2.0.0.tar.gz
              bioconductor: true
            - package_name: fontawesome
              package_path: fontawesome_0.5.3.tar.gz
              bioconductor: true
              dependencies:
                - package_name: digest
                  package_path: digest_0.6.39.tar.gz
                  bioconductor: true
                - package_name: fastmap
                  package_path: fastmap_1.2.0.tar.gz
                  bioconductor: true
                - package_name: htmltools
                  package_path: htmltools_0.5.9.tar.gz
                  bioconductor: true
                  dependencies:
                    - package_name: digest
                      package_path: digest_0.6.39.tar.gz
                      bioconductor: true
                    - package_name: rlang
                      package_path: rlang_1.1.7.tar.gz
                      bioconductor: true
                    - package_name: base64enc
                      package_path: base64enc_0.1-6.tar.gz
                      bioconductor: true
                    - package_name: fastmap
                      package_path: fastmap_1.2.0.tar.gz
                      bioconductor: true
                - package_name: rlang
                  package_path: rlang_1.1.7.tar.gz
                  bioconductor: true
                - package_name: base64enc
                  package_path: base64enc_0.1-6.tar.gz
                  bioconductor: true
            - package_name: lifecycle
              package_path: lifecycle_1.0.5.tar.gz
              bioconductor: true
              dependencies:
                - package_name: cli
                  package_path: cli_3.6.5.tar.gz
                  bioconductor: true
            - package_name: mime
              package_path: mime_0.13.tar.gz
              bioconductor: true
            - package_name: later
              package_path: later_1.4.8.tar.gz
              bioconductor: true
              dependencies:
                - package_name: rlang
                  package_path: rlang_1.1.7.tar.gz
                  bioconductor: true
                - package_name: Rcpp
                  package_path: Rcpp_1.1.1.tar.gz
                  bioconductor: true
            - package_name: R6
              package_path: R6_2.6.1.tar.gz
              bioconductor: true
            - package_name: rlang
              package_path: rlang_1.1.7.tar.gz
              bioconductor: true
            - package_name: withr
              package_path: withr_3.0.2.tar.gz
              bioconductor: true
            - package_name: base64enc
              package_path: base64enc_0.1-6.tar.gz
              bioconductor: true
            - package_name: jquerylib
              package_path: jquerylib_0.1.4.tar.gz
              bioconductor: true
              dependencies:
                - package_name: fastmap
                  package_path: fastmap_1.2.0.tar.gz
                  bioconductor: true
                - package_name: base64enc
                  package_path: base64enc_0.1-6.tar.gz
                  bioconductor: true
                - package_name: rlang
                  package_path: rlang_1.1.7.tar.gz
                  bioconductor: true
                - package_name: htmltools
                  package_path: htmltools_0.5.9.tar.gz
                  bioconductor: true
                  dependencies:
                    - package_name: digest
                      package_path: digest_0.6.39.tar.gz
                      bioconductor: true
                    - package_name: rlang
                      package_path: rlang_1.1.7.tar.gz
                      bioconductor: true
                    - package_name: base64enc
                      package_path: base64enc_0.1-6.tar.gz
                      bioconductor: true
                    - package_name: fastmap
                      package_path: fastmap_1.2.0.tar.gz
                      bioconductor: true
                - package_name: digest
                  package_path: digest_0.6.39.tar.gz
                  bioconductor: true
            - package_name: memoise
              package_path: memoise_2.0.1.tar.gz
              bioconductor: true
              dependencies:
                - package_name: cachem
                  package_path: cachem_1.1.0.tar.gz
                  bioconductor: true
                  dependencies:
                    - package_name: rlang
                      package_path: rlang_1.1.7.tar.gz
                      bioconductor: true
                    - package_name: fastmap
                      package_path: fastmap_1.2.0.tar.gz
                      bioconductor: true
                - package_name: fastmap
                  package_path: fastmap_1.2.0.tar.gz
                  bioconductor: true
                - package_name: rlang
                  package_path: rlang_1.1.7.tar.gz
                  bioconductor: true
            - package_name: sass
              package_path: sass_0.4.10.tar.gz
              bioconductor: true
              dependencies:
                - package_name: R6
                  package_path: R6_2.6.1.tar.gz
                  bioconductor: true
                - package_name: htmltools
                  package_path: htmltools_0.5.9.tar.gz
                  bioconductor: true
                  dependencies:
                    - package_name: digest
                      package_path: digest_0.6.39.tar.gz
                      bioconductor: true
                    - package_name: rlang
                      package_path: rlang_1.1.7.tar.gz
                      bioconductor: true
                    - package_name: base64enc
                      package_path: base64enc_0.1-6.tar.gz
                      bioconductor: true
                    - package_name: fastmap
                      package_path: fastmap_1.2.0.tar.gz
                      bioconductor: true
                - package_name: rlang
                  package_path: rlang_1.1.7.tar.gz
                  bioconductor: true
                - package_name: base64enc
                  package_path: base64enc_0.1-6.tar.gz
                  bioconductor: true
                - package_name: digest
                  package_path: digest_0.6.39.tar.gz
                  bioconductor: true
                - package_name: fastmap
                  package_path: fastmap_1.2.0.tar.gz
                  bioconductor: true
                - package_name: rappdirs
                  package_path: rappdirs_0.3.4.tar.gz
                  bioconductor: true
              external_dependencies:
                apt:
                    - name: make
                      dependencies:
                        - name: gcc-14-base
                        - name: libc6
                        - name: libgcc-s1
            - package_name: digest
              package_path: digest_0.6.39.tar.gz
              bioconductor: true
            - package_name: Rcpp
              package_path: Rcpp_1.1.1.tar.gz
              bioconductor: true
            - package_name: magrittr
              package_path: magrittr_2.0.4.tar.gz
              bioconductor: true
            - package_name: sourcetools
              package_path: sourcetools_0.1.7-1.tar.gz
              bioconductor: true
            - package_name: rappdirs
              package_path: rappdirs_0.3.4.tar.gz
              bioconductor: true
            - package_name: httpuv
              package_path: httpuv_1.6.16.tar.gz
              bioconductor: true
              dependencies:
                - package_name: Rcpp
                  package_path: Rcpp_1.1.1.tar.gz
                  bioconductor: true
                - package_name: R6
                  package_path: R6_2.6.1.tar.gz
                  bioconductor: true
                - package_name: rlang
                  package_path: rlang_1.1.7.tar.gz
                  bioconductor: true
                - package_name: fastmap
                  package_path: fastmap_1.2.0.tar.gz
                  bioconductor: true
                - package_name: lifecycle
                  package_path: lifecycle_1.0.5.tar.gz
                  bioconductor: true
                  dependencies:
                    - package_name: cli
                      package_path: cli_3.6.5.tar.gz
                      bioconductor: true
                - package_name: magrittr
                  package_path: magrittr_2.0.4.tar.gz
                  bioconductor: true
                - package_name: cli
                  package_path: cli_3.6.5.tar.gz
                  bioconductor: true
                - package_name: otel
                  package_path: otel_0.2.0.tar.gz
                  bioconductor: true
                - package_name: promises
                  package_path: promises_1.5.0.tar.gz
                  bioconductor: true
                  dependencies:
                    - package_name: magrittr
                      package_path: magrittr_2.0.4.tar.gz
                      bioconductor: true
                    - package_name: later
                      package_path: later_1.4.8.tar.gz
                      bioconductor: true
                      dependencies:
                        - package_name: rlang
                          package_path: rlang_1.1.7.tar.gz
                          bioconductor: true
                        - package_name: Rcpp
                          package_path: Rcpp_1.1.1.tar.gz
                          bioconductor: true
                    - package_name: R6
                      package_path: R6_2.6.1.tar.gz
                      bioconductor: true
                    - package_name: rlang
                      package_path: rlang_1.1.7.tar.gz
                      bioconductor: true
                    - package_name: Rcpp
                      package_path: Rcpp_1.1.1.tar.gz
                      bioconductor: true
                    - package_name: fastmap
                      package_path: fastmap_1.2.0.tar.gz
                      bioconductor: true
                    - package_name: lifecycle
                      package_path: lifecycle_1.0.5.tar.gz
                      bioconductor: true
                      dependencies:
                        - package_name: cli
                          package_path: cli_3.6.5.tar.gz
                          bioconductor: true
                    - package_name: cli
                      package_path: cli_3.6.5.tar.gz
                      bioconductor: true
                    - package_name: otel
                      package_path: otel_0.2.0.tar.gz
                      bioconductor: true
                - package_name: later
                  package_path: later_1.4.8.tar.gz
                  bioconductor: true
                  dependencies:
                    - package_name: Rcpp
                      package_path: Rcpp_1.1.1.tar.gz
                      bioconductor: true
                    - package_name: rlang
                      package_path: rlang_1.1.7.tar.gz
                      bioconductor: true
              external_dependencies:
                apt:
                    - name: make
                      dependencies:
                        - name: gcc-14-base
                        - name: libc6
                        - name: libgcc-s1
                    - name: zlib1g-dev
                      dependencies:
                        - name: libc-dev
                          optional: true
                        - name: gcc-14-base
                        - name: libc6
                        - name: libcrypt1
                        - name: libgcc-s1
                        - name: rpcsvc-proto
                        - name: libcrypt-dev
                        - name: linux-libc-dev
                        - name: libc-dev-bin
                        - name: libc6-dev
                        - name: zlib1g
            - package_name: xtable
              package_path: xtable_1.8-8.tar.gz
              bioconductor: true
        - package_name: Rtsne
          package_path: Rtsne_0.17.tar.gz
          bioconductor: true
          dependencies:
            - package_name: Rcpp
              package_path: Rcpp_1.1.1.tar.gz
              bioconductor: true
        - package_name: scattermore
          package_path: scattermore_1.2.tar.gz
          bioconductor: true
          dependencies:
            - package_name: scales
              package_path: scales_1.4.0.tar.gz
              bioconductor: true
              dependencies:
                - package_name: glue
                  package_path: glue_1.8.0.tar.gz
                  bioconductor: true
                - package_name: lifecycle
                  package_path: lifecycle_1.0.5.tar.gz
                  bioconductor: true
                  dependencies:
                    - package_name: cli
                      package_path: cli_3.6.5.tar.gz
                      bioconductor: true
                - package_name: cli
                  package_path: cli_3.6.5.tar.gz
                  bioconductor: true
                - package_name: RColorBrewer
                  package_path: RColorBrewer_1.1-3.tar.gz
                  bioconductor: true
                - package_name: rlang
                  package_path: rlang_1.1.7.tar.gz
                  bioconductor: true
                - package_name: farver
                  package_path: farver_2.1.2.tar.gz
                  bioconductor: true
                - package_name: viridisLite
                  package_path: viridisLite_0.4.3.tar.gz
                  bioconductor: true
                - package_name: labeling
                  package_path: labeling_0.4.3.tar.gz
                  bioconductor: true
            - package_name: ggplot2
              package_path: ggplot2_4.0.2.tar.gz
              bioconductor: true
              dependencies:
                - package_name: lifecycle
                  package_path: lifecycle_1.0.5.tar.gz
                  bioconductor: true
                  dependencies:
                    - package_name: cli
                      package_path: cli_3.6.5.tar.gz
                      bioconductor: true
                - package_name: gtable
                  package_path: gtable_0.3.6.tar.gz
                  bioconductor: true
                  dependencies:
                    - package_name: glue
                      package_path: glue_1.8.0.tar.gz
                      bioconductor: true
                    - package_name: rlang
                      package_path: rlang_1.1.7.tar.gz
                      bioconductor: true
                    - package_name: lifecycle
                      package_path: lifecycle_1.0.5.tar.gz
                      bioconductor: true
                - package_name: rlang
                  package_path: rlang_1.1.7.tar.gz
                  bioconductor: true
                - package_name: S7
                  package_path: S7_0.2.1.tar.gz
                  bioconductor: true
                - package_name: isoband
                  package_path: isoband_0.3.0.tar.gz
                  bioconductor: true
                  dependencies:
                    - package_name: rlang
                      package_path: rlang_1.1.7.tar.gz
                      bioconductor: true
                    - package_name: cpp11
                      package_path: cpp11_0.5.3.tar.gz
                      bioconductor: true
                - package_name: glue
                  package_path: glue_1.8.0.tar.gz
                  bioconductor: true
                - package_name: cpp11
                  package_path: cpp11_0.5.3.tar.gz
                  bioconductor: true
                - package_name: vctrs
                  package_path: vctrs_0.7.1.tar.gz
                  bioconductor: true
                  dependencies:
                    - package_name: rlang
                      package_path: rlang_1.1.7.tar.gz
                      bioconductor: true
                    - package_name: cli
                      package_path: cli_3.6.5.tar.gz
                      bioconductor: true
                    - package_name: lifecycle
                      package_path: lifecycle_1.0.5.tar.gz
                      bioconductor: true
                      dependencies:
                        - package_name: cli
                          package_path: cli_3.6.5.tar.gz
                          bioconductor: true
                    - package_name: glue
                      package_path: glue_1.8.0.tar.gz
                      bioconductor: true
                - package_name: farver
                  package_path: farver_2.1.2.tar.gz
                  bioconductor: true
                - package_name: RColorBrewer
                  package_path: RColorBrewer_1.1-3.tar.gz
                  bioconductor: true
                - package_name: labeling
                  package_path: labeling_0.4.3.tar.gz
                  bioconductor: true
                - package_name: scales
                  package_path: scales_1.4.0.tar.gz
                  bioconductor: true
                  dependencies:
                    - package_name: glue
                      package_path: glue_1.8.0.tar.gz
                      bioconductor: true
                    - package_name: lifecycle
                      package_path: lifecycle_1.0.5.tar.gz
                      bioconductor: true
                      dependencies:
                        - package_name: cli
                          package_path: cli_3.6.5.tar.gz
                          bioconductor: true
                    - package_name: cli
                      package_path: cli_3.6.5.tar.gz
                      bioconductor: true
                    - package_name: RColorBrewer
                      package_path: RColorBrewer_1.1-3.tar.gz
                      bioconductor: true
                    - package_name: rlang
                      package_path: rlang_1.1.7.tar.gz
                      bioconductor: true
                    - package_name: farver
                      package_path: farver_2.1.2.tar.gz
                      bioconductor: true
                    - package_name: viridisLite
                      package_path: viridisLite_0.4.3.tar.gz
                      bioconductor: true
                    - package_name: labeling
                      package_path: labeling_0.4.3.tar.gz
                      bioconductor: true
                - package_name: R6
                  package_path: R6_2.6.1.tar.gz
                  bioconductor: true
                - package_name: viridisLite
                  package_path: viridisLite_0.4.3.tar.gz
                  bioconductor: true
            - package_name: cli
              package_path: cli_3.6.5.tar.gz
              bioconductor: true
            - package_name: gtable
              package_path: gtable_0.3.6.tar.gz
              bioconductor: true
              dependencies:
                - package_name: rlang
                  package_path: rlang_1.1.7.tar.gz
                  bioconductor: true
            - package_name: isoband
              package_path: isoband_0.3.0.tar.gz
              bioconductor: true
              dependencies:
                - package_name: rlang
                  package_path: rlang_1.1.7.tar.gz
                  bioconductor: true
                - package_name: cpp11
                  package_path: cpp11_0.5.3.tar.gz
                  bioconductor: true
            - package_name: lifecycle
              package_path: lifecycle_1.0.5.tar.gz
              bioconductor: true
              dependencies:
                - package_name: cli
                  package_path: cli_3.6.5.tar.gz
                  bioconductor: true
            - package_name: rlang
              package_path: rlang_1.1.7.tar.gz
              bioconductor: true
            - package_name: S7
              package_path: S7_0.2.1.tar.gz
              bioconductor: true
            - package_name: vctrs
              package_path: vctrs_0.7.1.tar.gz
              bioconductor: true
              dependencies:
                - package_name: rlang
                  package_path: rlang_1.1.7.tar.gz
                  bioconductor: true
                - package_name: glue
                  package_path: glue_1.8.0.tar.gz
                  bioconductor: true
                - package_name: cli
                  package_path: cli_3.6.5.tar.gz
                  bioconductor: true
                - package_name: lifecycle
                  package_path: lifecycle_1.0.5.tar.gz
                  bioconductor: true
                  dependencies:
                    - package_name: cli
                      package_path: cli_3.6.5.tar.gz
                      bioconductor: true
            - package_name: withr
              package_path: withr_3.0.2.tar.gz
              bioconductor: true
            - package_name: farver
              package_path: farver_2.1.2.tar.gz
              bioconductor: true
            - package_name: glue
              package_path: glue_1.8.0.tar.gz
              bioconductor: true
            - package_name: labeling
              package_path: labeling_0.4.3.tar.gz
              bioconductor: true
            - package_name: R6
              package_path: R6_2.6.1.tar.gz
              bioconductor: true
            - package_name: RColorBrewer
              package_path: RColorBrewer_1.1-3.tar.gz
              bioconductor: true
            - package_name: viridisLite
              package_path: viridisLite_0.4.3.tar.gz
              bioconductor: true
            - package_name: cpp11
              package_path: cpp11_0.5.3.tar.gz
              bioconductor: true
        - package_name: tibble
          package_path: tibble_3.3.1.tar.gz
          bioconductor: true
          dependencies:
            - package_name: magrittr
              package_path: magrittr_2.0.4.tar.gz
              bioconductor: true
            - package_name: rlang
              package_path: rlang_1.1.7.tar.gz
              bioconductor: true
            - package_name: pkgconfig
              package_path: pkgconfig_2.0.3.tar.gz
              bioconductor: true
            - package_name: lifecycle
              package_path: lifecycle_1.0.5.tar.gz
              bioconductor: true
            - package_name: glue
              package_path: glue_1.8.0.tar.gz
              bioconductor: true
            - package_name: pillar
              package_path: pillar_1.11.1.tar.gz
              bioconductor: true
              dependencies:
                - package_name: rlang
                  package_path: rlang_1.1.7.tar.gz
                  bioconductor: true
                - package_name: glue
                  package_path: glue_1.8.0.tar.gz
                  bioconductor: true
                - package_name: utf8
                  package_path: utf8_1.2.6.tar.gz
                  bioconductor: true
                - package_name: lifecycle
                  package_path: lifecycle_1.0.5.tar.gz
                  bioconductor: true
                  dependencies:
                    - package_name: rlang
                      package_path: rlang_1.1.7.tar.gz
                      bioconductor: true
                    - package_name: cli
                      package_path: cli_3.6.5.tar.gz
                      bioconductor: true
            - package_name: utf8
              package_path: utf8_1.2.6.tar.gz
              bioconductor: true
            - package_name: vctrs
              package_path: vctrs_0.7.1.tar.gz
              bioconductor: true
              dependencies:
                - package_name: rlang
                  package_path: rlang_1.1.7.tar.gz
                  bioconductor: true
                - package_name: glue
                  package_path: glue_1.8.0.tar.gz
                  bioconductor: true
                - package_name: cli
                  package_path: cli_3.6.5.tar.gz
                  bioconductor: true
                - package_name: lifecycle
                  package_path: lifecycle_1.0.5.tar.gz
                  bioconductor: true
                  dependencies:
                    - package_name: cli
                      package_path: cli_3.6.5.tar.gz
                      bioconductor: true
        - package_name: sctransform
          package_path: sctransform_0.4.3.tar.gz
          bioconductor: true
          dependencies:
            - package_name: magrittr
              package_path: magrittr_2.0.4.tar.gz
              bioconductor: true
            - package_name: dplyr
              package_path: dplyr_1.2.0.tar.gz
              bioconductor: true
              dependencies:
                - package_name: lifecycle
                  package_path: lifecycle_1.0.5.tar.gz
                  bioconductor: true
                  dependencies:
                    - package_name: cli
                      package_path: cli_3.6.5.tar.gz
                      bioconductor: true
                - package_name: generics
                  package_path: generics_0.1.4.tar.gz
                  bioconductor: true
                - package_name: cli
                  package_path: cli_3.6.5.tar.gz
                  bioconductor: true
                - package_name: pillar
                  package_path: pillar_1.11.1.tar.gz
                  bioconductor: true
                  dependencies:
                    - package_name: rlang
                      package_path: rlang_1.1.7.tar.gz
                      bioconductor: true
                    - package_name: glue
                      package_path: glue_1.8.0.tar.gz
                      bioconductor: true
                    - package_name: utf8
                      package_path: utf8_1.2.6.tar.gz
                      bioconductor: true
                    - package_name: lifecycle
                      package_path: lifecycle_1.0.5.tar.gz
                      bioconductor: true
                      dependencies:
                        - package_name: rlang
                          package_path: rlang_1.1.7.tar.gz
                          bioconductor: true
                        - package_name: cli
                          package_path: cli_3.6.5.tar.gz
                          bioconductor: true
                - package_name: R6
                  package_path: R6_2.6.1.tar.gz
                  bioconductor: true
                - package_name: rlang
                  package_path: rlang_1.1.7.tar.gz
                  bioconductor: true
                - package_name: tibble
                  package_path: tibble_3.3.1.tar.gz
                  bioconductor: true
                  dependencies:
                    - package_name: magrittr
                      package_path: magrittr_2.0.4.tar.gz
                      bioconductor: true
                    - package_name: rlang
                      package_path: rlang_1.1.7.tar.gz
                      bioconductor: true
                    - package_name: pkgconfig
                      package_path: pkgconfig_2.0.3.tar.gz
                      bioconductor: true
                    - package_name: lifecycle
                      package_path: lifecycle_1.0.5.tar.gz
                      bioconductor: true
                    - package_name: glue
                      package_path: glue_1.8.0.tar.gz
                      bioconductor: true
                    - package_name: pillar
                      package_path: pillar_1.11.1.tar.gz
                      bioconductor: true
                      dependencies:
                        - package_name: rlang
                          package_path: rlang_1.1.7.tar.gz
                          bioconductor: true
                        - package_name: glue
                          package_path: glue_1.8.0.tar.gz
                          bioconductor: true
                        - package_name: utf8
                          package_path: utf8_1.2.6.tar.gz
                          bioconductor: true
                        - package_name: lifecycle
                          package_path: lifecycle_1.0.5.tar.gz
                          bioconductor: true
                          dependencies:
                            - package_name: rlang
                              package_path: rlang_1.1.7.tar.gz
                              bioconductor: true
                            - package_name: cli
                              package_path: cli_3.6.5.tar.gz
                              bioconductor: true
                    - package_name: utf8
                      package_path: utf8_1.2.6.tar.gz
                      bioconductor: true
                    - package_name: vctrs
                      package_path: vctrs_0.7.1.tar.gz
                      bioconductor: true
                      dependencies:
                        - package_name: rlang
                          package_path: rlang_1.1.7.tar.gz
                          bioconductor: true
                        - package_name: glue
                          package_path: glue_1.8.0.tar.gz
                          bioconductor: true
                        - package_name: cli
                          package_path: cli_3.6.5.tar.gz
                          bioconductor: true
                        - package_name: lifecycle
                          package_path: lifecycle_1.0.5.tar.gz
                          bioconductor: true
                          dependencies:
                            - package_name: cli
                              package_path: cli_3.6.5.tar.gz
                              bioconductor: true
                - package_name: glue
                  package_path: glue_1.8.0.tar.gz
                  bioconductor: true
                - package_name: magrittr
                  package_path: magrittr_2.0.4.tar.gz
                  bioconductor: true
                - package_name: vctrs
                  package_path: vctrs_0.7.1.tar.gz
                  bioconductor: true
                  dependencies:
                    - package_name: rlang
                      package_path: rlang_1.1.7.tar.gz
                      bioconductor: true
                    - package_name: glue
                      package_path: glue_1.8.0.tar.gz
                      bioconductor: true
                    - package_name: cli
                      package_path: cli_3.6.5.tar.gz
                      bioconductor: true
                    - package_name: lifecycle
                      package_path: lifecycle_1.0.5.tar.gz
                      bioconductor: true
                      dependencies:
                        - package_name: cli
                          package_path: cli_3.6.5.tar.gz
                          bioconductor: true
                - package_name: utf8
                  package_path: utf8_1.2.6.tar.gz
                  bioconductor: true
                - package_name: pkgconfig
                  package_path: pkgconfig_2.0.3.tar.gz
                  bioconductor: true
                - package_name: withr
                  package_path: withr_3.0.2.tar.gz
                  bioconductor: true
            - package_name: future.apply
              package_path: future.apply_1.20.2.tar.gz
              bioconductor: true
              dependencies:
                - package_name: listenv
                  package_path: listenv_0.10.1.tar.gz
                  bioconductor: true
                - package_name: codetools
                  package_path: codetools_0.2-20.tar.gz
                  bioconductor: true
                - package_name: parallelly
                  package_path: parallelly_1.46.1.tar.gz
                  bioconductor: true
                - package_name: future
                  package_path: future_1.69.0.tar.gz
                  bioconductor: true
                  dependencies:
                    - package_name: listenv
                      package_path: listenv_0.10.1.tar.gz
                      bioconductor: true
                    - package_name: globals
                      package_path: globals_0.19.0.tar.gz
                      bioconductor: true
                      dependencies:
                        - package_name: codetools
                          package_path: codetools_0.2-20.tar.gz
                          bioconductor: true
                    - package_name: codetools
                      package_path: codetools_0.2-20.tar.gz
                      bioconductor: true
                    - package_name: digest
                      package_path: digest_0.6.39.tar.gz
                      bioconductor: true
                    - package_name: parallelly
                      package_path: parallelly_1.46.1.tar.gz
                      bioconductor: true
            - package_name: future
              package_path: future_1.69.0.tar.gz
              bioconductor: true
              dependencies:
                - package_name: digest
                  package_path: digest_0.6.39.tar.gz
                  bioconductor: true
                - package_name: codetools
                  package_path: codetools_0.2-20.tar.gz
                  bioconductor: true
            - package_name: parallelly
              package_path: parallelly_1.46.1.tar.gz
              bioconductor: true
            - package_name: ggplot2
              package_path: ggplot2_4.0.2.tar.gz
              bioconductor: true
              dependencies:
                - package_name: lifecycle
                  package_path: lifecycle_1.0.5.tar.gz
                  bioconductor: true
                  dependencies:
                    - package_name: cli
                      package_path: cli_3.6.5.tar.gz
                      bioconductor: true
                - package_name: gtable
                  package_path: gtable_0.3.6.tar.gz
                  bioconductor: true
                  dependencies:
                    - package_name: glue
                      package_path: glue_1.8.0.tar.gz
                      bioconductor: true
                    - package_name: rlang
                      package_path: rlang_1.1.7.tar.gz
                      bioconductor: true
                    - package_name: lifecycle
                      package_path: lifecycle_1.0.5.tar.gz
                      bioconductor: true
                - package_name: rlang
                  package_path: rlang_1.1.7.tar.gz
                  bioconductor: true
                - package_name: S7
                  package_path: S7_0.2.1.tar.gz
                  bioconductor: true
                - package_name: isoband
                  package_path: isoband_0.3.0.tar.gz
                  bioconductor: true
                  dependencies:
                    - package_name: rlang
                      package_path: rlang_1.1.7.tar.gz
                      bioconductor: true
                    - package_name: cpp11
                      package_path: cpp11_0.5.3.tar.gz
                      bioconductor: true
                - package_name: glue
                  package_path: glue_1.8.0.tar.gz
                  bioconductor: true
                - package_name: cpp11
                  package_path: cpp11_0.5.3.tar.gz
                  bioconductor: true
                - package_name: vctrs
                  package_path: vctrs_0.7.1.tar.gz
                  bioconductor: true
                  dependencies:
                    - package_name: rlang
                      package_path: rlang_1.1.7.tar.gz
                      bioconductor: true
                    - package_name: cli
                      package_path: cli_3.6.5.tar.gz
                      bioconductor: true
                    - package_name: lifecycle
                      package_path: lifecycle_1.0.5.tar.gz
                      bioconductor: true
                      dependencies:
                        - package_name: cli
                          package_path: cli_3.6.5.tar.gz
                          bioconductor: true
                    - package_name: glue
                      package_path: glue_1.8.0.tar.gz
                      bioconductor: true
                - package_name: farver
                  package_path: farver_2.1.2.tar.gz
                  bioconductor: true
                - package_name: RColorBrewer
                  package_path: RColorBrewer_1.1-3.tar.gz
                  bioconductor: true
                - package_name: labeling
                  package_path: labeling_0.4.3.tar.gz
                  bioconductor: true
                - package_name: scales
                  package_path: scales_1.4.0.tar.gz
                  bioconductor: true
                  dependencies:
                    - package_name: glue
                      package_path: glue_1.8.0.tar.gz
                      bioconductor: true
                    - package_name: lifecycle
                      package_path: lifecycle_1.0.5.tar.gz
                      bioconductor: true
                      dependencies:
                        - package_name: cli
                          package_path: cli_3.6.5.tar.gz
                          bioconductor: true
                    - package_name: cli
                      package_path: cli_3.6.5.tar.gz
                      bioconductor: true
                    - package_name: RColorBrewer
                      package_path: RColorBrewer_1.1-3.tar.gz
                      bioconductor: true
                    - package_name: rlang
                      package_path: rlang_1.1.7.tar.gz
                      bioconductor: true
                    - package_name: farver
                      package_path: farver_2.1.2.tar.gz
                      bioconductor: true
                    - package_name: viridisLite
                      package_path: viridisLite_0.4.3.tar.gz
                      bioconductor: true
                    - package_name: labeling
                      package_path: labeling_0.4.3.tar.gz
                      bioconductor: true
                - package_name: R6
                  package_path: R6_2.6.1.tar.gz
                  bioconductor: true
                - package_name: viridisLite
                  package_path: viridisLite_0.4.3.tar.gz
                  bioconductor: true
            - package_name: rlang
              package_path: rlang_1.1.7.tar.gz
              bioconductor: true
            - package_name: matrixStats
              package_path: matrixStats_1.5.0.tar.gz
              bioconductor: true
            - package_name: Rcpp
              package_path: Rcpp_1.1.1.tar.gz
              bioconductor: true
            - package_name: cli
              package_path: cli_3.6.5.tar.gz
              bioconductor: true
            - package_name: generics
              package_path: generics_0.1.4.tar.gz
              bioconductor: true
            - package_name: glue
              package_path: glue_1.8.0.tar.gz
              bioconductor: true
            - package_name: lifecycle
              package_path: lifecycle_1.0.5.tar.gz
              bioconductor: true
              dependencies:
                - package_name: cli
                  package_path: cli_3.6.5.tar.gz
                  bioconductor: true
            - package_name: pillar
              package_path: pillar_1.11.1.tar.gz
              bioconductor: true
              dependencies:
                - package_name: rlang
                  package_path: rlang_1.1.7.tar.gz
                  bioconductor: true
                - package_name: glue
                  package_path: glue_1.8.0.tar.gz
                  bioconductor: true
                - package_name: utf8
                  package_path: utf8_1.2.6.tar.gz
                  bioconductor: true
                - package_name: lifecycle
                  package_path: lifecycle_1.0.5.tar.gz
                  bioconductor: true
                  dependencies:
                    - package_name: rlang
                      package_path: rlang_1.1.7.tar.gz
                      bioconductor: true
                    - package_name: cli
                      package_path: cli_3.6.5.tar.gz
                      bioconductor: true
            - package_name: R6
              package_path: R6_2.6.1.tar.gz
              bioconductor: true
            - package_name: tibble
              package_path: tibble_3.3.1.tar.gz
              bioconductor: true
              dependencies:
                - package_name: magrittr
                  package_path: magrittr_2.0.4.tar.gz
                  bioconductor: true
                - package_name: rlang
                  package_path: rlang_1.1.7.tar.gz
                  bioconductor: true
                - package_name: pkgconfig
                  package_path: pkgconfig_2.0.3.tar.gz
                  bioconductor: true
                - package_name: lifecycle
                  package_path: lifecycle_1.0.5.tar.gz
                  bioconductor: true
                - package_name: glue
                  package_path: glue_1.8.0.tar.gz
                  bioconductor: true
                - package_name: pillar
                  package_path: pillar_1.11.1.tar.gz
                  bioconductor: true
                  dependencies:
                    - package_name: rlang
                      package_path: rlang_1.1.7.tar.gz
                      bioconductor: true
                    - package_name: glue
                      package_path: glue_1.8.0.tar.gz
                      bioconductor: true
                    - package_name: utf8
                      package_path: utf8_1.2.6.tar.gz
                      bioconductor: true
                    - package_name: lifecycle
                      package_path: lifecycle_1.0.5.tar.gz
                      bioconductor: true
                      dependencies:
                        - package_name: rlang
                          package_path: rlang_1.1.7.tar.gz
                          bioconductor: true
                        - package_name: cli
                          package_path: cli_3.6.5.tar.gz
                          bioconductor: true
                - package_name: utf8
                  package_path: utf8_1.2.6.tar.gz
                  bioconductor: true
                - package_name: vctrs
                  package_path: vctrs_0.7.1.tar.gz
                  bioconductor: true
                  dependencies:
                    - package_name: rlang
                      package_path: rlang_1.1.7.tar.gz
                      bioconductor: true
                    - package_name: glue
                      package_path: glue_1.8.0.tar.gz
                      bioconductor: true
                    - package_name: cli
                      package_path: cli_3.6.5.tar.gz
                      bioconductor: true
                    - package_name: lifecycle
                      package_path: lifecycle_1.0.5.tar.gz
                      bioconductor: true
                      dependencies:
                        - package_name: cli
                          package_path: cli_3.6.5.tar.gz
                          bioconductor: true
            - package_name: tidyselect
              package_path: tidyselect_1.2.1.tar.gz
              bioconductor: true
              dependencies:
                - package_name: rlang
                  package_path: rlang_1.1.7.tar.gz
                  bioconductor: true
                - package_name: vctrs
                  package_path: vctrs_0.7.1.tar.gz
                  bioconductor: true
                  dependencies:
                    - package_name: rlang
                      package_path: rlang_1.1.7.tar.gz
                      bioconductor: true
                    - package_name: glue
                      package_path: glue_1.8.0.tar.gz
                      bioconductor: true
                    - package_name: cli
                      package_path: cli_3.6.5.tar.gz
                      bioconductor: true
                    - package_name: lifecycle
                      package_path: lifecycle_1.0.5.tar.gz
                      bioconductor: true
                      dependencies:
                        - package_name: cli
                          package_path: cli_3.6.5.tar.gz
                          bioconductor: true
                - package_name: withr
                  package_path: withr_3.0.2.tar.gz
                  bioconductor: true
                - package_name: glue
                  package_path: glue_1.8.0.tar.gz
                  bioconductor: true
                - package_name: lifecycle
                  package_path: lifecycle_1.0.5.tar.gz
                  bioconductor: true
                  dependencies:
                    - package_name: cli
                      package_path: cli_3.6.5.tar.gz
                      bioconductor: true
                - package_name: cli
                  package_path: cli_3.6.5.tar.gz
                  bioconductor: true
            - package_name: vctrs
              package_path: vctrs_0.7.1.tar.gz
              bioconductor: true
              dependencies:
                - package_name: rlang
                  package_path: rlang_1.1.7.tar.gz
                  bioconductor: true
                - package_name: glue
                  package_path: glue_1.8.0.tar.gz
                  bioconductor: true
                - package_name: cli
                  package_path: cli_3.6.5.tar.gz
                  bioconductor: true
                - package_name: lifecycle
                  package_path: lifecycle_1.0.5.tar.gz
                  bioconductor: true
                  dependencies:
                    - package_name: cli
                      package_path: cli_3.6.5.tar.gz
                      bioconductor: true
            - package_name: digest
              package_path: digest_0.6.39.tar.gz
              bioconductor: true
            - package_name: globals
              package_path: globals_0.19.0.tar.gz
              bioconductor: true
              dependencies:
                - package_name: codetools
                  package_path: codetools_0.2-20.tar.gz
                  bioconductor: true
            - package_name: listenv
              package_path: listenv_0.10.1.tar.gz
              bioconductor: true
            - package_name: gtable
              package_path: gtable_0.3.6.tar.gz
              bioconductor: true
              dependencies:
                - package_name: rlang
                  package_path: rlang_1.1.7.tar.gz
                  bioconductor: true
            - package_name: isoband
              package_path: isoband_0.3.0.tar.gz
              bioconductor: true
              dependencies:
                - package_name: rlang
                  package_path: rlang_1.1.7.tar.gz
                  bioconductor: true
                - package_name: cpp11
                  package_path: cpp11_0.5.3.tar.gz
                  bioconductor: true
            - package_name: S7
              package_path: S7_0.2.1.tar.gz
              bioconductor: true
            - package_name: scales
              package_path: scales_1.4.0.tar.gz
              bioconductor: true
              dependencies:
                - package_name: glue
                  package_path: glue_1.8.0.tar.gz
                  bioconductor: true
                - package_name: lifecycle
                  package_path: lifecycle_1.0.5.tar.gz
                  bioconductor: true
                  dependencies:
                    - package_name: cli
                      package_path: cli_3.6.5.tar.gz
                      bioconductor: true
                - package_name: cli
                  package_path: cli_3.6.5.tar.gz
                  bioconductor: true
                - package_name: RColorBrewer
                  package_path: RColorBrewer_1.1-3.tar.gz
                  bioconductor: true
                - package_name: rlang
                  package_path: rlang_1.1.7.tar.gz
                  bioconductor: true
                - package_name: farver
                  package_path: farver_2.1.2.tar.gz
                  bioconductor: true
                - package_name: viridisLite
                  package_path: viridisLite_0.4.3.tar.gz
                  bioconductor: true
                - package_name: labeling
                  package_path: labeling_0.4.3.tar.gz
                  bioconductor: true
            - package_name: gridExtra
              package_path: gridExtra_2.3.tar.gz
              bioconductor: true
              dependencies:
                - package_name: gtable
                  package_path: gtable_0.3.6.tar.gz
                  bioconductor: true
                  dependencies:
                    - package_name: rlang
                      package_path: rlang_1.1.7.tar.gz
                      bioconductor: true
                - package_name: cli
                  package_path: cli_3.6.5.tar.gz
                  bioconductor: true
                - package_name: glue
                  package_path: glue_1.8.0.tar.gz
                  bioconductor: true
                - package_name: lifecycle
                  package_path: lifecycle_1.0.5.tar.gz
                  bioconductor: true
                  dependencies:
                    - package_name: cli
                      package_path: cli_3.6.5.tar.gz
                      bioconductor: true
                - package_name: rlang
                  package_path: rlang_1.1.7.tar.gz
                  bioconductor: true
            - package_name: withr
              package_path: withr_3.0.2.tar.gz
              bioconductor: true
            - package_name: reshape2
              package_path: reshape2_1.4.5.tar.gz
              bioconductor: true
              dependencies:
                - package_name: Rcpp
                  package_path: Rcpp_1.1.1.tar.gz
                  bioconductor: true
                - package_name: stringr
                  package_path: stringr_1.6.0.tar.gz
                  bioconductor: true
                  dependencies:
                    - package_name: magrittr
                      package_path: magrittr_2.0.4.tar.gz
                      bioconductor: true
                    - package_name: glue
                      package_path: glue_1.8.0.tar.gz
                      bioconductor: true
                    - package_name: lifecycle
                      package_path: lifecycle_1.0.5.tar.gz
                      bioconductor: true
                      dependencies:
                        - package_name: cli
                          package_path: cli_3.6.5.tar.gz
                          bioconductor: true
                    - package_name: stringi
                      package_path: stringi_1.8.7.tar.gz
                      bioconductor: true
                      external_dependencies:
                        apt:
                            - name: libicu-dev
                              dependencies:
                                - name: libc-dev
                                  optional: true
                                - name: libc6
                                - name: libcrypt1
                                - name: gcc-14-base
                                - name: rpcsvc-proto
                                - name: libcrypt-dev
                                - name: linux-libc-dev
                                - name: libc-dev-bin
                                - name: libgcc-s1
                                - name: libstdc++6
                                - name: libc6-dev
                                - name: libicu74
                                - name: icu-devtools
                    - package_name: rlang
                      package_path: rlang_1.1.7.tar.gz
                      bioconductor: true
                - package_name: cli
                  package_path: cli_3.6.5.tar.gz
                  bioconductor: true
                - package_name: glue
                  package_path: glue_1.8.0.tar.gz
                  bioconductor: true
                - package_name: lifecycle
                  package_path: lifecycle_1.0.5.tar.gz
                  bioconductor: true
                  dependencies:
                    - package_name: cli
                      package_path: cli_3.6.5.tar.gz
                      bioconductor: true
                - package_name: magrittr
                  package_path: magrittr_2.0.4.tar.gz
                  bioconductor: true
                - package_name: rlang
                  package_path: rlang_1.1.7.tar.gz
                  bioconductor: true
                - package_name: stringi
                  package_path: stringi_1.8.7.tar.gz
                  bioconductor: true
                  external_dependencies:
                    apt:
                        - name: libicu-dev
                          dependencies:
                            - name: libc-dev
                              optional: true
                            - name: libc6
                            - name: libcrypt1
                            - name: gcc-14-base
                            - name: rpcsvc-proto
                            - name: libcrypt-dev
                            - name: linux-libc-dev
                            - name: libc-dev-bin
                            - name: libgcc-s1
                            - name: libstdc++6
                            - name: libc6-dev
                            - name: libicu74
                            - name: icu-devtools
                - package_name: vctrs
                  package_path: vctrs_0.7.1.tar.gz
                  bioconductor: true
                  dependencies:
                    - package_name: rlang
                      package_path: rlang_1.1.7.tar.gz
                      bioconductor: true
                    - package_name: glue
                      package_path: glue_1.8.0.tar.gz
                      bioconductor: true
                    - package_name: cli
                      package_path: cli_3.6.5.tar.gz
                      bioconductor: true
                    - package_name: lifecycle
                      package_path: lifecycle_1.0.5.tar.gz
                      bioconductor: true
                      dependencies:
                        - package_name: cli
                          package_path: cli_3.6.5.tar.gz
                          bioconductor: true
                - package_name: plyr
                  package_path: plyr_1.8.9.tar.gz
                  bioconductor: true
                  dependencies:
                    - package_name: Rcpp
                      package_path: Rcpp_1.1.1.tar.gz
                      bioconductor: true
            - package_name: plyr
              package_path: plyr_1.8.9.tar.gz
              bioconductor: true
              dependencies:
                - package_name: Rcpp
                  package_path: Rcpp_1.1.1.tar.gz
                  bioconductor: true
            - package_name: stringr
              package_path: stringr_1.6.0.tar.gz
              bioconductor: true
              dependencies:
                - package_name: magrittr
                  package_path: magrittr_2.0.4.tar.gz
                  bioconductor: true
                - package_name: glue
                  package_path: glue_1.8.0.tar.gz
                  bioconductor: true
                - package_name: lifecycle
                  package_path: lifecycle_1.0.5.tar.gz
                  bioconductor: true
                  dependencies:
                    - package_name: cli
                      package_path: cli_3.6.5.tar.gz
                      bioconductor: true
                - package_name: stringi
                  package_path: stringi_1.8.7.tar.gz
                  bioconductor: true
                  external_dependencies:
                    apt:
                        - name: libicu-dev
                          dependencies:
                            - name: libc-dev
                              optional: true
                            - name: libc6
                            - name: libcrypt1
                            - name: gcc-14-base
                            - name: rpcsvc-proto
                            - name: libcrypt-dev
                            - name: linux-libc-dev
                            - name: libc-dev-bin
                            - name: libgcc-s1
                            - name: libstdc++6
                            - name: libc6-dev
                            - name: libicu74
                            - name: icu-devtools
                - package_name: rlang
                  package_path: rlang_1.1.7.tar.gz
                  bioconductor: true
            - package_name: codetools
              package_path: codetools_0.2-20.tar.gz
              bioconductor: true
            - package_name: cpp11
              package_path: cpp11_0.5.3.tar.gz
              bioconductor: true
            - package_name: utf8
              package_path: utf8_1.2.6.tar.gz
              bioconductor: true
            - package_name: farver
              package_path: farver_2.1.2.tar.gz
              bioconductor: true
            - package_name: labeling
              package_path: labeling_0.4.3.tar.gz
              bioconductor: true
            - package_name: RColorBrewer
              package_path: RColorBrewer_1.1-3.tar.gz
              bioconductor: true
            - package_name: viridisLite
              package_path: viridisLite_0.4.3.tar.gz
              bioconductor: true
            - package_name: stringi
              package_path: stringi_1.8.7.tar.gz
              bioconductor: true
              external_dependencies:
                apt:
                    - name: libicu-dev
                      dependencies:
                        - name: libc-dev
                          optional: true
                        - name: libc6
                        - name: libcrypt1
                        - name: gcc-14-base
                        - name: rpcsvc-proto
                        - name: libcrypt-dev
                        - name: linux-libc-dev
                        - name: libc-dev-bin
                        - name: libgcc-s1
                        - name: libstdc++6
                        - name: libc6-dev
                        - name: libicu74
                        - name: icu-devtools
            - package_name: pkgconfig
              package_path: pkgconfig_2.0.3.tar.gz
              bioconductor: true
            - package_name: RcppArmadillo
              package_path: RcppArmadillo_15.2.3-1.tar.gz
              bioconductor: true
              dependencies:
                - package_name: Rcpp
                  package_path: Rcpp_1.1.1.tar.gz
                  bioconductor: true
              external_dependencies:
                apt:
                    - name: libblas-dev
                      dependencies:
                        - name: gcc-14-base
                        - name: libc6
                        - name: libgcc-s1
                        - name: libblas3
                    - name: liblapack-dev
                      dependencies:
                        - name: libblas.so.3
                          optional: true
                        - name: libblas.so
                          optional: true
                        - name: gcc-14-base
                        - name: libc6
                        - name: libgomp1
                        - name: libgcc-s1
                        - name: libgfortran5
                        - name: libopenblas0-serial
                        - name: libopenblas0-pthread
                        - name: libopenblas0-openmp
                        - name: libblis4-serial
                        - name: libblis4-pthread
                        - name: libblis4-openmp
                        - name: libatlas3-base
                        - name: libblas3
                        - name: libopenblas-serial-dev
                        - name: libopenblas-pthread-dev
                        - name: libopenblas-openmp-dev
                        - name: libblis-serial-dev
                        - name: libblis-pthread-dev
                        - name: libblis-openmp-dev
                        - name: libatlas-base-dev
                        - name: libblas-dev
                        - name: liblapack3
                    - name: r-base-dev
        - package_name: RcppEigen
          package_path: RcppEigen_0.3.4.0.2.tar.gz
          bioconductor: true
          dependencies:
            - package_name: Rcpp
              package_path: Rcpp_1.1.1.tar.gz
              bioconductor: true
        - package_name: spatstat.geom
          package_path: spatstat.geom_3.7-0.tar.gz
          bioconductor: true
          dependencies:
            - package_name: spatstat.univar
              package_path: spatstat.univar_3.1-6.tar.gz
              bioconductor: true
              dependencies:
                - package_name: spatstat.utils
                  package_path: spatstat.utils_3.2-2.tar.gz
                  bioconductor: true
            - package_name: deldir
              package_path: deldir_2.0-4.tar.gz
              bioconductor: true
            - package_name: spatstat.data
              package_path: spatstat.data_3.1-9.tar.gz
              bioconductor: true
              dependencies:
                - package_name: spatstat.utils
                  package_path: spatstat.utils_3.2-2.tar.gz
                  bioconductor: true
            - package_name: polyclip
              package_path: polyclip_1.10-7.tar.gz
              bioconductor: true
        - package_name: gtable
          package_path: gtable_0.3.6.tar.gz
          bioconductor: true
          dependencies:
            - package_name: rlang
              package_path: rlang_1.1.7.tar.gz
              bioconductor: true
        - package_name: data.table
          package_path: data.table_1.18.2.1.tar.gz
          bioconductor: true
        - package_name: stringr
          package_path: stringr_1.6.0.tar.gz
          bioconductor: true
          dependencies:
            - package_name: magrittr
              package_path: magrittr_2.0.4.tar.gz
              bioconductor: true
            - package_name: glue
              package_path: glue_1.8.0.tar.gz
              bioconductor: true
            - package_name: lifecycle
              package_path: lifecycle_1.0.5.tar.gz
              bioconductor: true
              dependencies:
                - package_name: cli
                  package_path: cli_3.6.5.tar.gz
                  bioconductor: true
            - package_name: stringi
              package_path: stringi_1.8.7.tar.gz
              bioconductor: true
              external_dependencies:
                apt:
                    - name: libicu-dev
                      dependencies:
                        - name: libc-dev
                          optional: true
                        - name: libc6
                        - name: libcrypt1
                        - name: gcc-14-base
                        - name: rpcsvc-proto
                        - name: libcrypt-dev
                        - name: linux-libc-dev
                        - name: libc-dev-bin
                        - name: libgcc-s1
                        - name: libstdc++6
                        - name: libc6-dev
                        - name: libicu74
                        - name: icu-devtools
            - package_name: rlang
              package_path: rlang_1.1.7.tar.gz
              bioconductor: true
        - package_name: survival
          package_path: survival_3.8-6.tar.gz
          bioconductor: true
        - package_name: digest
          package_path: digest_0.6.39.tar.gz
          bioconductor: true
        - package_name: globals
          package_path: globals_0.19.0.tar.gz
          bioconductor: true
          dependencies:
            - package_name: codetools
              package_path: codetools_0.2-20.tar.gz
              bioconductor: true
        - package_name: listenv
          package_path: listenv_0.10.1.tar.gz
          bioconductor: true
        - package_name: parallelly
          package_path: parallelly_1.46.1.tar.gz
          bioconductor: true
        - package_name: cli
          package_path: cli_3.6.5.tar.gz
          bioconductor: true
        - package_name: isoband
          package_path: isoband_0.3.0.tar.gz
          bioconductor: true
          dependencies:
            - package_name: rlang
              package_path: rlang_1.1.7.tar.gz
              bioconductor: true
            - package_name: cpp11
              package_path: cpp11_0.5.3.tar.gz
              bioconductor: true
        - package_name: S7
          package_path: S7_0.2.1.tar.gz
          bioconductor: true
        - package_name: vctrs
          package_path: vctrs_0.7.1.tar.gz
          bioconductor: true
          dependencies:
            - package_name: rlang
              package_path: rlang_1.1.7.tar.gz
              bioconductor: true
            - package_name: glue
              package_path: glue_1.8.0.tar.gz
              bioconductor: true
            - package_name: cli
              package_path: cli_3.6.5.tar.gz
              bioconductor: true
            - package_name: lifecycle
              package_path: lifecycle_1.0.5.tar.gz
              bioconductor: true
              dependencies:
                - package_name: cli
                  package_path: cli_3.6.5.tar.gz
                  bioconductor: true
        - package_name: withr
          package_path: withr_3.0.2.tar.gz
          bioconductor: true
        - package_name: curl
          package_path: curl_7.0.0.tar.gz
          bioconductor: true
          external_dependencies:
            apt:
                - name: libcurl4-openssl-dev
                  dependencies:
                    - name: libhogweed6
                      optional: true
                    - name: libgnutls30
                      optional: true
                    - name: libnettle8
                      optional: true
                    - name: libc6
                    - name: libdb5.3t64
                    - name: libffi8
                    - name: libsasl2-modules-db
                    - name: libtasn1-6
                    - name: libp11-kit0
                    - name: libkeyutils1
                    - name: gcc-14-base
                    - name: libnettle8t64
                    - name: libgmp10
                    - name: libhogweed6t64
                    - name: libssl3t64
                    - name: libsasl2-2
                    - name: libunistring5
                    - name: libidn2-0
                    - name: libgnutls30t64
                    - name: libkrb5support0
                    - name: libk5crypto3
                    - name: libcom-err2
                    - name: libkrb5-3
                    - name: libgcc-s1
                    - name: zlib1g
                    - name: libzstd1
                    - name: libgssapi-krb5-2
                    - name: libssh-4
                    - name: librtmp1
                    - name: libpsl5t64
                    - name: libnghttp2-14
                    - name: libldap2
                    - name: libbrotli1
                    - name: libcurl4t64
        - package_name: mime
          package_path: mime_0.13.tar.gz
          bioconductor: true
        - package_name: openssl
          package_path: openssl_2.3.5.tar.gz
          bioconductor: true
          dependencies:
            - package_name: sys
              package_path: sys_3.4.3.tar.gz
              bioconductor: true
            - package_name: askpass
              package_path: askpass_1.2.1.tar.gz
              bioconductor: true
              dependencies:
                - package_name: sys
                  package_path: sys_3.4.3.tar.gz
                  bioconductor: true
          external_dependencies:
            apt:
                - name: libssl-dev
                  dependencies:
                    - name: gcc-14-base
                    - name: libc6
                    - name: libgcc-s1
                    - name: libssl3t64
        - package_name: R6
          package_path: R6_2.6.1.tar.gz
          bioconductor: true
        - package_name: magrittr
          package_path: magrittr_2.0.4.tar.gz
          bioconductor: true
        - package_name: pkgconfig
          package_path: pkgconfig_2.0.3.tar.gz
          bioconductor: true
        - package_name: cpp11
          package_path: cpp11_0.5.3.tar.gz
          bioconductor: true
        - package_name: zoo
          package_path: zoo_1.8-15.tar.gz
          bioconductor: true
          dependencies:
            - package_name: lattice
              package_path: lattice_0.22-9.tar.gz
              bioconductor: true
        - package_name: htmltools
          package_path: htmltools_0.5.9.tar.gz
          bioconductor: true
          dependencies:
            - package_name: digest
              package_path: digest_0.6.39.tar.gz
              bioconductor: true
            - package_name: rlang
              package_path: rlang_1.1.7.tar.gz
              bioconductor: true
            - package_name: base64enc
              package_path: base64enc_0.1-6.tar.gz
              bioconductor: true
            - package_name: fastmap
              package_path: fastmap_1.2.0.tar.gz
              bioconductor: true
        - package_name: farver
          package_path: farver_2.1.2.tar.gz
          bioconductor: true
        - package_name: viridisLite
          package_path: viridisLite_0.4.3.tar.gz
          bioconductor: true
        - package_name: base64enc
          package_path: base64enc_0.1-6.tar.gz
          bioconductor: true
        - package_name: htmlwidgets
          package_path: htmlwidgets_1.6.4.tar.gz
          bioconductor: true
          dependencies:
            - package_name: jsonlite
              package_path: jsonlite_2.0.0.tar.gz
              bioconductor: true
            - package_name: htmltools
              package_path: htmltools_0.5.9.tar.gz
              bioconductor: true
              dependencies:
                - package_name: digest
                  package_path: digest_0.6.39.tar.gz
                  bioconductor: true
                - package_name: rlang
                  package_path: rlang_1.1.7.tar.gz
                  bioconductor: true
                - package_name: fastmap
                  package_path: fastmap_1.2.0.tar.gz
                  bioconductor: true
                - package_name: base64enc
                  package_path: base64enc_0.1-6.tar.gz
                  bioconductor: true
            - package_name: base64enc
              package_path: base64enc_0.1-6.tar.gz
              bioconductor: true
            - package_name: digest
              package_path: digest_0.6.39.tar.gz
              bioconductor: true
            - package_name: fastmap
              package_path: fastmap_1.2.0.tar.gz
              bioconductor: true
            - package_name: rlang
              package_path: rlang_1.1.7.tar.gz
              bioconductor: true
            - package_name: yaml
              package_path: yaml_2.3.12.tar.gz
              bioconductor: true
            - package_name: highr
              package_path: highr_0.12.tar.gz
              bioconductor: true
              dependencies:
                - package_name: xfun
                  package_path: xfun_0.56.tar.gz
                  bioconductor: true
            - package_name: fontawesome
              package_path: fontawesome_0.5.3.tar.gz
              bioconductor: true
              dependencies:
                - package_name: digest
                  package_path: digest_0.6.39.tar.gz
                  bioconductor: true
                - package_name: fastmap
                  package_path: fastmap_1.2.0.tar.gz
                  bioconductor: true
                - package_name: htmltools
                  package_path: htmltools_0.5.9.tar.gz
                  bioconductor: true
                  dependencies:
                    - package_name: digest
                      package_path: digest_0.6.39.tar.gz
                      bioconductor: true
                    - package_name: rlang
                      package_path: rlang_1.1.7.tar.gz
                      bioconductor: true
                    - package_name: base64enc
                      package_path: base64enc_0.1-6.tar.gz
                      bioconductor: true
                    - package_name: fastmap
                      package_path: fastmap_1.2.0.tar.gz
                      bioconductor: true
                - package_name: rlang
                  package_path: rlang_1.1.7.tar.gz
                  bioconductor: true
                - package_name: base64enc
                  package_path: base64enc_0.1-6.tar.gz
                  bioconductor: true
            - package_name: jquerylib
              package_path: jquerylib_0.1.4.tar.gz
              bioconductor: true
              dependencies:
                - package_name: fastmap
                  package_path: fastmap_1.2.0.tar.gz
                  bioconductor: true
                - package_name: base64enc
                  package_path: base64enc_0.1-6.tar.gz
                  bioconductor: true
                - package_name: rlang
                  package_path: rlang_1.1.7.tar.gz
                  bioconductor: true
                - package_name: htmltools
                  package_path: htmltools_0.5.9.tar.gz
                  bioconductor: true
                  dependencies:
                    - package_name: digest
                      package_path: digest_0.6.39.tar.gz
                      bioconductor: true
                    - package_name: rlang
                      package_path: rlang_1.1.7.tar.gz
                      bioconductor: true
                    - package_name: base64enc
                      package_path: base64enc_0.1-6.tar.gz
                      bioconductor: true
                    - package_name: fastmap
                      package_path: fastmap_1.2.0.tar.gz
                      bioconductor: true
                - package_name: digest
                  package_path: digest_0.6.39.tar.gz
                  bioconductor: true
            - package_name: knitr
              package_path: knitr_1.51.tar.gz
              bioconductor: true
              dependencies:
                - package_name: yaml
                  package_path: yaml_2.3.12.tar.gz
                  bioconductor: true
                - package_name: evaluate
                  package_path: evaluate_1.0.5.tar.gz
                  bioconductor: true
                - package_name: xfun
                  package_path: xfun_0.56.tar.gz
                  bioconductor: true
                - package_name: highr
                  package_path: highr_0.12.tar.gz
                  bioconductor: true
                  dependencies:
                    - package_name: xfun
                      package_path: xfun_0.56.tar.gz
                      bioconductor: true
            - package_name: cachem
              package_path: cachem_1.1.0.tar.gz
              bioconductor: true
              dependencies:
                - package_name: rlang
                  package_path: rlang_1.1.7.tar.gz
                  bioconductor: true
                - package_name: fastmap
                  package_path: fastmap_1.2.0.tar.gz
                  bioconductor: true
            - package_name: lifecycle
              package_path: lifecycle_1.0.5.tar.gz
              bioconductor: true
              dependencies:
                - package_name: cli
                  package_path: cli_3.6.5.tar.gz
                  bioconductor: true
            - package_name: memoise
              package_path: memoise_2.0.1.tar.gz
              bioconductor: true
              dependencies:
                - package_name: cachem
                  package_path: cachem_1.1.0.tar.gz
                  bioconductor: true
                  dependencies:
                    - package_name: rlang
                      package_path: rlang_1.1.7.tar.gz
                      bioconductor: true
                    - package_name: fastmap
                      package_path: fastmap_1.2.0.tar.gz
                      bioconductor: true
                - package_name: fastmap
                  package_path: fastmap_1.2.0.tar.gz
                  bioconductor: true
                - package_name: rlang
                  package_path: rlang_1.1.7.tar.gz
                  bioconductor: true
            - package_name: mime
              package_path: mime_0.13.tar.gz
              bioconductor: true
            - package_name: sass
              package_path: sass_0.4.10.tar.gz
              bioconductor: true
              dependencies:
                - package_name: R6
                  package_path: R6_2.6.1.tar.gz
                  bioconductor: true
                - package_name: htmltools
                  package_path: htmltools_0.5.9.tar.gz
                  bioconductor: true
                  dependencies:
                    - package_name: digest
                      package_path: digest_0.6.39.tar.gz
                      bioconductor: true
                    - package_name: rlang
                      package_path: rlang_1.1.7.tar.gz
                      bioconductor: true
                    - package_name: base64enc
                      package_path: base64enc_0.1-6.tar.gz
                      bioconductor: true
                    - package_name: fastmap
                      package_path: fastmap_1.2.0.tar.gz
                      bioconductor: true
                - package_name: rlang
                  package_path: rlang_1.1.7.tar.gz
                  bioconductor: true
                - package_name: base64enc
                  package_path: base64enc_0.1-6.tar.gz
                  bioconductor: true
                - package_name: digest
                  package_path: digest_0.6.39.tar.gz
                  bioconductor: true
                - package_name: fastmap
                  package_path: fastmap_1.2.0.tar.gz
                  bioconductor: true
                - package_name: rappdirs
                  package_path: rappdirs_0.3.4.tar.gz
                  bioconductor: true
              external_dependencies:
                apt:
                    - name: make
                      dependencies:
                        - name: gcc-14-base
                        - name: libc6
                        - name: libgcc-s1
            - package_name: cli
              package_path: cli_3.6.5.tar.gz
              bioconductor: true
            - package_name: tinytex
              package_path: tinytex_0.58.tar.gz
              bioconductor: true
              dependencies:
                - package_name: xfun
                  package_path: xfun_0.56.tar.gz
                  bioconductor: true
            - package_name: R6
              package_path: R6_2.6.1.tar.gz
              bioconductor: true
            - package_name: rappdirs
              package_path: rappdirs_0.3.4.tar.gz
              bioconductor: true
            - package_name: rmarkdown
              package_path: rmarkdown_2.30.tar.gz
              bioconductor: true
              dependencies:
                - package_name: htmltools
                  package_path: htmltools_0.5.9.tar.gz
                  bioconductor: true
                  dependencies:
                    - package_name: digest
                      package_path: digest_0.6.39.tar.gz
                      bioconductor: true
                    - package_name: rlang
                      package_path: rlang_1.1.7.tar.gz
                      bioconductor: true
                    - package_name: base64enc
                      package_path: base64enc_0.1-6.tar.gz
                      bioconductor: true
                    - package_name: fastmap
                      package_path: fastmap_1.2.0.tar.gz
                      bioconductor: true
                - package_name: fontawesome
                  package_path: fontawesome_0.5.3.tar.gz
                  bioconductor: true
                  dependencies:
                    - package_name: digest
                      package_path: digest_0.6.39.tar.gz
                      bioconductor: true
                    - package_name: fastmap
                      package_path: fastmap_1.2.0.tar.gz
                      bioconductor: true
                    - package_name: htmltools
                      package_path: htmltools_0.5.9.tar.gz
                      bioconductor: true
                      dependencies:
                        - package_name: digest
                          package_path: digest_0.6.39.tar.gz
                          bioconductor: true
                        - package_name: rlang
                          package_path: rlang_1.1.7.tar.gz
                          bioconductor: true
                        - package_name: base64enc
                          package_path: base64enc_0.1-6.tar.gz
                          bioconductor: true
                        - package_name: fastmap
                          package_path: fastmap_1.2.0.tar.gz
                          bioconductor: true
                    - package_name: rlang
                      package_path: rlang_1.1.7.tar.gz
                      bioconductor: true
                    - package_name: base64enc
                      package_path: base64enc_0.1-6.tar.gz
                      bioconductor: true
                - package_name: jsonlite
                  package_path: jsonlite_2.0.0.tar.gz
                  bioconductor: true
                - package_name: xfun
                  package_path: xfun_0.56.tar.gz
                  bioconductor: true
                - package_name: yaml
                  package_path: yaml_2.3.12.tar.gz
                  bioconductor: true
                - package_name: base64enc
                  package_path: base64enc_0.1-6.tar.gz
                  bioconductor: true
                - package_name: cachem
                  package_path: cachem_1.1.0.tar.gz
                  bioconductor: true
                  dependencies:
                    - package_name: rlang
                      package_path: rlang_1.1.7.tar.gz
                      bioconductor: true
                    - package_name: fastmap
                      package_path: fastmap_1.2.0.tar.gz
                      bioconductor: true
                - package_name: fastmap
                  package_path: fastmap_1.2.0.tar.gz
                  bioconductor: true
                - package_name: lifecycle
                  package_path: lifecycle_1.0.5.tar.gz
                  bioconductor: true
                  dependencies:
                    - package_name: cli
                      package_path: cli_3.6.5.tar.gz
                      bioconductor: true
                - package_name: memoise
                  package_path: memoise_2.0.1.tar.gz
                  bioconductor: true
                  dependencies:
                    - package_name: cachem
                      package_path: cachem_1.1.0.tar.gz
                      bioconductor: true
                      dependencies:
                        - package_name: rlang
                          package_path: rlang_1.1.7.tar.gz
                          bioconductor: true
                        - package_name: fastmap
                          package_path: fastmap_1.2.0.tar.gz
                          bioconductor: true
                    - package_name: fastmap
                      package_path: fastmap_1.2.0.tar.gz
                      bioconductor: true
                    - package_name: rlang
                      package_path: rlang_1.1.7.tar.gz
                      bioconductor: true
                - package_name: mime
                  package_path: mime_0.13.tar.gz
                  bioconductor: true
                - package_name: rlang
                  package_path: rlang_1.1.7.tar.gz
                  bioconductor: true
                - package_name: jquerylib
                  package_path: jquerylib_0.1.4.tar.gz
                  bioconductor: true
                  dependencies:
                    - package_name: fastmap
                      package_path: fastmap_1.2.0.tar.gz
                      bioconductor: true
                    - package_name: htmltools
                      package_path: htmltools_0.5.9.tar.gz
                      bioconductor: true
                      dependencies:
                        - package_name: digest
                          package_path: digest_0.6.39.tar.gz
                          bioconductor: true
                        - package_name: rlang
                          package_path: rlang_1.1.7.tar.gz
                          bioconductor: true
                        - package_name: base64enc
                          package_path: base64enc_0.1-6.tar.gz
                          bioconductor: true
                        - package_name: fastmap
                          package_path: fastmap_1.2.0.tar.gz
                          bioconductor: true
                    - package_name: rlang
                      package_path: rlang_1.1.7.tar.gz
                      bioconductor: true
                    - package_name: base64enc
                      package_path: base64enc_0.1-6.tar.gz
                      bioconductor: true
                    - package_name: digest
                      package_path: digest_0.6.39.tar.gz
                      bioconductor: true
                - package_name: digest
                  package_path: digest_0.6.39.tar.gz
                  bioconductor: true
                - package_name: highr
                  package_path: highr_0.12.tar.gz
                  bioconductor: true
                  dependencies:
                    - package_name: xfun
                      package_path: xfun_0.56.tar.gz
                      bioconductor: true
                - package_name: cli
                  package_path: cli_3.6.5.tar.gz
                  bioconductor: true
                - package_name: R6
                  package_path: R6_2.6.1.tar.gz
                  bioconductor: true
                - package_name: rappdirs
                  package_path: rappdirs_0.3.4.tar.gz
                  bioconductor: true
                - package_name: tinytex
                  package_path: tinytex_0.58.tar.gz
                  bioconductor: true
                  dependencies:
                    - package_name: xfun
                      package_path: xfun_0.56.tar.gz
                      bioconductor: true
                - package_name: fs
                  package_path: fs_1.6.7.tar.gz
                  bioconductor: true
                  external_dependencies:
                    apt:
                        - name: make
                          dependencies:
                            - name: gcc-14-base
                            - name: libc6
                            - name: libgcc-s1
                - package_name: sass
                  package_path: sass_0.4.10.tar.gz
                  bioconductor: true
                  dependencies:
                    - package_name: R6
                      package_path: R6_2.6.1.tar.gz
                      bioconductor: true
                    - package_name: rappdirs
                      package_path: rappdirs_0.3.4.tar.gz
                      bioconductor: true
                    - package_name: rlang
                      package_path: rlang_1.1.7.tar.gz
                      bioconductor: true
                    - package_name: base64enc
                      package_path: base64enc_0.1-6.tar.gz
                      bioconductor: true
                    - package_name: digest
                      package_path: digest_0.6.39.tar.gz
                      bioconductor: true
                    - package_name: fastmap
                      package_path: fastmap_1.2.0.tar.gz
                      bioconductor: true
                    - package_name: htmltools
                      package_path: htmltools_0.5.9.tar.gz
                      bioconductor: true
                      dependencies:
                        - package_name: digest
                          package_path: digest_0.6.39.tar.gz
                          bioconductor: true
                        - package_name: rlang
                          package_path: rlang_1.1.7.tar.gz
                          bioconductor: true
                        - package_name: base64enc
                          package_path: base64enc_0.1-6.tar.gz
                          bioconductor: true
                        - package_name: fastmap
                          package_path: fastmap_1.2.0.tar.gz
                          bioconductor: true
                  external_dependencies:
                    apt:
                        - name: make
                          dependencies:
                            - name: gcc-14-base
                            - name: libc6
                            - name: libgcc-s1
              external_dependencies:
                apt:
                    - name: pandoc
                      dependencies:
                        - name: gcc-14-base
                        - name: libgcc-s1
                        - name: libc6
                        - name: libstdc++6
                        - name: zlib1g
                        - name: libyaml-0-2
                        - name: liblua5.4-0
                        - name: libgmp10
                        - name: libffi8
                        - name: pandoc-data
            - package_name: bslib
              package_path: bslib_0.10.0.tar.gz
              bioconductor: true
              dependencies:
                - package_name: cachem
                  package_path: cachem_1.1.0.tar.gz
                  bioconductor: true
                  dependencies:
                    - package_name: rlang
                      package_path: rlang_1.1.7.tar.gz
                      bioconductor: true
                    - package_name: fastmap
                      package_path: fastmap_1.2.0.tar.gz
                      bioconductor: true
                - package_name: htmltools
                  package_path: htmltools_0.5.9.tar.gz
                  bioconductor: true
                  dependencies:
                    - package_name: digest
                      package_path: digest_0.6.39.tar.gz
                      bioconductor: true
                    - package_name: rlang
                      package_path: rlang_1.1.7.tar.gz
                      bioconductor: true
                    - package_name: base64enc
                      package_path: base64enc_0.1-6.tar.gz
                      bioconductor: true
                    - package_name: fastmap
                      package_path: fastmap_1.2.0.tar.gz
                      bioconductor: true
                - package_name: jquerylib
                  package_path: jquerylib_0.1.4.tar.gz
                  bioconductor: true
                  dependencies:
                    - package_name: fastmap
                      package_path: fastmap_1.2.0.tar.gz
                      bioconductor: true
                    - package_name: base64enc
                      package_path: base64enc_0.1-6.tar.gz
                      bioconductor: true
                    - package_name: rlang
                      package_path: rlang_1.1.7.tar.gz
                      bioconductor: true
                    - package_name: htmltools
                      package_path: htmltools_0.5.9.tar.gz
                      bioconductor: true
                      dependencies:
                        - package_name: digest
                          package_path: digest_0.6.39.tar.gz
                          bioconductor: true
                        - package_name: rlang
                          package_path: rlang_1.1.7.tar.gz
                          bioconductor: true
                        - package_name: base64enc
                          package_path: base64enc_0.1-6.tar.gz
                          bioconductor: true
                        - package_name: fastmap
                          package_path: fastmap_1.2.0.tar.gz
                          bioconductor: true
                    - package_name: digest
                      package_path: digest_0.6.39.tar.gz
                      bioconductor: true
                - package_name: jsonlite
                  package_path: jsonlite_2.0.0.tar.gz
                  bioconductor: true
                - package_name: lifecycle
                  package_path: lifecycle_1.0.5.tar.gz
                  bioconductor: true
                  dependencies:
                    - package_name: cli
                      package_path: cli_3.6.5.tar.gz
                      bioconductor: true
                - package_name: memoise
                  package_path: memoise_2.0.1.tar.gz
                  bioconductor: true
                  dependencies:
                    - package_name: cachem
                      package_path: cachem_1.1.0.tar.gz
                      bioconductor: true
                      dependencies:
                        - package_name: rlang
                          package_path: rlang_1.1.7.tar.gz
                          bioconductor: true
                        - package_name: fastmap
                          package_path: fastmap_1.2.0.tar.gz
                          bioconductor: true
                    - package_name: fastmap
                      package_path: fastmap_1.2.0.tar.gz
                      bioconductor: true
                    - package_name: rlang
                      package_path: rlang_1.1.7.tar.gz
                      bioconductor: true
                - package_name: mime
                  package_path: mime_0.13.tar.gz
                  bioconductor: true
                - package_name: rlang
                  package_path: rlang_1.1.7.tar.gz
                  bioconductor: true
                - package_name: sass
                  package_path: sass_0.4.10.tar.gz
                  bioconductor: true
                  dependencies:
                    - package_name: R6
                      package_path: R6_2.6.1.tar.gz
                      bioconductor: true
                    - package_name: htmltools
                      package_path: htmltools_0.5.9.tar.gz
                      bioconductor: true
                      dependencies:
                        - package_name: digest
                          package_path: digest_0.6.39.tar.gz
                          bioconductor: true
                        - package_name: rlang
                          package_path: rlang_1.1.7.tar.gz
                          bioconductor: true
                        - package_name: base64enc
                          package_path: base64enc_0.1-6.tar.gz
                          bioconductor: true
                        - package_name: fastmap
                          package_path: fastmap_1.2.0.tar.gz
                          bioconductor: true
                    - package_name: rlang
                      package_path: rlang_1.1.7.tar.gz
                      bioconductor: true
                    - package_name: base64enc
                      package_path: base64enc_0.1-6.tar.gz
                      bioconductor: true
                    - package_name: digest
                      package_path: digest_0.6.39.tar.gz
                      bioconductor: true
                    - package_name: fastmap
                      package_path: fastmap_1.2.0.tar.gz
                      bioconductor: true
                    - package_name: rappdirs
                      package_path: rappdirs_0.3.4.tar.gz
                      bioconductor: true
                  external_dependencies:
                    apt:
                        - name: make
                          dependencies:
                            - name: gcc-14-base
                            - name: libc6
                            - name: libgcc-s1
                - package_name: digest
                  package_path: digest_0.6.39.tar.gz
                  bioconductor: true
                - package_name: base64enc
                  package_path: base64enc_0.1-6.tar.gz
                  bioconductor: true
                - package_name: fastmap
                  package_path: fastmap_1.2.0.tar.gz
                  bioconductor: true
                - package_name: cli
                  package_path: cli_3.6.5.tar.gz
                  bioconductor: true
                - package_name: R6
                  package_path: R6_2.6.1.tar.gz
                  bioconductor: true
                - package_name: rappdirs
                  package_path: rappdirs_0.3.4.tar.gz
                  bioconductor: true
                - package_name: fs
                  package_path: fs_1.6.7.tar.gz
                  bioconductor: true
                  external_dependencies:
                    apt:
                        - name: make
                          dependencies:
                            - name: gcc-14-base
                            - name: libc6
                            - name: libgcc-s1
        - package_name: tidyr
          package_path: tidyr_1.3.2.tar.gz
          bioconductor: true
          dependencies:
            - package_name: lifecycle
              package_path: lifecycle_1.0.5.tar.gz
              bioconductor: true
              dependencies:
                - package_name: cli
                  package_path: cli_3.6.5.tar.gz
                  bioconductor: true
            - package_name: magrittr
              package_path: magrittr_2.0.4.tar.gz
              bioconductor: true
            - package_name: glue
              package_path: glue_1.8.0.tar.gz
              bioconductor: true
            - package_name: rlang
              package_path: rlang_1.1.7.tar.gz
              bioconductor: true
            - package_name: stringr
              package_path: stringr_1.6.0.tar.gz
              bioconductor: true
              dependencies:
                - package_name: magrittr
                  package_path: magrittr_2.0.4.tar.gz
                  bioconductor: true
                - package_name: glue
                  package_path: glue_1.8.0.tar.gz
                  bioconductor: true
                - package_name: lifecycle
                  package_path: lifecycle_1.0.5.tar.gz
                  bioconductor: true
                  dependencies:
                    - package_name: cli
                      package_path: cli_3.6.5.tar.gz
                      bioconductor: true
                - package_name: stringi
                  package_path: stringi_1.8.7.tar.gz
                  bioconductor: true
                  external_dependencies:
                    apt:
                        - name: libicu-dev
                          dependencies:
                            - name: libc-dev
                              optional: true
                            - name: libc6
                            - name: libcrypt1
                            - name: gcc-14-base
                            - name: rpcsvc-proto
                            - name: libcrypt-dev
                            - name: linux-libc-dev
                            - name: libc-dev-bin
                            - name: libgcc-s1
                            - name: libstdc++6
                            - name: libc6-dev
                            - name: libicu74
                            - name: icu-devtools
                - package_name: rlang
                  package_path: rlang_1.1.7.tar.gz
                  bioconductor: true
            - package_name: tibble
              package_path: tibble_3.3.1.tar.gz
              bioconductor: true
              dependencies:
                - package_name: magrittr
                  package_path: magrittr_2.0.4.tar.gz
                  bioconductor: true
                - package_name: rlang
                  package_path: rlang_1.1.7.tar.gz
                  bioconductor: true
                - package_name: pkgconfig
                  package_path: pkgconfig_2.0.3.tar.gz
                  bioconductor: true
                - package_name: lifecycle
                  package_path: lifecycle_1.0.5.tar.gz
                  bioconductor: true
                - package_name: glue
                  package_path: glue_1.8.0.tar.gz
                  bioconductor: true
                - package_name: pillar
                  package_path: pillar_1.11.1.tar.gz
                  bioconductor: true
                  dependencies:
                    - package_name: rlang
                      package_path: rlang_1.1.7.tar.gz
                      bioconductor: true
                    - package_name: glue
                      package_path: glue_1.8.0.tar.gz
                      bioconductor: true
                    - package_name: utf8
                      package_path: utf8_1.2.6.tar.gz
                      bioconductor: true
                    - package_name: lifecycle
                      package_path: lifecycle_1.0.5.tar.gz
                      bioconductor: true
                      dependencies:
                        - package_name: rlang
                          package_path: rlang_1.1.7.tar.gz
                          bioconductor: true
                        - package_name: cli
                          package_path: cli_3.6.5.tar.gz
                          bioconductor: true
                - package_name: utf8
                  package_path: utf8_1.2.6.tar.gz
                  bioconductor: true
                - package_name: vctrs
                  package_path: vctrs_0.7.1.tar.gz
                  bioconductor: true
                  dependencies:
                    - package_name: rlang
                      package_path: rlang_1.1.7.tar.gz
                      bioconductor: true
                    - package_name: glue
                      package_path: glue_1.8.0.tar.gz
                      bioconductor: true
                    - package_name: cli
                      package_path: cli_3.6.5.tar.gz
                      bioconductor: true
                    - package_name: lifecycle
                      package_path: lifecycle_1.0.5.tar.gz
                      bioconductor: true
                      dependencies:
                        - package_name: cli
                          package_path: cli_3.6.5.tar.gz
                          bioconductor: true
            - package_name: cli
              package_path: cli_3.6.5.tar.gz
              bioconductor: true
            - package_name: vctrs
              package_path: vctrs_0.7.1.tar.gz
              bioconductor: true
              dependencies:
                - package_name: rlang
                  package_path: rlang_1.1.7.tar.gz
                  bioconductor: true
                - package_name: glue
                  package_path: glue_1.8.0.tar.gz
                  bioconductor: true
                - package_name: cli
                  package_path: cli_3.6.5.tar.gz
                  bioconductor: true
                - package_name: lifecycle
                  package_path: lifecycle_1.0.5.tar.gz
                  bioconductor: true
                  dependencies:
                    - package_name: cli
                      package_path: cli_3.6.5.tar.gz
                      bioconductor: true
            - package_name: cpp11
              package_path: cpp11_0.5.3.tar.gz
              bioconductor: true
            - package_name: generics
              package_path: generics_0.1.4.tar.gz
              bioconductor: true
            - package_name: purrr
              package_path: purrr_1.2.1.tar.gz
              bioconductor: true
              dependencies:
                - package_name: rlang
                  package_path: rlang_1.1.7.tar.gz
                  bioconductor: true
                - package_name: vctrs
                  package_path: vctrs_0.7.1.tar.gz
                  bioconductor: true
                  dependencies:
                    - package_name: rlang
                      package_path: rlang_1.1.7.tar.gz
                      bioconductor: true
                    - package_name: glue
                      package_path: glue_1.8.0.tar.gz
                      bioconductor: true
                    - package_name: cli
                      package_path: cli_3.6.5.tar.gz
                      bioconductor: true
                    - package_name: lifecycle
                      package_path: lifecycle_1.0.5.tar.gz
                      bioconductor: true
                      dependencies:
                        - package_name: cli
                          package_path: cli_3.6.5.tar.gz
                          bioconductor: true
                - package_name: lifecycle
                  package_path: lifecycle_1.0.5.tar.gz
                  bioconductor: true
                  dependencies:
                    - package_name: cli
                      package_path: cli_3.6.5.tar.gz
                      bioconductor: true
                - package_name: glue
                  package_path: glue_1.8.0.tar.gz
                  bioconductor: true
                - package_name: cli
                  package_path: cli_3.6.5.tar.gz
                  bioconductor: true
                - package_name: magrittr
                  package_path: magrittr_2.0.4.tar.gz
                  bioconductor: true
            - package_name: pillar
              package_path: pillar_1.11.1.tar.gz
              bioconductor: true
              dependencies:
                - package_name: rlang
                  package_path: rlang_1.1.7.tar.gz
                  bioconductor: true
                - package_name: glue
                  package_path: glue_1.8.0.tar.gz
                  bioconductor: true
                - package_name: utf8
                  package_path: utf8_1.2.6.tar.gz
                  bioconductor: true
                - package_name: lifecycle
                  package_path: lifecycle_1.0.5.tar.gz
                  bioconductor: true
                  dependencies:
                    - package_name: rlang
                      package_path: rlang_1.1.7.tar.gz
                      bioconductor: true
                    - package_name: cli
                      package_path: cli_3.6.5.tar.gz
                      bioconductor: true
            - package_name: R6
              package_path: R6_2.6.1.tar.gz
              bioconductor: true
            - package_name: stringi
              package_path: stringi_1.8.7.tar.gz
              bioconductor: true
              external_dependencies:
                apt:
                    - name: libicu-dev
                      dependencies:
                        - name: libc-dev
                          optional: true
                        - name: libc6
                        - name: libcrypt1
                        - name: gcc-14-base
                        - name: rpcsvc-proto
                        - name: libcrypt-dev
                        - name: linux-libc-dev
                        - name: libc-dev-bin
                        - name: libgcc-s1
                        - name: libstdc++6
                        - name: libc6-dev
                        - name: libicu74
                        - name: icu-devtools
            - package_name: pkgconfig
              package_path: pkgconfig_2.0.3.tar.gz
              bioconductor: true
            - package_name: withr
              package_path: withr_3.0.2.tar.gz
              bioconductor: true
            - package_name: utf8
              package_path: utf8_1.2.6.tar.gz
              bioconductor: true
            - package_name: tidyselect
              package_path: tidyselect_1.2.1.tar.gz
              bioconductor: true
              dependencies:
                - package_name: rlang
                  package_path: rlang_1.1.7.tar.gz
                  bioconductor: true
                - package_name: vctrs
                  package_path: vctrs_0.7.1.tar.gz
                  bioconductor: true
                  dependencies:
                    - package_name: rlang
                      package_path: rlang_1.1.7.tar.gz
                      bioconductor: true
                    - package_name: glue
                      package_path: glue_1.8.0.tar.gz
                      bioconductor: true
                    - package_name: cli
                      package_path: cli_3.6.5.tar.gz
                      bioconductor: true
                    - package_name: lifecycle
                      package_path: lifecycle_1.0.5.tar.gz
                      bioconductor: true
                      dependencies:
                        - package_name: cli
                          package_path: cli_3.6.5.tar.gz
                          bioconductor: true
                - package_name: withr
                  package_path: withr_3.0.2.tar.gz
                  bioconductor: true
                - package_name: glue
                  package_path: glue_1.8.0.tar.gz
                  bioconductor: true
                - package_name: lifecycle
                  package_path: lifecycle_1.0.5.tar.gz
                  bioconductor: true
                  dependencies:
                    - package_name: cli
                      package_path: cli_3.6.5.tar.gz
                      bioconductor: true
                - package_name: cli
                  package_path: cli_3.6.5.tar.gz
                  bioconductor: true
            - package_name: dplyr
              package_path: dplyr_1.2.0.tar.gz
              bioconductor: true
              dependencies:
                - package_name: lifecycle
                  package_path: lifecycle_1.0.5.tar.gz
                  bioconductor: true
                  dependencies:
                    - package_name: cli
                      package_path: cli_3.6.5.tar.gz
                      bioconductor: true
                - package_name: magrittr
                  package_path: magrittr_2.0.4.tar.gz
                  bioconductor: true
                - package_name: cli
                  package_path: cli_3.6.5.tar.gz
                  bioconductor: true
                - package_name: pillar
                  package_path: pillar_1.11.1.tar.gz
                  bioconductor: true
                  dependencies:
                    - package_name: rlang
                      package_path: rlang_1.1.7.tar.gz
                      bioconductor: true
                    - package_name: glue
                      package_path: glue_1.8.0.tar.gz
                      bioconductor: true
                    - package_name: utf8
                      package_path: utf8_1.2.6.tar.gz
                      bioconductor: true
                    - package_name: lifecycle
                      package_path: lifecycle_1.0.5.tar.gz
                      bioconductor: true
                      dependencies:
                        - package_name: rlang
                          package_path: rlang_1.1.7.tar.gz
                          bioconductor: true
                        - package_name: cli
                          package_path: cli_3.6.5.tar.gz
                          bioconductor: true
                - package_name: glue
                  package_path: glue_1.8.0.tar.gz
                  bioconductor: true
                - package_name: rlang
                  package_path: rlang_1.1.7.tar.gz
                  bioconductor: true
                - package_name: tibble
                  package_path: tibble_3.3.1.tar.gz
                  bioconductor: true
                  dependencies:
                    - package_name: magrittr
                      package_path: magrittr_2.0.4.tar.gz
                      bioconductor: true
                    - package_name: rlang
                      package_path: rlang_1.1.7.tar.gz
                      bioconductor: true
                    - package_name: pkgconfig
                      package_path: pkgconfig_2.0.3.tar.gz
                      bioconductor: true
                    - package_name: lifecycle
                      package_path: lifecycle_1.0.5.tar.gz
                      bioconductor: true
                    - package_name: glue
                      package_path: glue_1.8.0.tar.gz
                      bioconductor: true
                    - package_name: pillar
                      package_path: pillar_1.11.1.tar.gz
                      bioconductor: true
                      dependencies:
                        - package_name: rlang
                          package_path: rlang_1.1.7.tar.gz
                          bioconductor: true
                        - package_name: glue
                          package_path: glue_1.8.0.tar.gz
                          bioconductor: true
                        - package_name: utf8
                          package_path: utf8_1.2.6.tar.gz
                          bioconductor: true
                        - package_name: lifecycle
                          package_path: lifecycle_1.0.5.tar.gz
                          bioconductor: true
                          dependencies:
                            - package_name: rlang
                              package_path: rlang_1.1.7.tar.gz
                              bioconductor: true
                            - package_name: cli
                              package_path: cli_3.6.5.tar.gz
                              bioconductor: true
                    - package_name: utf8
                      package_path: utf8_1.2.6.tar.gz
                      bioconductor: true
                    - package_name: vctrs
                      package_path: vctrs_0.7.1.tar.gz
                      bioconductor: true
                      dependencies:
                        - package_name: rlang
                          package_path: rlang_1.1.7.tar.gz
                          bioconductor: true
                        - package_name: glue
                          package_path: glue_1.8.0.tar.gz
                          bioconductor: true
                        - package_name: cli
                          package_path: cli_3.6.5.tar.gz
                          bioconductor: true
                        - package_name: lifecycle
                          package_path: lifecycle_1.0.5.tar.gz
                          bioconductor: true
                          dependencies:
                            - package_name: cli
                              package_path: cli_3.6.5.tar.gz
                              bioconductor: true
                - package_name: R6
                  package_path: R6_2.6.1.tar.gz
                  bioconductor: true
                - package_name: generics
                  package_path: generics_0.1.4.tar.gz
                  bioconductor: true
                - package_name: vctrs
                  package_path: vctrs_0.7.1.tar.gz
                  bioconductor: true
                  dependencies:
                    - package_name: rlang
                      package_path: rlang_1.1.7.tar.gz
                      bioconductor: true
                    - package_name: glue
                      package_path: glue_1.8.0.tar.gz
                      bioconductor: true
                    - package_name: cli
                      package_path: cli_3.6.5.tar.gz
                      bioconductor: true
                    - package_name: lifecycle
                      package_path: lifecycle_1.0.5.tar.gz
                      bioconductor: true
                      dependencies:
                        - package_name: cli
                          package_path: cli_3.6.5.tar.gz
                          bioconductor: true
                - package_name: utf8
                  package_path: utf8_1.2.6.tar.gz
                  bioconductor: true
                - package_name: pkgconfig
                  package_path: pkgconfig_2.0.3.tar.gz
                  bioconductor: true
                - package_name: withr
                  package_path: withr_3.0.2.tar.gz
                  bioconductor: true
                - package_name: tidyselect
                  package_path: tidyselect_1.2.1.tar.gz
                  bioconductor: true
                  dependencies:
                    - package_name: rlang
                      package_path: rlang_1.1.7.tar.gz
                      bioconductor: true
                    - package_name: vctrs
                      package_path: vctrs_0.7.1.tar.gz
                      bioconductor: true
                      dependencies:
                        - package_name: rlang
                          package_path: rlang_1.1.7.tar.gz
                          bioconductor: true
                        - package_name: glue
                          package_path: glue_1.8.0.tar.gz
                          bioconductor: true
                        - package_name: cli
                          package_path: cli_3.6.5.tar.gz
                          bioconductor: true
                        - package_name: lifecycle
                          package_path: lifecycle_1.0.5.tar.gz
                          bioconductor: true
                          dependencies:
                            - package_name: cli
                              package_path: cli_3.6.5.tar.gz
                              bioconductor: true
                    - package_name: withr
                      package_path: withr_3.0.2.tar.gz
                      bioconductor: true
                    - package_name: cli
                      package_path: cli_3.6.5.tar.gz
                      bioconductor: true
                    - package_name: glue
                      package_path: glue_1.8.0.tar.gz
                      bioconductor: true
                    - package_name: lifecycle
                      package_path: lifecycle_1.0.5.tar.gz
                      bioconductor: true
                      dependencies:
                        - package_name: cli
                          package_path: cli_3.6.5.tar.gz
                          bioconductor: true
        - package_name: dplyr
          package_path: dplyr_1.2.0.tar.gz
          bioconductor: true
          dependencies:
            - package_name: lifecycle
              package_path: lifecycle_1.0.5.tar.gz
              bioconductor: true
              dependencies:
                - package_name: cli
                  package_path: cli_3.6.5.tar.gz
                  bioconductor: true
            - package_name: generics
              package_path: generics_0.1.4.tar.gz
              bioconductor: true
            - package_name: cli
              package_path: cli_3.6.5.tar.gz
              bioconductor: true
            - package_name: pillar
              package_path: pillar_1.11.1.tar.gz
              bioconductor: true
              dependencies:
                - package_name: rlang
                  package_path: rlang_1.1.7.tar.gz
                  bioconductor: true
                - package_name: glue
                  package_path: glue_1.8.0.tar.gz
                  bioconductor: true
                - package_name: utf8
                  package_path: utf8_1.2.6.tar.gz
                  bioconductor: true
                - package_name: lifecycle
                  package_path: lifecycle_1.0.5.tar.gz
                  bioconductor: true
                  dependencies:
                    - package_name: rlang
                      package_path: rlang_1.1.7.tar.gz
                      bioconductor: true
                    - package_name: cli
                      package_path: cli_3.6.5.tar.gz
                      bioconductor: true
            - package_name: R6
              package_path: R6_2.6.1.tar.gz
              bioconductor: true
            - package_name: rlang
              package_path: rlang_1.1.7.tar.gz
              bioconductor: true
            - package_name: tibble
              package_path: tibble_3.3.1.tar.gz
              bioconductor: true
              dependencies:
                - package_name: magrittr
                  package_path: magrittr_2.0.4.tar.gz
                  bioconductor: true
                - package_name: rlang
                  package_path: rlang_1.1.7.tar.gz
                  bioconductor: true
                - package_name: pkgconfig
                  package_path: pkgconfig_2.0.3.tar.gz
                  bioconductor: true
                - package_name: lifecycle
                  package_path: lifecycle_1.0.5.tar.gz
                  bioconductor: true
                - package_name: glue
                  package_path: glue_1.8.0.tar.gz
                  bioconductor: true
                - package_name: pillar
                  package_path: pillar_1.11.1.tar.gz
                  bioconductor: true
                  dependencies:
                    - package_name: rlang
                      package_path: rlang_1.1.7.tar.gz
                      bioconductor: true
                    - package_name: glue
                      package_path: glue_1.8.0.tar.gz
                      bioconductor: true
                    - package_name: utf8
                      package_path: utf8_1.2.6.tar.gz
                      bioconductor: true
                    - package_name: lifecycle
                      package_path: lifecycle_1.0.5.tar.gz
                      bioconductor: true
                      dependencies:
                        - package_name: rlang
                          package_path: rlang_1.1.7.tar.gz
                          bioconductor: true
                        - package_name: cli
                          package_path: cli_3.6.5.tar.gz
                          bioconductor: true
                - package_name: utf8
                  package_path: utf8_1.2.6.tar.gz
                  bioconductor: true
                - package_name: vctrs
                  package_path: vctrs_0.7.1.tar.gz
                  bioconductor: true
                  dependencies:
                    - package_name: rlang
                      package_path: rlang_1.1.7.tar.gz
                      bioconductor: true
                    - package_name: glue
                      package_path: glue_1.8.0.tar.gz
                      bioconductor: true
                    - package_name: cli
                      package_path: cli_3.6.5.tar.gz
                      bioconductor: true
                    - package_name: lifecycle
                      package_path: lifecycle_1.0.5.tar.gz
                      bioconductor: true
                      dependencies:
                        - package_name: cli
                          package_path: cli_3.6.5.tar.gz
                          bioconductor: true
            - package_name: glue
              package_path: glue_1.8.0.tar.gz
              bioconductor: true
            - package_name: magrittr
              package_path: magrittr_2.0.4.tar.gz
              bioconductor: true
            - package_name: vctrs
              package_path: vctrs_0.7.1.tar.gz
              bioconductor: true
              dependencies:
                - package_name: rlang
                  package_path: rlang_1.1.7.tar.gz
                  bioconductor: true
                - package_name: glue
                  package_path: glue_1.8.0.tar.gz
                  bioconductor: true
                - package_name: cli
                  package_path: cli_3.6.5.tar.gz
                  bioconductor: true
                - package_name: lifecycle
                  package_path: lifecycle_1.0.5.tar.gz
                  bioconductor: true
                  dependencies:
                    - package_name: cli
                      package_path: cli_3.6.5.tar.gz
                      bioconductor: true
            - package_name: utf8
              package_path: utf8_1.2.6.tar.gz
              bioconductor: true
            - package_name: pkgconfig
              package_path: pkgconfig_2.0.3.tar.gz
              bioconductor: true
            - package_name: withr
              package_path: withr_3.0.2.tar.gz
              bioconductor: true
        - package_name: lazyeval
          package_path: lazyeval_0.2.2.tar.gz
          bioconductor: true
        - package_name: crosstalk
          package_path: crosstalk_1.2.2.tar.gz
          bioconductor: true
          dependencies:
            - package_name: R6
              package_path: R6_2.6.1.tar.gz
              bioconductor: true
            - package_name: base64enc
              package_path: base64enc_0.1-6.tar.gz
              bioconductor: true
            - package_name: digest
              package_path: digest_0.6.39.tar.gz
              bioconductor: true
            - package_name: fastmap
              package_path: fastmap_1.2.0.tar.gz
              bioconductor: true
            - package_name: htmltools
              package_path: htmltools_0.5.9.tar.gz
              bioconductor: true
              dependencies:
                - package_name: digest
                  package_path: digest_0.6.39.tar.gz
                  bioconductor: true
                - package_name: rlang
                  package_path: rlang_1.1.7.tar.gz
                  bioconductor: true
                - package_name: base64enc
                  package_path: base64enc_0.1-6.tar.gz
                  bioconductor: true
                - package_name: fastmap
                  package_path: fastmap_1.2.0.tar.gz
                  bioconductor: true
            - package_name: rlang
              package_path: rlang_1.1.7.tar.gz
              bioconductor: true
            - package_name: lazyeval
              package_path: lazyeval_0.2.2.tar.gz
              bioconductor: true
            - package_name: jsonlite
              package_path: jsonlite_2.0.0.tar.gz
              bioconductor: true
        - package_name: purrr
          package_path: purrr_1.2.1.tar.gz
          bioconductor: true
          dependencies:
            - package_name: rlang
              package_path: rlang_1.1.7.tar.gz
              bioconductor: true
            - package_name: vctrs
              package_path: vctrs_0.7.1.tar.gz
              bioconductor: true
              dependencies:
                - package_name: rlang
                  package_path: rlang_1.1.7.tar.gz
                  bioconductor: true
                - package_name: glue
                  package_path: glue_1.8.0.tar.gz
                  bioconductor: true
                - package_name: cli
                  package_path: cli_3.6.5.tar.gz
                  bioconductor: true
                - package_name: lifecycle
                  package_path: lifecycle_1.0.5.tar.gz
                  bioconductor: true
                  dependencies:
                    - package_name: cli
                      package_path: cli_3.6.5.tar.gz
                      bioconductor: true
            - package_name: lifecycle
              package_path: lifecycle_1.0.5.tar.gz
              bioconductor: true
              dependencies:
                - package_name: cli
                  package_path: cli_3.6.5.tar.gz
                  bioconductor: true
            - package_name: glue
              package_path: glue_1.8.0.tar.gz
              bioconductor: true
            - package_name: cli
              package_path: cli_3.6.5.tar.gz
              bioconductor: true
            - package_name: magrittr
              package_path: magrittr_2.0.4.tar.gz
              bioconductor: true
        - package_name: promises
          package_path: promises_1.5.0.tar.gz
          bioconductor: true
          dependencies:
            - package_name: magrittr
              package_path: magrittr_2.0.4.tar.gz
              bioconductor: true
            - package_name: later
              package_path: later_1.4.8.tar.gz
              bioconductor: true
              dependencies:
                - package_name: rlang
                  package_path: rlang_1.1.7.tar.gz
                  bioconductor: true
                - package_name: Rcpp
                  package_path: Rcpp_1.1.1.tar.gz
                  bioconductor: true
            - package_name: R6
              package_path: R6_2.6.1.tar.gz
              bioconductor: true
            - package_name: rlang
              package_path: rlang_1.1.7.tar.gz
              bioconductor: true
            - package_name: Rcpp
              package_path: Rcpp_1.1.1.tar.gz
              bioconductor: true
            - package_name: fastmap
              package_path: fastmap_1.2.0.tar.gz
              bioconductor: true
            - package_name: lifecycle
              package_path: lifecycle_1.0.5.tar.gz
              bioconductor: true
              dependencies:
                - package_name: cli
                  package_path: cli_3.6.5.tar.gz
                  bioconductor: true
            - package_name: cli
              package_path: cli_3.6.5.tar.gz
              bioconductor: true
            - package_name: otel
              package_path: otel_0.2.0.tar.gz
              bioconductor: true
        - package_name: RcppTOML
          package_path: RcppTOML_0.2.3.tar.gz
          bioconductor: true
          dependencies:
            - package_name: Rcpp
              package_path: Rcpp_1.1.1.tar.gz
              bioconductor: true
        - package_name: here
          package_path: here_1.0.2.tar.gz
          bioconductor: true
          dependencies:
            - package_name: rprojroot
              package_path: rprojroot_2.1.1.tar.gz
              bioconductor: true
        - package_name: rappdirs
          package_path: rappdirs_0.3.4.tar.gz
          bioconductor: true
        - package_name: gplots
          package_path: gplots_3.3.0.tar.gz
          bioconductor: true
          dependencies:
            - package_name: KernSmooth
              package_path: KernSmooth_2.23-26.tar.gz
              bioconductor: true
            - package_name: caTools
              package_path: caTools_1.18.3.tar.gz
              bioconductor: true
              dependencies:
                - package_name: bitops
                  package_path: bitops_1.0-9.tar.gz
                  bioconductor: true
            - package_name: bitops
              package_path: bitops_1.0-9.tar.gz
              bioconductor: true
            - package_name: gtools
              package_path: gtools_3.9.5.tar.gz
              bioconductor: true
        - package_name: glue
          package_path: glue_1.8.0.tar.gz
          bioconductor: true
        - package_name: labeling
          package_path: labeling_0.4.3.tar.gz
          bioconductor: true
        - package_name: reshape2
          package_path: reshape2_1.4.5.tar.gz
          bioconductor: true
          dependencies:
            - package_name: Rcpp
              package_path: Rcpp_1.1.1.tar.gz
              bioconductor: true
            - package_name: stringr
              package_path: stringr_1.6.0.tar.gz
              bioconductor: true
              dependencies:
                - package_name: magrittr
                  package_path: magrittr_2.0.4.tar.gz
                  bioconductor: true
                - package_name: glue
                  package_path: glue_1.8.0.tar.gz
                  bioconductor: true
                - package_name: lifecycle
                  package_path: lifecycle_1.0.5.tar.gz
                  bioconductor: true
                  dependencies:
                    - package_name: cli
                      package_path: cli_3.6.5.tar.gz
                      bioconductor: true
                - package_name: stringi
                  package_path: stringi_1.8.7.tar.gz
                  bioconductor: true
                  external_dependencies:
                    apt:
                        - name: libicu-dev
                          dependencies:
                            - name: libc-dev
                              optional: true
                            - name: libc6
                            - name: libcrypt1
                            - name: gcc-14-base
                            - name: rpcsvc-proto
                            - name: libcrypt-dev
                            - name: linux-libc-dev
                            - name: libc-dev-bin
                            - name: libgcc-s1
                            - name: libstdc++6
                            - name: libc6-dev
                            - name: libicu74
                            - name: icu-devtools
                - package_name: rlang
                  package_path: rlang_1.1.7.tar.gz
                  bioconductor: true
            - package_name: cli
              package_path: cli_3.6.5.tar.gz
              bioconductor: true
            - package_name: glue
              package_path: glue_1.8.0.tar.gz
              bioconductor: true
            - package_name: lifecycle
              package_path: lifecycle_1.0.5.tar.gz
              bioconductor: true
              dependencies:
                - package_name: cli
                  package_path: cli_3.6.5.tar.gz
                  bioconductor: true
            - package_name: magrittr
              package_path: magrittr_2.0.4.tar.gz
              bioconductor: true
            - package_name: rlang
              package_path: rlang_1.1.7.tar.gz
              bioconductor: true
            - package_name: stringi
              package_path: stringi_1.8.7.tar.gz
              bioconductor: true
              external_dependencies:
                apt:
                    - name: libicu-dev
                      dependencies:
                        - name: libc-dev
                          optional: true
                        - name: libc6
                        - name: libcrypt1
                        - name: gcc-14-base
                        - name: rpcsvc-proto
                        - name: libcrypt-dev
                        - name: linux-libc-dev
                        - name: libc-dev-bin
                        - name: libgcc-s1
                        - name: libstdc++6
                        - name: libc6-dev
                        - name: libicu74
                        - name: icu-devtools
            - package_name: vctrs
              package_path: vctrs_0.7.1.tar.gz
              bioconductor: true
              dependencies:
                - package_name: rlang
                  package_path: rlang_1.1.7.tar.gz
                  bioconductor: true
                - package_name: glue
                  package_path: glue_1.8.0.tar.gz
                  bioconductor: true
                - package_name: cli
                  package_path: cli_3.6.5.tar.gz
                  bioconductor: true
                - package_name: lifecycle
                  package_path: lifecycle_1.0.5.tar.gz
                  bioconductor: true
                  dependencies:
                    - package_name: cli
                      package_path: cli_3.6.5.tar.gz
                      bioconductor: true
            - package_name: plyr
              package_path: plyr_1.8.9.tar.gz
              bioconductor: true
              dependencies:
                - package_name: Rcpp
                  package_path: Rcpp_1.1.1.tar.gz
                  bioconductor: true
        - package_name: gridExtra
          package_path: gridExtra_2.3.tar.gz
          bioconductor: true
          dependencies:
            - package_name: gtable
              package_path: gtable_0.3.6.tar.gz
              bioconductor: true
              dependencies:
                - package_name: rlang
                  package_path: rlang_1.1.7.tar.gz
                  bioconductor: true
            - package_name: cli
              package_path: cli_3.6.5.tar.gz
              bioconductor: true
            - package_name: glue
              package_path: glue_1.8.0.tar.gz
              bioconductor: true
            - package_name: lifecycle
              package_path: lifecycle_1.0.5.tar.gz
              bioconductor: true
              dependencies:
                - package_name: cli
                  package_path: cli_3.6.5.tar.gz
                  bioconductor: true
            - package_name: rlang
              package_path: rlang_1.1.7.tar.gz
              bioconductor: true
        - package_name: RcppArmadillo
          package_path: RcppArmadillo_15.2.3-1.tar.gz
          bioconductor: true
          dependencies:
            - package_name: Rcpp
              package_path: Rcpp_1.1.1.tar.gz
              bioconductor: true
          external_dependencies:
            apt:
                - name: libblas-dev
                  dependencies:
                    - name: gcc-14-base
                    - name: libc6
                    - name: libgcc-s1
                    - name: libblas3
                - name: liblapack-dev
                  dependencies:
                    - name: libblas.so.3
                      optional: true
                    - name: libblas.so
                      optional: true
                    - name: gcc-14-base
                    - name: libc6
                    - name: libgomp1
                    - name: libgcc-s1
                    - name: libgfortran5
                    - name: libopenblas0-serial
                    - name: libopenblas0-pthread
                    - name: libopenblas0-openmp
                    - name: libblis4-serial
                    - name: libblis4-pthread
                    - name: libblis4-openmp
                    - name: libatlas3-base
                    - name: libblas3
                    - name: libopenblas-serial-dev
                    - name: libopenblas-pthread-dev
                    - name: libopenblas-openmp-dev
                    - name: libblis-serial-dev
                    - name: libblis-pthread-dev
                    - name: libblis-openmp-dev
                    - name: libatlas-base-dev
                    - name: libblas-dev
                    - name: liblapack3
                - name: r-base-dev
        - package_name: sp
          package_path: sp_2.2-1.tar.gz
          bioconductor: true
          dependencies:
            - package_name: lattice
              package_path: lattice_0.22-9.tar.gz
              bioconductor: true
        - package_name: spam
          package_path: spam_2.11-3.tar.gz
          bioconductor: true
          dependencies:
            - package_name: Rcpp
              package_path: Rcpp_1.1.1.tar.gz
              bioconductor: true
            - package_name: dotCall64
              package_path: dotCall64_1.2.tar.gz
              bioconductor: true
        - package_name: bslib
          package_path: bslib_0.10.0.tar.gz
          bioconductor: true
          dependencies:
            - package_name: fastmap
              package_path: fastmap_1.2.0.tar.gz
              bioconductor: true
            - package_name: htmltools
              package_path: htmltools_0.5.9.tar.gz
              bioconductor: true
              dependencies:
                - package_name: digest
                  package_path: digest_0.6.39.tar.gz
                  bioconductor: true
                - package_name: rlang
                  package_path: rlang_1.1.7.tar.gz
                  bioconductor: true
                - package_name: base64enc
                  package_path: base64enc_0.1-6.tar.gz
                  bioconductor: true
                - package_name: fastmap
                  package_path: fastmap_1.2.0.tar.gz
                  bioconductor: true
            - package_name: base64enc
              package_path: base64enc_0.1-6.tar.gz
              bioconductor: true
            - package_name: jsonlite
              package_path: jsonlite_2.0.0.tar.gz
              bioconductor: true
            - package_name: lifecycle
              package_path: lifecycle_1.0.5.tar.gz
              bioconductor: true
              dependencies:
                - package_name: cli
                  package_path: cli_3.6.5.tar.gz
                  bioconductor: true
            - package_name: cachem
              package_path: cachem_1.1.0.tar.gz
              bioconductor: true
              dependencies:
                - package_name: rlang
                  package_path: rlang_1.1.7.tar.gz
                  bioconductor: true
                - package_name: fastmap
                  package_path: fastmap_1.2.0.tar.gz
                  bioconductor: true
            - package_name: mime
              package_path: mime_0.13.tar.gz
              bioconductor: true
            - package_name: rlang
              package_path: rlang_1.1.7.tar.gz
              bioconductor: true
            - package_name: digest
              package_path: digest_0.6.39.tar.gz
              bioconductor: true
            - package_name: cli
              package_path: cli_3.6.5.tar.gz
              bioconductor: true
            - package_name: memoise
              package_path: memoise_2.0.1.tar.gz
              bioconductor: true
              dependencies:
                - package_name: cachem
                  package_path: cachem_1.1.0.tar.gz
                  bioconductor: true
                  dependencies:
                    - package_name: rlang
                      package_path: rlang_1.1.7.tar.gz
                      bioconductor: true
                    - package_name: fastmap
                      package_path: fastmap_1.2.0.tar.gz
                      bioconductor: true
                - package_name: rlang
                  package_path: rlang_1.1.7.tar.gz
                  bioconductor: true
                - package_name: fastmap
                  package_path: fastmap_1.2.0.tar.gz
                  bioconductor: true
            - package_name: R6
              package_path: R6_2.6.1.tar.gz
              bioconductor: true
            - package_name: jquerylib
              package_path: jquerylib_0.1.4.tar.gz
              bioconductor: true
              dependencies:
                - package_name: base64enc
                  package_path: base64enc_0.1-6.tar.gz
                  bioconductor: true
                - package_name: fastmap
                  package_path: fastmap_1.2.0.tar.gz
                  bioconductor: true
                - package_name: rlang
                  package_path: rlang_1.1.7.tar.gz
                  bioconductor: true
                - package_name: htmltools
                  package_path: htmltools_0.5.9.tar.gz
                  bioconductor: true
                  dependencies:
                    - package_name: digest
                      package_path: digest_0.6.39.tar.gz
                      bioconductor: true
                    - package_name: rlang
                      package_path: rlang_1.1.7.tar.gz
                      bioconductor: true
                    - package_name: base64enc
                      package_path: base64enc_0.1-6.tar.gz
                      bioconductor: true
                    - package_name: fastmap
                      package_path: fastmap_1.2.0.tar.gz
                      bioconductor: true
                - package_name: digest
                  package_path: digest_0.6.39.tar.gz
                  bioconductor: true
            - package_name: sass
              package_path: sass_0.4.10.tar.gz
              bioconductor: true
              dependencies:
                - package_name: R6
                  package_path: R6_2.6.1.tar.gz
                  bioconductor: true
                - package_name: htmltools
                  package_path: htmltools_0.5.9.tar.gz
                  bioconductor: true
                  dependencies:
                    - package_name: digest
                      package_path: digest_0.6.39.tar.gz
                      bioconductor: true
                    - package_name: rlang
                      package_path: rlang_1.1.7.tar.gz
                      bioconductor: true
                    - package_name: base64enc
                      package_path: base64enc_0.1-6.tar.gz
                      bioconductor: true
                    - package_name: fastmap
                      package_path: fastmap_1.2.0.tar.gz
                      bioconductor: true
                - package_name: rlang
                  package_path: rlang_1.1.7.tar.gz
                  bioconductor: true
                - package_name: base64enc
                  package_path: base64enc_0.1-6.tar.gz
                  bioconductor: true
                - package_name: digest
                  package_path: digest_0.6.39.tar.gz
                  bioconductor: true
                - package_name: fastmap
                  package_path: fastmap_1.2.0.tar.gz
                  bioconductor: true
                - package_name: rappdirs
                  package_path: rappdirs_0.3.4.tar.gz
                  bioconductor: true
              external_dependencies:
                apt:
                    - name: make
                      dependencies:
                        - name: gcc-14-base
                        - name: libc6
                        - name: libgcc-s1
            - package_name: rappdirs
              package_path: rappdirs_0.3.4.tar.gz
              bioconductor: true
        - package_name: cachem
          package_path: cachem_1.1.0.tar.gz
          bioconductor: true
          dependencies:
            - package_name: rlang
              package_path: rlang_1.1.7.tar.gz
              bioconductor: true
            - package_name: fastmap
              package_path: fastmap_1.2.0.tar.gz
              bioconductor: true
        - package_name: commonmark
          package_path: commonmark_2.0.0.tar.gz
          bioconductor: true
        - package_name: fastmap
          package_path: fastmap_1.2.0.tar.gz
          bioconductor: true
        - package_name: fontawesome
          package_path: fontawesome_0.5.3.tar.gz
          bioconductor: true
          dependencies:
            - package_name: digest
              package_path: digest_0.6.39.tar.gz
              bioconductor: true
            - package_name: fastmap
              package_path: fastmap_1.2.0.tar.gz
              bioconductor: true
            - package_name: htmltools
              package_path: htmltools_0.5.9.tar.gz
              bioconductor: true
              dependencies:
                - package_name: digest
                  package_path: digest_0.6.39.tar.gz
                  bioconductor: true
                - package_name: rlang
                  package_path: rlang_1.1.7.tar.gz
                  bioconductor: true
                - package_name: base64enc
                  package_path: base64enc_0.1-6.tar.gz
                  bioconductor: true
                - package_name: fastmap
                  package_path: fastmap_1.2.0.tar.gz
                  bioconductor: true
            - package_name: rlang
              package_path: rlang_1.1.7.tar.gz
              bioconductor: true
            - package_name: base64enc
              package_path: base64enc_0.1-6.tar.gz
              bioconductor: true
        - package_name: httpuv
          package_path: httpuv_1.6.16.tar.gz
          bioconductor: true
          dependencies:
            - package_name: Rcpp
              package_path: Rcpp_1.1.1.tar.gz
              bioconductor: true
            - package_name: R6
              package_path: R6_2.6.1.tar.gz
              bioconductor: true
            - package_name: rlang
              package_path: rlang_1.1.7.tar.gz
              bioconductor: true
            - package_name: fastmap
              package_path: fastmap_1.2.0.tar.gz
              bioconductor: true
            - package_name: lifecycle
              package_path: lifecycle_1.0.5.tar.gz
              bioconductor: true
              dependencies:
                - package_name: cli
                  package_path: cli_3.6.5.tar.gz
                  bioconductor: true
            - package_name: magrittr
              package_path: magrittr_2.0.4.tar.gz
              bioconductor: true
            - package_name: cli
              package_path: cli_3.6.5.tar.gz
              bioconductor: true
            - package_name: otel
              package_path: otel_0.2.0.tar.gz
              bioconductor: true
            - package_name: promises
              package_path: promises_1.5.0.tar.gz
              bioconductor: true
              dependencies:
                - package_name: magrittr
                  package_path: magrittr_2.0.4.tar.gz
                  bioconductor: true
                - package_name: later
                  package_path: later_1.4.8.tar.gz
                  bioconductor: true
                  dependencies:
                    - package_name: rlang
                      package_path: rlang_1.1.7.tar.gz
                      bioconductor: true
                    - package_name: Rcpp
                      package_path: Rcpp_1.1.1.tar.gz
                      bioconductor: true
                - package_name: R6
                  package_path: R6_2.6.1.tar.gz
                  bioconductor: true
                - package_name: rlang
                  package_path: rlang_1.1.7.tar.gz
                  bioconductor: true
                - package_name: Rcpp
                  package_path: Rcpp_1.1.1.tar.gz
                  bioconductor: true
                - package_name: fastmap
                  package_path: fastmap_1.2.0.tar.gz
                  bioconductor: true
                - package_name: lifecycle
                  package_path: lifecycle_1.0.5.tar.gz
                  bioconductor: true
                  dependencies:
                    - package_name: cli
                      package_path: cli_3.6.5.tar.gz
                      bioconductor: true
                - package_name: cli
                  package_path: cli_3.6.5.tar.gz
                  bioconductor: true
                - package_name: otel
                  package_path: otel_0.2.0.tar.gz
                  bioconductor: true
            - package_name: later
              package_path: later_1.4.8.tar.gz
              bioconductor: true
              dependencies:
                - package_name: Rcpp
                  package_path: Rcpp_1.1.1.tar.gz
                  bioconductor: true
                - package_name: rlang
                  package_path: rlang_1.1.7.tar.gz
                  bioconductor: true
          external_dependencies:
            apt:
                - name: make
                  dependencies:
                    - name: gcc-14-base
                    - name: libc6
                    - name: libgcc-s1
                - name: zlib1g-dev
                  dependencies:
                    - name: libc-dev
                      optional: true
                    - name: gcc-14-base
                    - name: libc6
                    - name: libcrypt1
                    - name: libgcc-s1
                    - name: rpcsvc-proto
                    - name: libcrypt-dev
                    - name: linux-libc-dev
                    - name: libc-dev-bin
                    - name: libc6-dev
                    - name: zlib1g
        - package_name: later
          package_path: later_1.4.8.tar.gz
          bioconductor: true
          dependencies:
            - package_name: rlang
              package_path: rlang_1.1.7.tar.gz
              bioconductor: true
            - package_name: Rcpp
              package_path: Rcpp_1.1.1.tar.gz
              bioconductor: true
        - package_name: otel
          package_path: otel_0.2.0.tar.gz
          bioconductor: true
        - package_name: sourcetools
          package_path: sourcetools_0.1.7-1.tar.gz
          bioconductor: true
        - package_name: xtable
          package_path: xtable_1.8-8.tar.gz
          bioconductor: true
        - package_name: spatstat.data
```
